# No broad decline of breeding monarch butterflies in North America: implications for conservation efforts

**DOI:** 10.1101/2021.08.03.454948

**Authors:** Andrew K. Davis, Michael S. Crossley, Matthew D. Moran, Jeffrey Glassberg, William E. Snyder

**Affiliations:** Odum School of Ecology, University of Georgia, Athens, GA, USA; Department of Entomology, University of Georgia, Athens, GA, USA; Department of Biology and Health Sciences, Hendrix College, Conway, AR, USA; North American Butterfly Association, Morristown, NJ, USA; Rice University, Houston, TX, USA

**Keywords:** monarch butterfly, population trends, decline, conservation efforts

## Abstract

Many insects are in clear decline, with monarch butterflies (*Danaus plexippus*) drawing particular attention as a flagship species. Falling numbers of overwintering monarchs are well documented, but there has been debate regarding population trends of summer breeding populations. Here, we compile a series of long-term monarch monitoring datasets, some which are analyzed here for the first time, that reveal highly variable responses across the migratory geographic range, but no broad net decline in numbers of breeding monarchs. We also did not find evidence that sampling biased towards natural sites was masking declines at disturbed sites. Overall, our results suggest a robust resiliency in summer populations that thus far has allowed recovery from losses during the winter. Thus, monarchs may not require as much breeding habitat restoration as once thought, and focus should be on conserving the fall and spring migration.

## Introduction

Despite considerable variability through time, between sites, and among taxa, it is increasingly clear that some of the world’s insects are in steep decline. This is perhaps best documented among bees and other pollinators, whose loss would have devastating consequences for global ecosystems and the human food supply (Wagner 2020). Beyond pollination, insects are key providers of a full suite of provisioning, regulating, cultural, and supporting ecosystem services. Human degradation of the environment, at a range of scales, is often implicated in falling insect numbers. A key local driver has been heavy herbicide and insecticide applications associated with agricultural intensification (Habel et al. 2019). Urbanization and associated automobile collisions (Baxter-Gilbert et al. 2015, Kantola et al. 2019) and light pollution bring additional challenges (Owens et al. 2020). At global scales, climate change can heighten physiological stress to insects while triggering spatiotemporal misalignment with, or reduced quality of, host plants or other resources (Bale et al. 2002, Jamieson et al. 2012), although even climate change can create variable regions of insect decreases and increases (Koltz et al. 2018, Crossley et al. 2021). Particularly damaging are cases where local and global drivers both are moving in harmful directions, for example when long-distance migrants must move through increasingly hot and dry regions that also are seeing more intense land use (Saunders et al. 2019).

Monarch butterflies (*Danaus plexippus*) in North America have become the public face of insect declines (Gustafsson et al. 2015). Monarchs are iconic insects due to their large size, attractive and distinctive coloration, wide range, host association with horticulturally popular milkweeds (*Asclepias* spp.), and fascinating long-distance seasonal migration to dense overwintering colonies in Mexico. This has led to the prominent use of monarchs as ambassadors to engage the general public in insect conservation, for example, by facilitating the widespread planting of milkweed in home gardens (Thogmartin et al. 2017b). However, some of these same traits that make monarchs so charismatic to humans also subject the butterflies to particular risk. Best documented is habitat loss and changing climate at concentrated overwintering sites, which has apparently led to an ongoing, multi-decadal decline of those colonies (Brower et al. 2012, Thogmartin et al. 2017a, Pelton et al. 2019). A second widely-touted threat is removal of milkweed from agricultural fields within monarch’s core breeding range in the American Midwest, following widespread adoption of glyphosate-tolerant corn and soybean (Stenoien et al. 2018). Thirdly, since migration in the human-dominated world is risky (Wilcove and Wikelski 2008), their particularly long-distance movements could expose monarchs to multiple threats (e.g., deaths from traffic collisions (McKenna et al. 2001, Kantola et al. 2019)). Additionally, agricultural and residential pesticides (Olaya-Arenas and Kaplan 2019) and sensitivity to temperature and precipitation extremes as the climate changes (Lemoine 2015, Saunders et al. 2018) may be adversely affecting monarchs at various stages of their life cycle.

The barrage of dire-sounding threats listed above has led to the widespread belief that monarchs are in trouble throughout their annual cycle, and, has even led to the recent decision by USFWS that federal protection is warranted in the United States (USFWS 2020). Despite this action, evidence is ambiguous whether monarchs are in consistent decline across the annual cycle, with studies variously reporting steady or falling monarch numbers at different places and seasonal milestones (Davis and Dyer 2015, Ries et al. 2015, Espeset et al. 2016, Inamine et al. 2016, Brower et al. 2018, Ethier 2020). Uncertainty about whether breeding populations are increasing, decreasing, or relatively unchanged complicates efforts to target conservation programs to points in the life-cycle where they will be most effective.

Here, we combine analysis across a series of long-term datasets to consider abundance trends for monarch butterflies during the breeding season (i.e., 3-5 generations that occur in Nearctic growing season) (Reppert and de Roode 2018), from the initiation of spring migration through travel back to overwintering sites in the fall. We do not consider abundance trends over the winter, where declines are already well documented (Brower et al. 2012, Thogmartin et al. 2017a, Saunders et al. 2019). The abundance datasets that we considered fall into two broad categories. We first compiled observations of monarch abundance from 15 sampling networks and published papers (Table 1, and see suppl. file for more details). Data collected variously included counts of breeding adults, counts of museum records, summaries of breeding range size, surveys of fall migrants, and even annual estimates of regional abundance from isotope analyses of specimens. Specifically, two datasets tracked monarchs in the spring after leaving overwintering sites, seven tracked monarchs during the summer breeding period at various sites along the migration route, and six tracked monarchs during return migration in the fall.The second category of data that we considered was observations of monarchs during the North American Butterfly Association’s summer citizen-science counts. These data are broad in scope, collectively recording 134,684 monarchs at 341 sites across North America, over time periods of 10-26 years from 1993-2018. From the NABA data, we calculated monarch abundance trends while accounting for annual variation in sampling effort as well as temporal autocorrelation in monarch counts. Our central goals were to (1) search for a general pattern in abundance trends, up or down, across these spatiotemporally diverse datasets, (2) seek evidence that commonly-proposed local (agricultural intensification, road density) or global (temperature or precipitation change) drivers are impacting abundance trends, and (3) compare recent abundance change in monarchs to that seen for other North American butterfly species.

**Table 1.** Summary of datasets used in analyses of monarch abundance. See Suppl. file for more details for each project. Abundance trends for each project depicted in Fig. 1 and 2a.

| ID | Dataset/Source | Measure | Season | Time span | Abundance trend (sd/yr) | p value |
| --- | --- | --- | --- | --- | --- | --- |
| a | (Inamine et al. 2016) | NABA reports from southern US in early spring (# monarchs/hr) | Spring Recolonization | 2005-2014 | -0.247 | 0.014 |
| b | (Brower et al. 2018) | No. monarchs counted at the same pasture in Florida each year | Spring Recolonization | 1994-2017 | -0.093 | 0.019 |
| c | (Wepprich et al. 2019) | Walking transects throughout state of Ohio (# monarchs/hr) | Summer breeding | 1996-2016 | -0.093 | 0.010 |
| d | (Ries et al. 2019) | Proportion of butterfly specimens in U.S. museums that were monarchs | Mostly summer | 1906-2016 | 0.008 | 0.017 |
| e | (Flockhart et al. 2019) | Canadian breeding range (km<sup>2</sup>) - based on sightings from two citizen science programs | Summer breeding | 2000-2015 | 0.108 | 0.049 |
| f | Illinois Butterfly Monitoring project, unpubl. data | Walking transects throughout Illinois (# monarchs/hr) | Summer Breeding | 1993-2019 | -0.034 | 0.280 |
| g | (Crewe et al. 2019) | Monarch abundance index calculated by Crewe et al. | Summer Breeding | 2003-2015 | -0.020 | 0.817 |
| h | (Flockhart et al. 2017) | Proportion of monarchs in Mexico that came from the Midwest US | Summer breeding | 1975-2014 | -0.004 | 0.909 |
| i | Iowa DNR, unpubl. data | Walking transects (random placement) throughout state of Iowa | Summer Breeding | 2007-2019 | -0.009 | 0.934 |
| j | Long Point, ON Monitoring project, unpubl. Data | Average no. monarchs counted per census | Fall migration | 1990-2018 | -0.047 | 0.045 |
| k | Massachusetts Butterfly Club, unpubl. data | No. monarchs per census throughout state of MA | Summer and Fall | 1992-2019 | 0.042 | 0.088 |
| l | Journey North Roost Sightings, unpubl. data | Average roost size per year (# monarchs) | Fall migration | 2003-2020 | -0.031 | 0.459 |
| m | (Badgett and Davis 2015) | Average No. monarchs counted per census | Fall migration | 1996-2014 | -0.017 | 0.507 |
| n | (Ethier 2020) | Average no. monarchs roosting per year | Fall migration | 1984-2018 | -0.011 | 0.532 |
| o | Cape May NJ, unpubl. data | Average no. monarchs/hour seen on driving transect | Fall Migration | 1992-2020 | -0.010 | 0.628 |

## Methods

### Data sources

The abundance datasets that we considered fall into two broad categories. We first compiled observations from 15 different monarch sampling networks in North America using publicly-available information (i.e., extracted from published studies), or where unpublished data had been provided directly to the lead author. These datasets included projects where monarchs had been tracked over many years at local sites, citizen science programs with a state-level or continental-scale range, or from researcher-led investigations, such as counts of museum records, estimates of breeding range size, or estimates of regional abundance from isotopic analyses of monarch specimens. The time spans of data range from 10-96 years between 1901-2020 (Table 1). Full descriptions of each data source are provided in the Supplemental File. Of these 15 datasets, 2 tracked monarchs in the spring after leaving overwintering sites, 7 tracked monarchs during the summer breeding period at various sites along the migration route, and 6 tracked monarchs during return migration in the fall.

The second category of data that we considered was direct counts of monarch adults from the North American Butterfly Association’s summer citizen-science counts (https://www.naba.org/). Butterfly counts are made within a 15-mile (∼24 km) diameter circle, typically in July, and are open to participation from the public. For each count event, the abundances of butterfly species are tallied and the sum of associated party hours (a measure of sampling effort that aggregates the number of hours spent by each observer) is recorded. To minimize bias due to differences among sites in the day of year when butterfly counts were conducted, we limited our analysis to butterfly counts that occurred between June-August. Prior to estimating trends in abundance, we removed sites that had < 5 data points and that spanned < 10 years (Didham et al. 2020). Lastly, we retained for analysis only sites that witnessed at least 30 butterflies in total over the entire time series. The curated dataset recorded a total of 134,684 monarchs from 341 sites across North America, over time periods of 10-26 years from 1993-2018.

### Estimating abundance trends

Monarch abundance trends were estimated using generalized least squares regression (Pinheiro 2020). Models included a first-order autocorrelation term (R parameter “corAR1(form=∼1)”) to account for temporal autocorrelation in residuals. For each abundance time series, we first scaled the independent variable, time in years, to be between zero and the length of the time series by subtracting the start year from each year. This enabled trends to be interpreted in terms of change per year. For the 15 monarch sampling networks, the response variable was the log-transformed measure of monarch abundance (see Table 1 for a description of abundance measures for each dataset). To avoid negative infinity values where the abundance measure was zero, we added half of the minimum non-zero value to each zero value prior to log transformation. Log-transformed values were then z-score-transformed to enable interpretation of trends as change in standard deviations per year. For the Ontario Butterfly Atlas dataset (letter ‘g’ in Figure 1), abundance measures were already a relative index, so values were only z-score-transformed (i.e., no log transformation was done).

**Figure 1.**
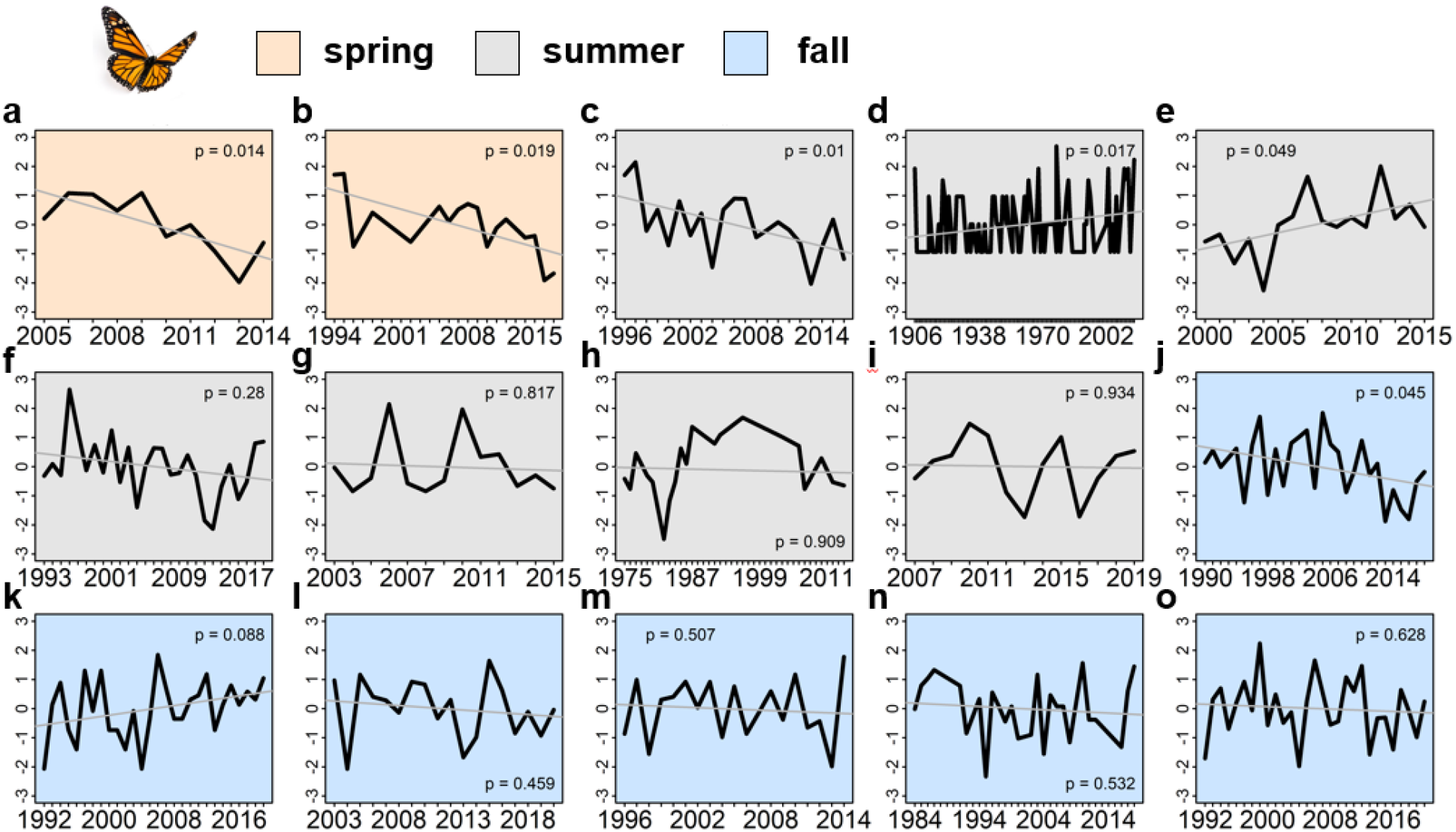
Trends in 15 datasets monitoring adult monarch abundance in North America in the spring, summer, and fall. Trends were estimated using generalized least squares regression of monarch abundance on year while including a first-order autoregressive error structure to account for temporal autocorrelation of counts among years. Monarch abundance was natural log and z-score transformed prior to regression (except for the dataset in panel g, which reported a derived annual index that was only z-score transformed) to put trends on a common standard deviation per year scale. a, North American Butterfly Association volunteer surveys of adults in the spring colonization flyway March-June, 2005-2014; b, Spring surveys of arriving adults and immatures at Cross Creek, FL, 1994-2017; c, Transect surveys of adults at sites throughout Ohio, 1996-2016; d, Proportion of monarchs among butterfly museum specimens in the eastern breeding range, 1906-2013; e, Breeding range size in eastern Canada, 2000-2015, estimated from Journey North and Ebutterfly; f, Pollard walk transects throughout Illinois June-August, 1993-2019; g, Ontario Butterfly Atlas annual adult counts throughout Ontario, Canada 2003-2017; h, Annual proportion of monarchs arriving in Mexico from the Midwest, 1975-2014, based on isotopic signatures; i, Pollard walk transects at randomly chosen sites throughout, Iowa 2006-2019; j, Daily surveys of fall migrating monarchs at two locations along Long Point peninsula in Ontario, Canada, 1995-2019; k, Massachusetts Butterfly Club walks conducted throughout the summer and fall, 1992-2019; l, Average size of roosts reported to Journey North throughout the entire fall migration flyway, 2003-2019; m, Daily censuses of fall migrating monarchs at Peninsula Point, Michigan, 1996-2014; n, Annual census of fall migrating monarchs roosting at Point Pelee, Ontario, Canada, 1984-2018; o, Daily counts of fall migrating monarchs in Cape May, NJ, 1992-2018. Full dataset descriptions are given in Supplementary file.

For the North American Butterfly Association data, we explored two ways of handling annual variation in sampling effort (party hours) in each time series, which could bias abundance trend estimates if sampling effort generally increased or decreased through time. The first approach was to include log-transformed party hours as a weighting term in the model. We used a log transformation to account for the expectation of decreasing returns in numbers of monarchs observed as sampling effort became exceptionally large (rather than linear or exponential increase with each additional hour of sampling effort). The second approach was to divide monarch counts by the associated number of party hours, yielding values of number of monarchs per hour of sampling effort. Because the abundance measure in the North American Butterfly Association data was consistent across sites (number of monarchs), we only log transformed abundance values prior to modeling (no z-score transformation was necessary). The two approaches thus resulted in trends that could be transformed to percent change per year by multiplying the exponent of the trend minus one by 100. Because resulting trends were similar in magnitude and direction (Supplemental Fig. 2), in further analyses we only included the trends in number of monarchs weighted by party hours (modeling approach #1).

### Environmental covariates

Monarch abundance trends were modeled as a function of three land use and four climate covariates. The first two land use covariates concern impacts of agriculture on monarch larval habitat: the amount of cropland and the amount of glyphosate applied to the landscape (Pleasants and Oberhauser 2013). The third land use covariate, road density, captures the potential severity of roadkill among dispersing monarch adults (Kantola et al. 2019, Tracy et al. 2019). Cropland data were obtained from the United States Department of Agriculture – National Statistics Service Cropland Data Layer from 2008-2018 (USDA-NASS 2018), and road density data were obtained from the United States Geological Survey National Map (USGS 2014). Cropland was defined as land in any cultivated crop (e. g., soybean; 107 categories in total). Proportion cropland was calculated as the number of 30×30 m pixels classified as cropland divided by the total number of pixels within each ∼24 km diameter count circle, averaged between 2008-2018. Road density was similarly calculated as the number of 30×30 m pixels that contained a road segment divided by the total number of pixels within each count circle. Glyphosate use data were derived from the United States Geological Survey – National Water-Quality Assessment Project (USGS 2021), which provides county-level summaries of the upper and lower estimates of the amount of >400 pesticides applied to agricultural crops from 1992 to 2017. To obtain a measure of glyphosate use in each count circle, we first calculated the average upper estimate of glyphosate use (lbs) for each county across years from 1993 to 2017. We used the upper estimate because lower use estimates are not available for many counties. Because count circles often overlapped multiple counties, we used areal-weighting (summing across counties the product of glyphosate use and the proportion of the county overlapped by a count circle) to arrive at an estimate of average glyphosate use within each count circle. We acknowledge that glyphosate applied specifically to genetically modified corn and soybean has been a focus of previous work (Pleasants and Oberhauser 2013), but we retained glyphosate use (lbs), agnostic to land cover composition in a county, because we wished to minimize multi-collinearity with cropland and reasoned that glyphosate applications are detrimental to milkweed regardless of landscape composition.

Climate covariates included contemporary measures of mean temperature and cumulative precipitation, as well as long-term changes in each. Climate data were obtained from CRU TS 4.03 (Harris et al. 2014), which provides monthly climate data from 1901-2018. Thus, to obtain a contemporary estimate of mean temperature and cumulative precipitation, we calculated the annual average of monthly values, then took the average of annual values between 1993-2018. To differentiate effects of contemporary climate from historic changes in climate over the study period, we calculated trends in temperature and precipitation between 1901-2018 using autoregressive models fit using Restricted Maximum Likelihood (Ives et al. 2010), first Z-score transforming climate observations and transforming year according to *y - 1901*. These models allowed estimation of linear trends in precipitation and temperature, whose slopes are interpreted as the change in units of standard deviations year, while accounting for temporal autocorrelation.

Spatial operations were all conducted in R 4.0.4 (R Core Team 2021) using functions available in the packages ‘raster’, ‘rgdal’, ‘rgeos’, and ‘sp’. Environmental covariates were visualized using the R package ‘tmap’. Maps depicting land use covariates across sites are provided in Supplemental Fig. 4.

### Environmental correlation analysis

We examined correlations between monarch abundance trends and environmental covariates using generalized linear models fit with *glm* in base R, hypothesizing that variation in land use and climate change could explain site-level differences in abundance trends. Prior to modeling, we excluded sites that did not have complete representation of environmental covariates (several sites in Canada), leaving monarch abundance trends from 305 sites for analysis, and we z-score transformed covariates to put them on a standard deviation scale. We confirmed that there was no spatial autocorrelation in residuals and that variance inflation factors were < 2 using functions available in the ‘DHARMa’ and ‘performance’ R packages (Hartig 2020, Lüdecke 2020). We used Akaike Information Criterion, corrected for small sample sizes (AIC_c_), to identify best-supported combinations of covariates in models using the *dredge* function in the ‘MuMIn’ R package (Barton 2020). Covariate effects were then estimated by model averaging among the AIC_c_-best model plus the 18 models that were within two AIC_c_ (Supplementary Table 4), and estimates were considered significant if their 95% confidence intervals did not overlap zero. Results suggested that long-term precipitation trend had a significant negative effect on monarch abundance trends, and that all other covariates had non-significant effects. Because there was a strong negative correlation between long-term precipitation trend and contemporary temperature, we repeated the model selection procedure after excluding climate change (long-term precipitation and temperature trend) covariates. This time, nine models within two AIC_c_ were included in model averaging (Supplementary Table 5). Again, we found no significant effect of land use or contemporary precipitation covariates, but this time contemporary temperature had a significant positive effect on monarch abundance trends (Supplementary Table 3). We thus present results from model selection including all land use and climate covariates, but interpret the effect of long-term precipitation trends in light of its strong correlation with contemporary temperature.

## Results

We first considered the set of 15 monarch abundance datasets, from different places and points in the seasonal migration, originally compiled by Davis (2020) (‘Methods’). While trends broadly varied in direction and magnitude (Fig. 1), they were strongly centered near zero with a slim majority (8/15) being statistically indistinguishable from no net change (Fig. 2a; Table 1). While just two spring time series were available, it is notable that both exhibited significant negative trends (−21.9%/year, p = 0.019; -8.9%/year, p = 0.014). One summer trend was significantly negative (−8.8%/year, p = 0.010) while two were significantly positive (+0.8%/year, p = 0.017; +11.4%/year, p = 0.049), with the remaining four exhibiting no significant difference from zero (Table 1). Fall trends were likewise split, with one significantly negative (−4.6%/year, p = 0.045), one marginally significantly positive (+4.2%/year, p = 0.088), and the other four indistinguishable from zero (Table 1). Overall, these collective data indicate that numbers of spring migrants are falling in concert with declining numbers of monarchs that successfully overwinter, but then, generally stable or increasing summer and fall numbers suggest that monarch numbers routinely rebound during the successive summer generations.

**Figure 2.**
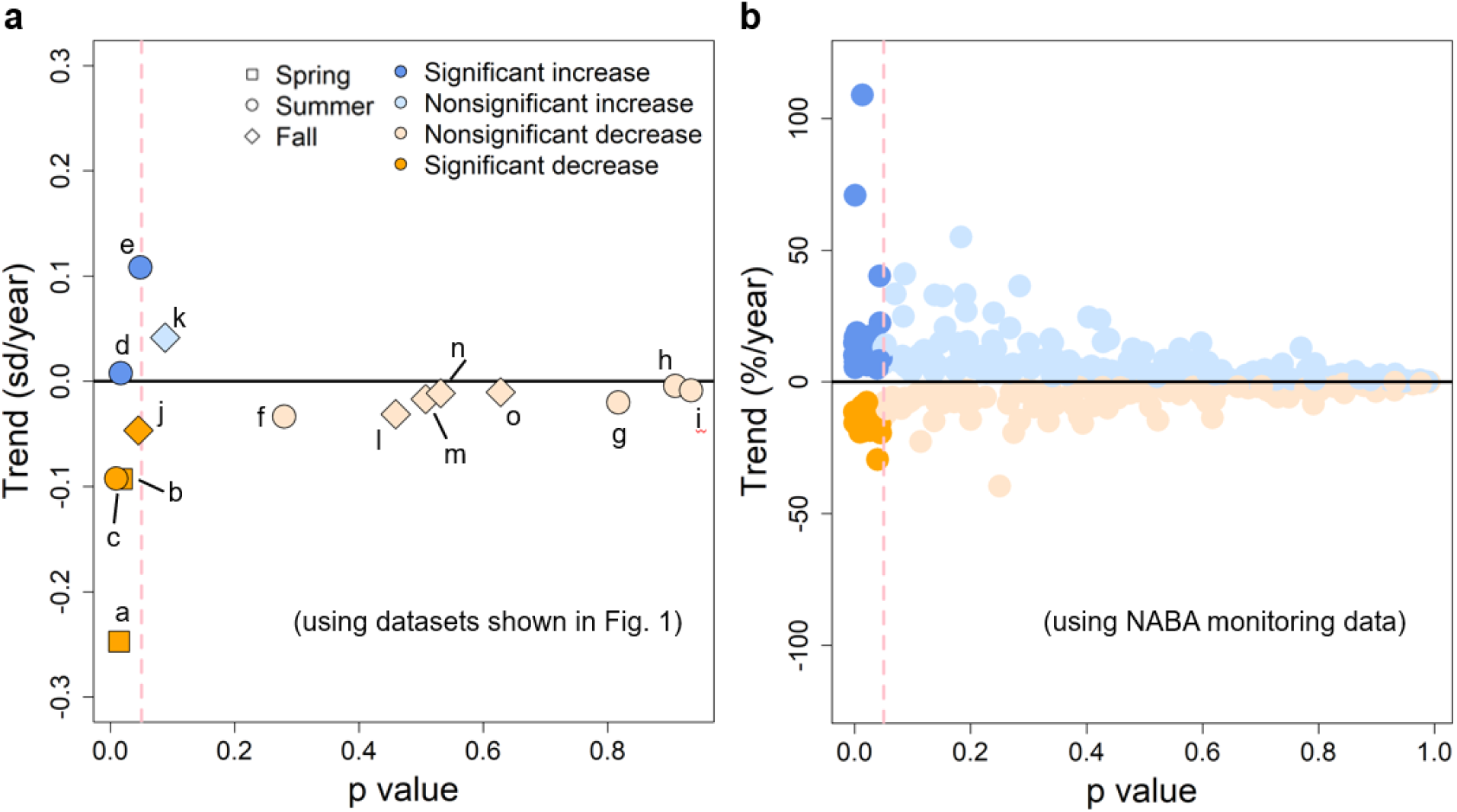
Summary of monarch abundance trends throughout the North American breeding range. a, Trends in selected datasets monitoring monarch abundance in North America in the spring, summer, and fall. Trends were estimated using generalized least squares regression of monarch abundance on year while including a first-order autoregressive error structure to account for temporal autocorrelation of counts among years. Monarch abundance was natural log and z-score transformed prior to regression (except for dataset “g”, which reported a derived annual index that was only z-score transformed) to put trends on a common standard deviation per year scale. Symbol letters correspond to the panels in Fig. 1. b, Trends among sites included in the North American Butterfly Association dataset. Trends were estimated using generalized least squares regression of natural log-transformed monarch abundance on year while including a first-order autoregressive error structure to account for temporal autocorrelation of counts among years, and weighting observations by the number of party hours to account for variation in sampling effort over time. Trends were considered significantly different from zero when p value < 0.05.

The North American Butterfly Association counts were a particularly abundant data source, but only for roughly mid-summer (Crossley et al. 2021). Notably, the distribution of abundance trends at these spatiotemporally well-dispersed sites were broadly consistent with those from the other sampling networks (compare Fig. 2a-b). Abundance trends were variously positive, negative, or unchanged at different sites, clustering close to zero (Fig. 2b). Consistent with this, 88% (350/400) were statistically indistinguishable from zero, with 8% significantly increasing and 4% significantly declining (Supplementary Table 2). So, on the whole, trends were weakly positive, but not substantially so. Overall, we again found that summer monarch abundance trends were broadly unchanged through time, with significant increases or declines at particular sites largely offsetting each other.

We are aware that a prior argument against using evidence from non-random, citizen science monitoring programs such as NABA, is that they provide misleading trends, because people tend to monitor where they know butterflies would be present (Pleasants et al. 2015, Pleasants et al. 2017, Kinkead et al. 2019). It has also been argued that losses of milkweed in the agricultural fields of the Midwest has “shifted” adults away from these areas and into the more pristine monitoring sites (Pleasants et al. 2015, Pleasants et al. 2017), which could artificially inflate abundance at these sites. The NABA data we used does include sites in the Midwest, though it should not be overlooked how large the monitoring areas are (458km^2^ each), and the fact that they range in agricultural use, from those almost entirely surrounded by annual crops to those almost entirely surrounded by natural habitats (Fig. 3a). This provides some ability to examine any effect of surrounding farmlands on monarch abundance trends (Crossley et al. 2021). A simple mapping of increasing, decreasing, or relatively unchanged monarch abundance trends revealed no obvious regional clumping (Fig. 3b), although sampling points were relatively dense in the northeastern and upper-central US with sparser coverage elsewhere.

**Figure 3.**
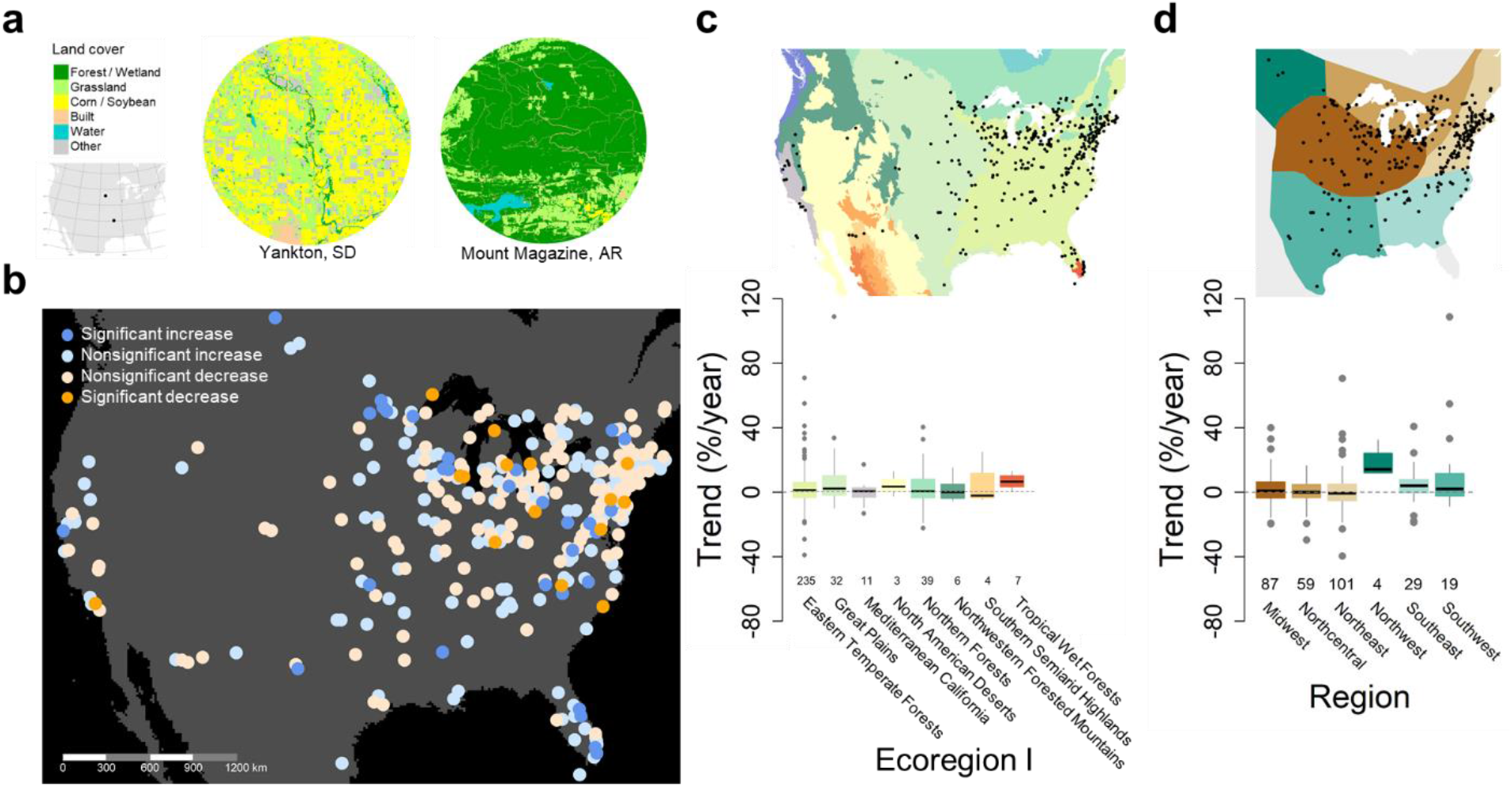
Spatial variation in monarch trends across the summer breeding range. **a**, North American Butterfly Association butterfly count circles are ∼15 mi in diameter and span a range of agriculturally dominated and seminatural habitats. **b**, Increases and decreases in monarch abundance throughout the summer breeding range, mapping the trends depicted in Fig. 1b. **c**, Monarch abundance trends grouped by EPA level I Ecoregion. **d**, Monarch abundance trends grouped by monarch breeding regions *sensu* Flockhart et al. (2017). Boxplots depict trends among monarchs as medians (thick line), 25th and 75th percentiles (box edges), 95th percentiles (whiskers) and outliers (circles).

We are also aware that conversion to glyphosate-tolerant corn and soybean has been particularly complete in central North America, a heavily farmed region overall, which has led to near elimination of milkweeds from fields (Hartzler 2010). However, breeding monarch abundance trends were variable but clustered near zero when considering either the Great Plains Ecoregion compared to other Ecoregions (Fig. 3c), or the “Midwest” monarch region compared to other parts of North America as defined by Flockhart et al. (2017) (Fig. 3d). Likewise, the amount of cropland and glyphosate use in surrounding landscapes had negative but nonsignificant effects on monarch summer abundance trends (Supplementary Table 3). We similarly found a negative, nonsignificant effect of road density, another important source of monarch mortality (Supplementary Table 3). In the end, there was little evidence that these trends were misleading because of changes in agricultural practices or shifting adult distribution. Also important is the consistency we observed in trends from the NABA data with those from fall migration monitoring, where habitat biases are less important.

## Discussion

From the collective evidence compiled here, which included data from spring, summer and fall monitoring, and from a variety of monitoring schemes and projects, we conclude that the *breeding* population of monarchs in North America is not currently in decline. Our analysis considered multiple sources of data, different time frames, and took into account sampling effort, climatic variables, and anthropogenic factors like glyphosate use. We are aware that this conclusion is in stark contrast to studies that focus on winter colony size as measures of population abundance, where there are clearly long-term declines (Brower et al. 2012, Semmens et al. 2016, Oberhauser et al. 2017, Thogmartin et al. 2017a, Zylstra 2021). Our analyses nonetheless suggest that these winter declines may not reflect the population status at other stages, casting doubt on the otherwise sensible assumption that winter declines necessarily lead to concomitant decline in summer monarchs (e.g., USFWS 2020). If there currently are sufficient resources (habitat, hostplants, nectar, etc.) to sustain relatively robust numbers of breeding monarchs, it may be time to reconsider the idea that monarchs are in dire trouble throughout the year.

Of course, there are several caveats to our meta-data and analyses that must be acknowledged. First, there are other long-term monarch datasets that were not available to us and so could not be included. Nonetheless, the broad diversity of positive and negative summer and fall abundance trends collected here, across so many different sampling networks, suggest a limited ability for additional data points to greatly alter the overall pattern. A second concern is that most of the data assembled here were collected by citizen scientists (Table 1). Indeed, the number of party hours spent monitoring butterflies in the North American Butterfly Association dataset increased on average by 1.2% (±0.3%) per year between 1993-2017 (Supplemental Fig. 1). However, our abundance trend models accounted for annual variation in sampling effort, and we note that monarch abundance trends were relatively robust to whether we standardized monarch counts by associated party hours or included party hours as a weighting term in models (Supplemental Fig. 2). Importantly, increasing trends in sampling effort were not clustered around sites dominated by cropland or forest, suggesting that changes in sampling effort have neither masked declines nor exaggerated abundance increases (Supplemental Fig. 3).

Overall, our findings suggest monarch populations have a remarkable ability to recover, on average, from declines at overwintering colonies. Of course, the total loss of overwintering monarchs would make it impossible for any summer rebound to be ignited, and there almost certainly is some inflection point well before total winter extinction where spring migrants would be too few to reliably spark a summer resurgence. However, our results at least indicate the current trends observed in winter are not affecting the breeding population now. Our analysis indicating breeding monarchs are not in decline also argues that the loss of agricultural milkweed in the U.S. Midwest has not doomed summer monarchs, as was once thought (Agrawal and Inamine 2018). Likewise, we could not find evidence that growing road construction has substantially harmed breeding monarchs. Of course, the negative effect of roads may be more pronounced during the southward fall migration (McKenna et al. 2001, Alvarez et al. 2019, Kantola et al. 2019).

Recent analyses indicate that changing climate is driving increases and decreases in overall butterfly numbers across North America (Crossley et al. 2021, Forister et al. 2021), and, there is evidence that climate in North America is indirectly impacting abundances of overwintering monarchs (Zylstra 2021). In line with this, we recovered a significant, weakly negative correlation between increasing long-term precipitation and fewer breeding monarchs (Supplementary Table 2). Sites where precipitation has been increasing also have experienced relatively cool recent temperatures (see ‘Methods’; Supplementary Table 3), suggesting that falling monarch numbers at these sites might reflect slower development, which lengthens exposure of larvae to natural enemies and other risks, at relatively cool and wet locations.

Limited sampling of the arid Western US (Fig. 2b), where broad butterfly declines at increasingly hot and dry sites are clearly apparent, might explain why we did not find the same pattern for monarchs (Crossley et al. 2021, Forister et al. 2021). The eastern U.S. and Canada, the area corresponding to the major monarch summer breeding ground for the Mexican long-distance migration subpopulation, has generally seen increases in precipitation and only modest increases in summer temperature (National Oceanic and Atmospheric Administration 2021), conditions that have apparently been providing favorable conditions for many butterfly species (Crossley et al. 2021). However, Texas and the northern portions of Mexico, a vital corridor region, have seen recent pronounced increases in temperatures (Cuervo-Robayo et al. 2020) which could be affecting survivorship during the arduous migration. In total, this evidence suggests, alongside the ongoing declines at winter colonies, that monarchs must be experiencing increasingly higher levels of mortality during their fall migration. Contrasting evidence of no change in the number of tagged monarchs returning to Mexico in the fall suggests otherwise (Taylor et al. 2020), but that finding remains contested due to difficulties in accounting for changing tagging effort through time (Fordyce et al. 2020). Our results argue that following the winter period, monarchs experience high population growth, perhaps facilitated by reduced intraspecific competition among larvae and generally favorable environmental conditions during breeding. Therefore, conservation attention along the migration routes may be more imperative for the monarch’s survival compared to efforts on the breeding grounds.

When placed in context with other reports, our results underscore the difficulty in pinpointing long-term trends of a species with such a broad geographic distribution, different life stages, and using different monitoring schemes. For example, winter colony sizes have declined in both the eastern and western subpopulations (Thogmartin et al. 2017a, Pelton et al. 2019), yet those trends do not reflect the status of the summer cohort. Even the location of monitoring appears to matter; analyses of long-term counts of summer monarchs in one section of California revealed declines (Espeset et al. 2016), yet our summary of another dataset from the same region revealed no obvious trend (see California region in Fig. 3B). Recent research suggests that the California monarchs are undergoing a rapid shift in life history success with resident populations becoming more common while migratory populations continue to decline (Crone and Schultz 2021, James 2021), which also complicates attempts to track abundance of the traditionally migratory western subpopulation. The manner in which data are examined also is relevant; recent analyses of citizen science data, including some used here (but using different analytical approaches), appeared to show modest declines in abundance of breeding monarchs (Zylstra 2021). Meanwhile, our approach considered these same citizen science data, plus evidence from researcher-derived investigations, and including data from spring, summer and fall abundance metrics. Thus, while the monarch is fortunate to have such a plethora of data and monitoring programs, it is clear that to obtain the clearest picture of its status, one must consider as much evidence as possible, and from multiple life cycle phases, as was done here.

The conservation of insects has received far less attention than most other taxa, despite the ubiquity of insects in terrestrial ecosystems. Undoubtedly, citizen-science efforts targeting the charismatic monarch have exposed many non-scientists in North America to the importance of insects and the value of their conservation. Given our results, we suggest that there could be considerable ecological gain from turning citizen scientists’ attention to the many butterfly species who do appear to be experiencing major summer declines across North America. For example, the summer butterfly count data suggest that *Lycaeides melissa* is declining across much of its broad range (Fig. 4), and even the well-known west coast painted lady, *Vanessa annabella*, appears to be faring worse than the monarch (see Supplemental Table 6). In fact, of the 456 butterfly species tracked by NABA, we determined there are 320 species with trends less positive than monarch butterflies (Supplemental Table 6).

**Figure 4.**
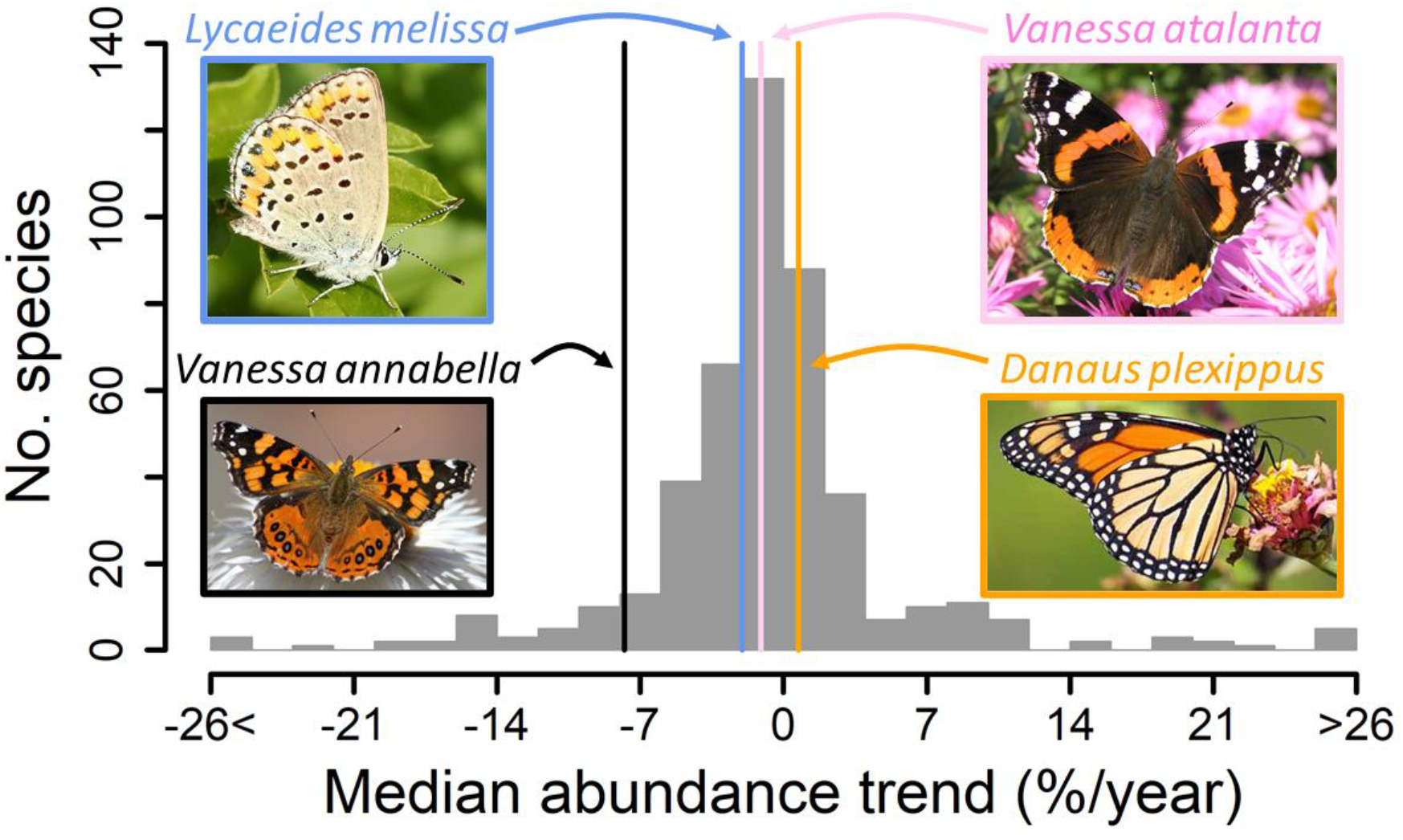
Monarch abundance trend compared to other North American butterflies. Histogram depicts median abundance trends (%/year) of >450 species monitored by the North American Butterfly Association. Trend for *Danaus plexippus* (+0.7%/year) is highlighted and compared to three other well-known species, *Lycaeides melissa* (−2.0%/year), *Vanessa annabella* (−7.8%/year) and *Vanessa atalanta* (−1.1%/year). Trends based on sites where butterflies were recorded at least five times over a span of ten years. See Crossley et al. (2021) for details on trend estimation. All butterfly species trends are available in Supplemental Table 6.

More broadly, these results are consistent with other recent analyses of large-scale insect data, and which have also appeared to counter long-standing notions of widespread declines. For example, a warming climate in Europe is shifting some moth ranges northward, with species unable to do so declining, but leading to a net range increase overall (Fox 2021). Similarly, recent drops in U.K. moths seem modest relative to increases seen over prior decades (Macgregor et al. 2019), leading to no net change over time. In North America, close examination of long-term insect counts revealed declines in some taxa, but increases in others (Crossley et al. 2020). The same is true with butterflies, where species declines in western North America may be at least partially offset by abundance increases elsewhere on the continent (Crossley et al. 2021), again, leading to no net change. These recent studies highlight pitfalls in extrapolating from declines seen at finer spatial, temporal, or taxonomic scales to a larger, all-encompassing narrative of an “insect apocalypse”. The situation with monarchs appears to be similar, where there is general acceptance that the North American population is in trouble, yet a close look at its abundance metrics reveals a more complex story. Our analyses show that for monarchs, for now, summer abundance increases appear sufficient to buffer winter declines. It will be increasingly important to understand complex interactions among species traits and mechanistic drivers, in order to understand and successfully predict how an ever-more-rapidly changing environment will impact the future persistence of monarchs and other insects.

## Acknowledgements

We acknowledge funding from USDA-NIFA-OREI 2015-51300-24155 and USDA-NIFA-SCRI 2015-51181-24292 to W.E.S. We thank Anurag Agrawal for helpful feedback on an early version of the manuscript. This paper was developed following many lengthy and stimulating discussions with colleagues who study monarch butterflies, and who we believe all have the same goal: to understand and identify potential threats to the monarch population in North America.

## Author contributions

A.K.D., M.S.C., M.D.M., and W.E.S. conceived of the idea for the paper, and M.S.C. conducted analyses; A.K.D. and M.S.C. led data collection and curation; A.K.D., M.S.C., and W.E.S. and primarily wrote the paper, although all authors contributed to the final manuscript.

## Data availability

The data supporting the findings of this study (abundance trends and environmental covariates) are available at GitHub (https://github.com/mcrossley3/MonarchTrends). Monarch count data from published studies in Table 1 are viewable within the articles. Data from unpubl. sources in Table 1, and from NABA should be requested from the project coordinators.

## Code availability

The R code used to analyze data are available at GitHub (https://github.com/mcrossley3/MonarchTrends).

