## Supplemental file for "No broad decline of breeding monarch butterflies in North America: implications for conservation efforts"

**This PDF file includes:**

Summaries of datasets used in analyses.

Supplementary Figures 1-4

Supplementary Tables 1-6

**Summary of datasets used**

The following are descriptions of where the data on monarchs originated, and how they were originally collected. Links to the programs or publications are provided for more information. The letter for each project corresponds to its row letter in Table 1 of the main paper, and of the trend points presented in Fig. 2a.

a. North American Butterfly Association

This is a continent-wide citizen science organization that has been operational since 1993, and it is composed of butterfly enthusiasts and naturalists. It asks its volunteers to conduct annual surveys of butterflies in the middle of the summer breeding season. Surveyors are meant to keep track of the time spent searching, and the number of observers are taken into consideration. The observers count all butterflies seen within a circle with a ~24 km (15 mile) diameter (458 square km), during one day, typically on the 4th of July (although now observers can record more than one survey day). There are over 450 sites monitored in this program.

More information on this program is available here - <https://www.naba.org/>

We used these data in two ways. In one, we used the index of early spring monarch abundance from the southern U.S., which had been calculated and used previously by Inamine et al (2016). This index spanned the years 2005 and 2014, and is a useful assessment of the number of returning adults from Mexico. Second, we used the complete NABA dataset, provided to us by JG, for analyses of continental monarch trends, as well as comparison with trends of other butterflies (see Fig. 4 in main paper).

b. Spring surveys at Cross Creek Florida

Beginning in 1981 and onward, the late Lincoln Brower and colleagues have been monitoring the number of eggs and larvae found on milkweeds during the spring in a cattle pasture and surrounding areas in Florida. The 9 ha pasture is at “Cross Creek” in Alachua Co., FL (N 29°31.75′, W 82° 11.86′), and it contains more than 1000 *Ascepias humistrata* (sandhill milkweed) plants. Surveys are conducted every 3-4 days throughout the spring recolonization (mid-March to mid-May). The data from this project were recently compiled and presented (Brower et al. 2018). In that paper, Figure 10 reports the annual average number of eggs/stem in the surveys. We extracted these annual numbers from the figure and used these data here.

c. Ohio Butterfly Monitoring Program

The Ohio Lepidopterists’ Society has been conducting weekly walking censuses of butterflies throughout the state since 1996. Volunteers walk the same transects each week and record all butterflies. For more information on this program, see <http://www.ohiolepidopterists.org/index.html>

These data have been recently examined for trends in abundance of a wide variety of butterfly species (Wepprich et al. 2019), including monarchs. T. Wepprich provided the lead author with data for monarchs.

d. Museum records of monarchs

A recent study collated records of monarch specimens in museums in North America, in an attempt to track changes in their abundance over the last century (Boyle et al. 2019). The study was highly-criticized (Ries et al. 2019, Wepprich 2019) for the approach used to account for temporal variation in collection effort, which was to present the annual number of monarch specimens out of the total number of Lepidopteran specimens. The reason is that collection of night-flying moths is vastly different than collecting butterflies (UV lights versus hand nets), so the two should not be lumped together to control for effort. A more logical approach would be to calculate the annual proportion of butterfly specimens that are monarchs. Ries et al. (2019) did so in their rebuttal of the original paper, and presented a graph showing the corrected estimates of monarch abundance over time. These authors also demonstrated that this approach is also biased by outlier years, and so removal of these years is prudent. We extracted these data from this graph for use here. Thus, the dataset here represents the annual proportion of specimens that are monarchs. This dataset extends from 1906 to 2016.

e. Annual Estimates of Breeding Range size in Canada

A recent study (Flockhart et al. 2019) compiled citizen science sightings of monarchs in eastern and western Canada to map out the distribution of the breeding range in that country. They used sightings from both Journey North, and eButterfly, and across 16 years (2000-2015). They estimated the breeding range size per year using spatial modelling, while taking into account bias from human density. They presented results on the variation in breeding size across years, populations, and in relation to weather variables.

We extracted the relevant information on the annual size of the eastern population breeding range from Figure 1B, which showed the change in breeding range size over time.

The study is open access and available here - <https://www.facetsjournal.com/doi/pdf/10.1139/facets-2018-0011>

f. Illinois Butterfly Monitoring Program

This citizen science program has been operational since the early 1990s. It has its volunteers survey specific transects (walking) throughout the state of Illinois, reporting all butterflies seen. The sites are monitored on a weekly basis each summer (at least 6 visits per season). The standardized nature of the surveys, and the wide variety of habitats covered, make these data invaluable for tracking changes in abundance of butterflies. Moreover, given the location of the state within the core breeding range of the eastern population, and within the agricultural Midwest, the counts of monarchs seen are especially important. For more information about this program, see <https://bfly.org/>

The lead author was kindly supplied data on the annual tally of monarchs across the entire state by the program coordinator, Doug Taron. These data span 26 years (1993-2019).

g. Ontario Butterfly Atlas

Run by the Toronto Entomologists’ Association, the Ontario Butterfly Atlas (<https://www.ontarioinsects.org/atlas/>) compiles sightings and records of butterflies in grid squares across the province, to map out their individual breeding distributions. The sightings come from a variety of sources, including museum collections, eButterfly, Butterflies and Moths of North America (BAMONA), and iNaturalist. The data for monarchs (from 2003-2017) was examined in a recent study (Crewe et al. 2019) that used these records to compile an index of annual abundance. This index took into account the varying level of search effort and inherent latitudinal variation. The annual index of abundance was presented in a figure in the paper (Fig. 2); we extracted these indices for use here.

h. Estimates of Midwestern monarch production

In 2017, a study was published that examined archived collections of monarch specimens from the Mexican overwintering colonies (Flockhart et al. 2017) to estimate where in North America they originated from. The authors used isotopic analysis of wing tissues to determine natal origins of specimens dating back to the mid-1970s, when the winter colonies were first discovered. The authors delineated the population’s entire breeding range into 6 distinct “breeding regions”, of which one was the central Midwest (Fig. 1 in the paper). This region is of particular importance because of its known history as being the core of the breeding range (Wassenaar and Hobson 1998), and the area that is most affected by agricultural practices.

By compiling the total number of specimens that originated from the Midwest each year (out of the total examined that year), this approach allowed for annual estimates of “monarch production” from the Midwest vs other regions. In other words, how many monarchs at the overwintering colonies came from the Midwest. Specifically, the researchers presented a graph showing the temporal changes in regional production (Fig. 1). From this graph we extracted the information on the proportion of monarchs that came from the Midwest each year, over the 38-year period (1976-2014). While these numbers represent percentages, not counts, the long-term time series, plus the relevance of the Midwest to the monarch story, means they are useful nonetheless.

i. Iowa MSIM Program

The Iowa Department of Natural Resources has been conducting standardized wildlife surveys (including monarchs and other butterflies) each summer throughout the state since 2006, which they call the Multiple Species Inventory Monitoring (MSIM) Program. These are conducted on wildlife management areas as well as in private lands, and they entail personnel visiting the sites on a weekly basis and recording all butterflies seen along a transect while walking at a uniform pace (Kinkead et al. 2019). In a recent analysis of these data, Kinkead et al (2019) stated there were 420 sites throughout the state. However, not all of the sites were randomly-chosen and one of the take-aways from that paper was that this can affect the interpretation of results from a long-term study.

The lead author of the study (Kinkead) provided Davis with an updated dataset from this program (2007-2019), which includes only data from sites that were randomly-chosen (n=333 sites). The units of measurement in these data are the number of monarchs seen per transect.

j. Long Point Migration Census

The Long Point Bird Observatory in Ontario Canada employs seasonal workers to track migrating bird abundance at their peninsula site on the north shore of Lake Erie, and starting in 1995 they also began tracking daily numbers of monarchs each fall. The standardized counts consist of a 1-h afternoon walking census conducted between 1300–1700 hours along a delineated path, during which a single surveyor counts the number of monarchs seen foraging or passing through the count area (Crewe and McCracken 2015). Two sites along the peninsula are monitored each year, called “Breakwater” and the “Tip”.

These monarch census data are available for download (with permission) from the NatureCounts web portal (<https://www.birdscanada.org/birdmon/default/main.jsp>), run by Bird Studies Canada. The lead author downloaded the most up-to-date census data (1995-2019). For each year we compiled the average census count for both sites, and then derived the overall average of these to create a time series consisting of a single abundance index per year (since using data from both sites is redundant).

k. Massachusetts Butterfly Club

This is a group of experienced naturalists that conduct semi-regular field surveys of all butterflies at specific sites in the state of Massachusetts. It was formed in early 1990s “to promote the continued appreciation and documentation of the state’s butterflies.” It has a records compiler, and publishes a semi-annual journal, Massachusetts Butterflies. The journal publishes sightings and season summaries. From the end of the 1990’s through 2010, the club consisted of a core of about 150 members, most of whom are seasoned butterfly observers.

Since surveys were not regular, and with varying personnel, the number of monarchs seen per year is derived from calculating the total number of monarchs seen each year, per the number of field reports each year. All data was provided to the lead author directly from the records compiler, Mark Fairbrother.

More information about this group is provided here - <https://www.naba.org/chapters/nabambc/>

l. Journey North roost sightings

Each fall since 2003 the Journey North citizen science program asks its participants to report sightings of migratory roosts. These sightings can be from anywhere in the flyway - typically they are in a tree on a homeowner’s property, and they can range from a few to many thousands of monarchs. If possible, the volunteer is asked to estimate the number of monarchs in the roost(s). These roost estimates have been systematically reported since 2005 on the program web portal (<https://journeynorth.org/sightings/>). Not all roost sightings have counts.

The lead author downloaded these sightings for the purposes of this analysis. There were 3557 sightings across the 17 years (2003-2019). We removed all sightings from western states, and also from Florida, and we removed any records of clusters forming at or near the overwintering sites (which are sometimes reported as migratory roosts). Then, we determined the average roost size for each year, for use in analyses here.

m. Peninsula Point Migration Project

This monitoring project began in 1996 and ended in 2014 (18yrs). It was a daily census of migrating and roosting monarchs at the Stonington, Michigan lighthouse. It was conducted nearly entirely by one person – CJ Meitner and later by Gina Badgett. The survey was a walking census along a trail that traversed a variety of habitats. The location of the peninsula on the north shore of Lake Michigan made it an ideal site for assessing the relative size of the migratory cohort in this region; its northern location meant that the census was counting the beginning of the migratory phase.

The data collected from this program had been examined in three prior studies (Meitner et al. 2004, Davis 2012, Badgett and Davis 2015). From these data we used the average census count per year for analyses in the current paper.

n. Point Pelee Migration Roost Counts

Point Pelee National Park is the most southern point of mainland Canada situated on the northern shore of Lake Erie (41° 55' N, 82° 30' W) and is known as an important fall roosting location for monarchs. Since 1984, staff and volunteers at Point Pelee National Park have been conducting daily counts of roosting monarchs from September till the end of October. Prior to 2017 an evening walk was conducted by park staff just prior to sunset around a designated area at the tip of the peninsula. During this walk, all roosting monarch were counted by a single observer. The length of the survey route and duration of the walk was not standardized, nor was this information recorded. Thus, the survey is considered consistent, yet unstandardized. However, given that roosting monarchs remain in place throughout the night and into the morning, standardization is less important.

Information on the park is given here - <https://www.pc.gc.ca/en/pn-np/on/pelee>

The data from the roost counts have recently been analyzed and reported on in a recent publication (Ethier 2020). We extracted the annual index of monarch abundance (average roost size) from a figure (Fig. 2C) in that paper.

o. Cape May Migration Monitoring Project

Starting in the early 1990s, a census of migrating monarchs has been conducted by volunteers each day during the fall at Cape May, New Jersey. The census is made from a vehicle, driving at low speed, and traversing a variety of habitats and back roads in the town. The census route is the same in each case. Only one observer counts, recording all monarchs seen (and without stopping). These census data have been examined previously (Walton and Brower 1996, Walton et al. 2005, Gibbs et al. 2006, Davis 2012) and serve as a useful index of the annual size of the migratory cohort on the Atlantic coast.

The annual census totals (average number of monarchs seen/hr) are public on the program website - <http://www.monarchmonitoringproject.com/mmptwo.html>.

Here, we used the average census count per year for analysis, and using the most recent data available (1992-2018).

**Supplementary Figures**

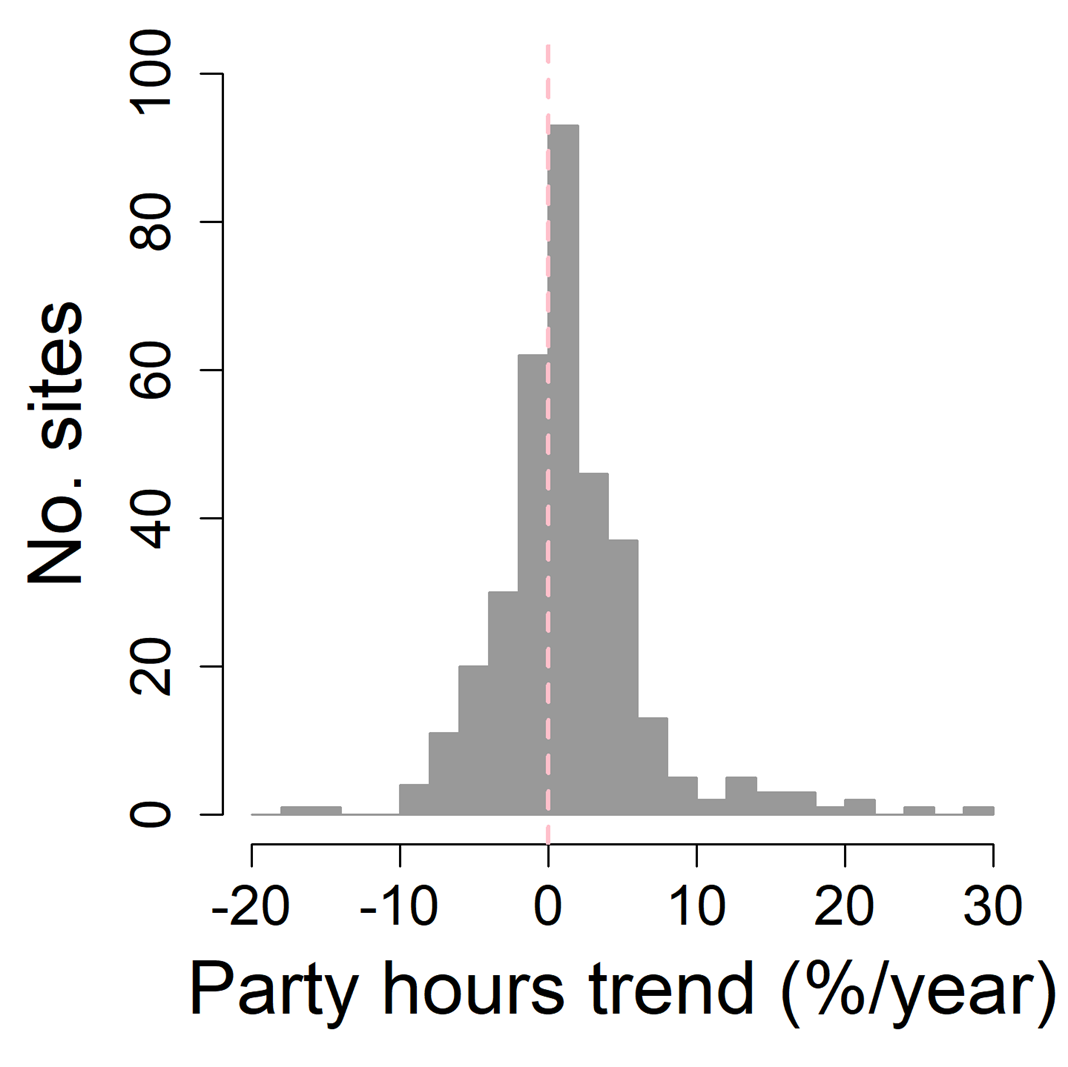

**Supplemental Figure 1.** **Trends in sampling effort among North American Butterfly Association (NABA) count circles.** Histogram depicts trends (%/year) in party hours associated with butterfly counts at each NABA count circle.

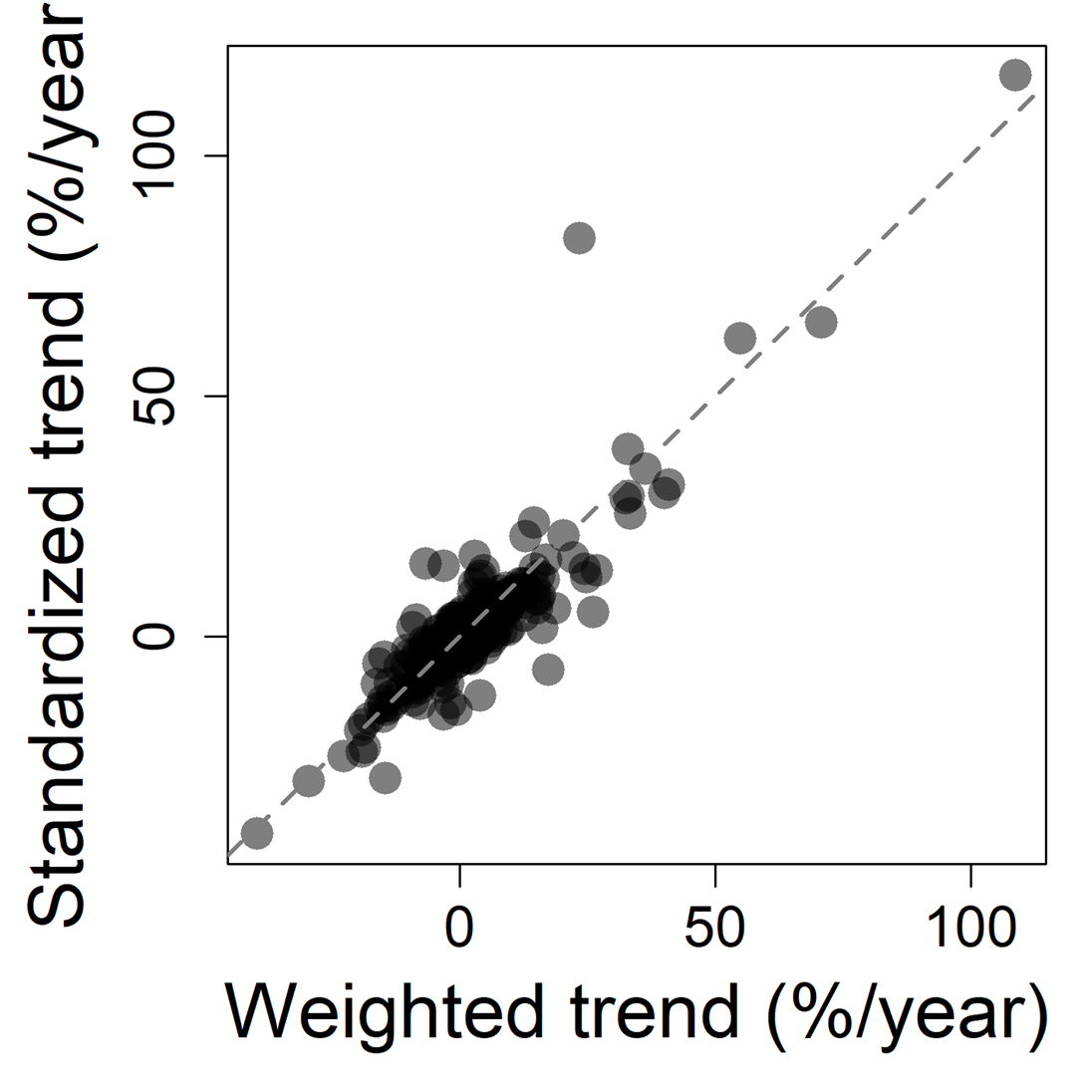

**Supplemental Figure 2.** **Robustness of monarch abundance trends to method of sampling effort standardization.** Scatterplot depicts the relationship between monarch abundance trends (%/year) when log(party hours) were included as a weighting term in generalized least squares regression models (weighted trend), and monarch abundance trends where calculated after dividing monarch counts at each site and year by the associated party hours (standardized trend).

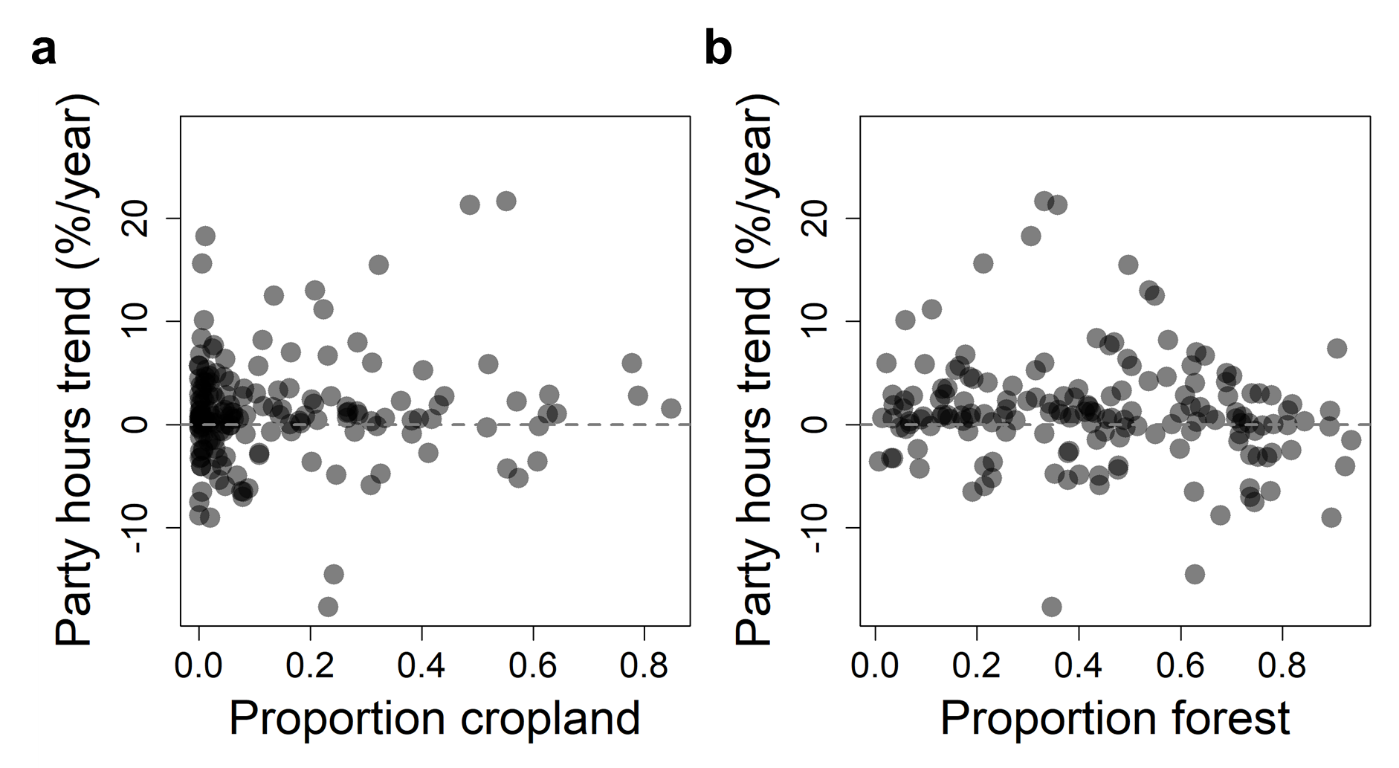

**Supplemental Figure 3.** **Lack of clustering of trends in sampling effort with land cover.** a, Scatterplot depicts the relationship between trend (%/year) in party hours and proportion cropland within ~24 km diameter North American Butterfly Association (NABA) count circles. b, Scatterplot depicts the relationship between trend in party hours and proportion forest and wetland within NABA count circles.

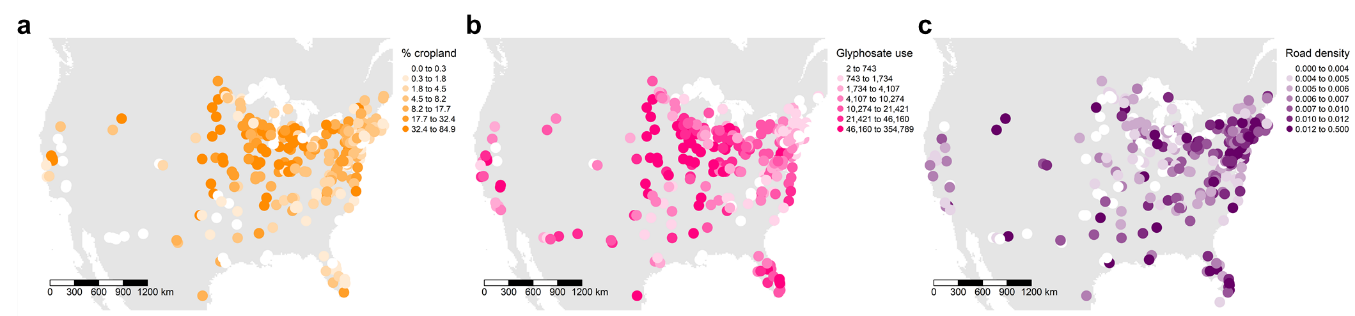

**Supplemental Figure 4. Land use covariates.** a, Average percent cropland between 2008-2018. b, Average glyphosate use (lbs) between 1993-2017. c, Road density.

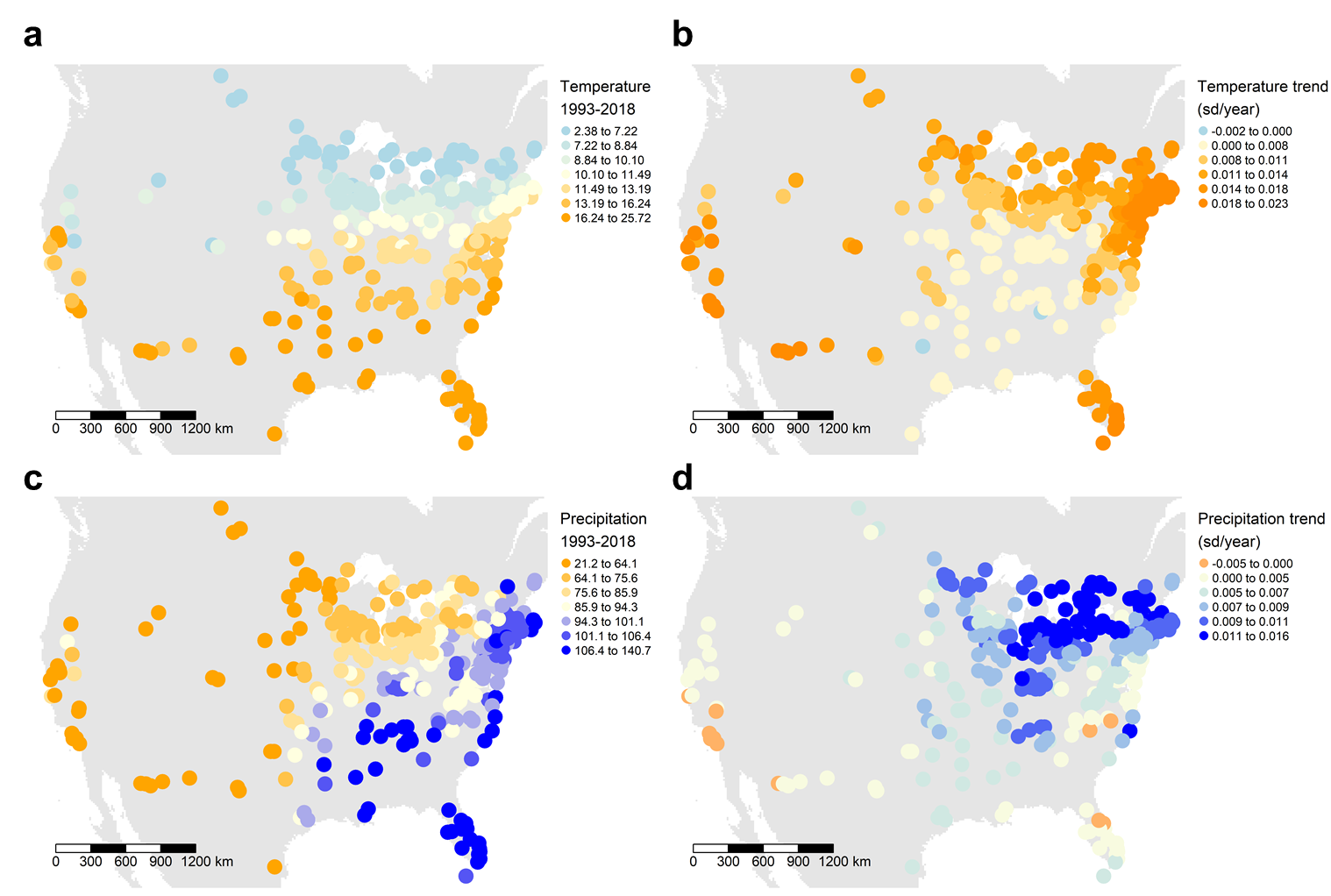

**Supplemental Figure 5.** **Climate covariates.** **a**, Average annual temperature (°C) between 1993-2018. **b**, Trend (sd/year) in average annual temperature between 1901-2018. **c**, Average cumulative precipitation (mm) between 1993-2018. **d**, Trend (sd/year) in average cumulative precipitation between 1901-2018.

**Supplementary Tables**

**Supplementary Table 1**. Summary of abundance trends of *Danaus plexippus* estimated from summer breeding grounds monitored by the North American Butterfly Association between 1993-2018.

| Site | Longitude | Latitude | No. years with counts | Time span | Total no. monarchs | Total party hours | Abundance trend (%/year) | p value |
| --- | --- | --- | --- | --- | --- | --- | --- | --- |
| ACAS, MD. | -77.1931 | 39.4234 | 20 | 1996-2018 | 78 | 191.67 | 6.8 | 0.051 |
| Adams Co. | -83.4136 | 38.7718 | 23 | 1993-2018 | 279 | 743.95 | 0.8 | 0.774 |
| Airlie Center, VA. | -77.7943 | 38.7566 | 15 | 1995-2018 | 262 | 254.5 | 4.8 | 0.531 |
| Akron, OH. | -81.5637 | 41.0605 | 11 | 2002-2018 | 66 | 125.5 | 8.4 | 0.129 |
| Alexandria, LA. | -92.467 | 31.2742 | 9 | 2000-2018 | 28 | 70.5 | 3.0 | 0.489 |
| Algonquin East, ON, Canada. | -77.9 | 45.9833 | 9 | 1998-2010 | 117 | 269 | -3.7 | 0.899 |
| Algonquin Park, ON, Canada. | -78.4 | 45.5667 | 18 | 1995-2014 | 979 | 982 | -0.6 | 0.915 |
| Allegan, MI. | -86.0074 | 42.5492 | 13 | 2001-2018 | 97 | 102.25 | -0.8 | 0.805 |
| Allen Co. | -85.0833 | 41.2167 | 12 | 1996-2009 | 62 | 71 | -0.9 | 0.963 |
| Alva, FL. | -81.6726 | 26.7153 | 16 | 1995-2018 | 33 | 104 | 3.6 | 0.224 |
| Ann Arbor City, MI. | -83.7066 | 42.2805 | 16 | 1999-2018 | 235 | 374 | 6.2 | 0.736 |
| Archer, FL. | -82.3392 | 29.6667 | 14 | 1998-2018 | 70 | 129 | 4.1 | 0.464 |
| Arkansas City, KS. | -97.0385 | 37.0412 | 10 | 2007-2018 | 36 | 147 | 2.1 | 0.808 |
| Ashe County, NC. | -81.45 | 36.4 | 7 | 1995-2004 | 21 | 63 | -1.6 | 0.672 |
| Ashland Co. | -82.3393 | 40.7191 | 8 | 2009-2018 | 109 | 104 | -14.9 | 0.138 |
| Atascosa Highlands, AZ. | -111.16 | 31.48 | 7 | 1994-2001 | 66 | 357.5 | 24.8 | 0.084 |
| Aullwood Audubon Center and Farm, OH. | -84.2754 | 39.8732 | 23 | 1993-2018 | 301 | 192.75 | -2.4 | 0.421 |
| Avoca | -90.4267 | 43.1829 | 12 | 2007-2018 | 187 | 80.5 | 20.3 | 0.268 |
| Baker Ponds, NH. | -71.9685 | 43.8898 | 10 | 2006-2018 | 213 | 121 | -3.7 | 0.726 |
| Bald Mountain Recreation Area, MI. | -83.21 | 42.75 | 13 | 1997-2012 | 131 | 49 | 4.8 | 0.388 |
| Baraboo, WI. | -89.7387 | 43.4891 | 26 | 1993-2018 | 508 | 205.98 | 5.0 | 0.039 |
| Barataria, LA. | -90.12 | 29.72 | 7 | 1996-2007 | 16 | 92 | -13.9 | 0.512 |
| Barnegat, NJ. | -74.2231 | 39.7533 | 14 | 2004-2018 | 210 | 101.5 | -0.1 | 0.991 |
| Barnstable | -70.2996 | 41.7012 | 11 | 2005-2017 | 347 | 71 | 4.7 | 0.781 |
| Bath Co. | -79.8028 | 38 | 19 | 1995-2017 | 195 | 271 | -5.8 | 0.226 |
| Bear Trap Junction, MN. | -92.4833 | 46.8833 | 12 | 2001-2013 | 116 | 72.33 | 12.6 | 0.244 |
| Beardstown, IL. | -90.4278 | 39.9484 | 17 | 1995-2012 | 331 | 94.75 | -14.6 | 0.285 |
| Beaver Creek Reserve, WI. | -91.2701 | 44.8134 | 17 | 2001-2018 | 1156 | 263.5 | -10.2 | 0.081 |
| Belleplain, NJ. | -74.9333 | 39.25 | 20 | 1993-2015 | 529 | 748.5 | -4.4 | 0.414 |
| Bemidji, MN. | -94.8837 | 47.4601 | 24 | 1995-2018 | 3059 | 356.97 | 7.7 | 0.076 |
| Benicia, CA. | -122.1847 | 38.0691 | 7 | 1998-2016 | 19 | 73 | -2.6 | 0.701 |
| Bentsen-Rio Grande Valley State Park, TX. | -98.3135 | 26.1915 | 5 | 2004-2016 | 82 | 57.5 | 33.3 | 0.070 |
| Berkeley, CA. | -122.2577 | 37.872 | 14 | 1996-2017 | 29 | 402.5 | 4.7 | 0.143 |
| Big Bear, CA. | -116.8249 | 34.2432 | 6 | 1997-2011 | 14 | 293 | -9.9 | 0.222 |
| Big Chico Creek Ecological Reserve | -121.7021 | 39.8445 | 7 | 2009-2018 | 26 | 107.5 | 12.8 | 0.544 |
| Big Oaks National Wildlife Refuge, IN. | -85.4167 | 38.9333 | 19 | 1999-2018 | 952 | 411 | 4.1 | 0.400 |
| Big Pine Key, FL. | -81.3699 | 24.6908 | 7 | 1998-2018 | 17 | 77.5 | 3.3 | 0.339 |
| Binghamton, NY. | -75.9 | 42.1167 | 12 | 1996-2009 | 75 | 98.75 | -4.7 | 0.737 |
| Blackstone Valley Corridor, MA. | -71.6117 | 42.1243 | 14 | 2001-2018 | 344 | 466.25 | 2.8 | 0.839 |
| Blue Mounds, MN. | -96.1127 | 43.7035 | 22 | 1997-2018 | 601 | 127.37 | 0.2 | 0.963 |
| Blue Ridge Mountains, VA. | -78.88 | 37.82 | 6 | 1997-2007 | 26 | 40.5 | -3.1 | 0.889 |
| Blue Ridge Parkway, VA. | -79.0326 | 37.9609 | 23 | 1993-2018 | 235 | 114.34 | -4.1 | 0.130 |
| Bluestem, MN. | -96.4734 | 46.8552 | 16 | 1993-2018 | 505 | 144.92 | 15.7 | 0.005 |
| Boise Front, ID. | -116.144 | 43.7127 | 8 | 1997-2016 | 26 | 65.5 | 11.6 | 0.000 |
| Bond Swamp National Wildlife Refuge, GA. | -83.5765 | 32.7288 | 9 | 2001-2018 | 17 | 214 | -3.6 | 0.067 |
| Bonnet Carre Spillway, LA. | -90.4583 | 30 | 9 | 1998-2016 | 16 | 55.5 | 2.9 | 0.360 |
| Boone County, IN. | -86.25 | 39.95 | 8 | 1997-2006 | 55 | 48.75 | 1.6 | 0.876 |
| Brazos Bend, TX. | -95.6688 | 29.3734 | 7 | 1995-2004 | 13 | 162.25 | 8.4 | 0.004 |
| Brazos Valley, TX. | -96.0502 | 29.9008 | 10 | 2003-2014 | 31 | 135.55 | -7.9 | 0.146 |
| Brewster, MA. | -70.0167 | 41.7833 | 12 | 2004-2015 | 294 | 145 | 15.1 | 0.428 |
| Bristol Co. | -70.9679 | 41.6326 | 18 | 1993-2016 | 107 | 95.5 | 1.2 | 0.796 |
| BronxManhattan Counties, NY. | -73.9 | 40.86 | 10 | 1993-2005 | 52 | 339.5 | 2.5 | 0.676 |
| Broward Co. North | -80.157 | 26.2313 | 15 | 2004-2018 | 109 | 241.75 | 9.8 | 0.083 |
| Broward Co. South | -80.3135 | 26.0634 | 15 | 2004-2018 | 121 | 229.5 | 6.2 | 0.022 |
| Bryce Canyon, UT. | -112.1576 | 37.6055 | 5 | 2001-2014 | 7 | 37.5 | 4.9 | 0.058 |
| Bryn Mawr, PA. | -75.3406 | 40.0689 | 23 | 1993-2018 | 419 | 298.75 | 1.3 | 0.722 |
| Buffalo Jump Prairie, MO. | -93.48 | 37.77 | 6 | 1993-2000 | 32 | 26.8 | -1.7 | 0.899 |
| Bull Creek, FL. | -80.9672 | 28.0844 | 12 | 2007-2018 | 195 | 239 | 22.3 | 0.044 |
| Busch Conservation Area, MO. | -90.74 | 38.7051 | 13 | 1998-2018 | 34 | 115.4 | 6.9 | 0.369 |
| Byron Hatchery Watchable Wildlife Area, OK. | -98.18 | 36.83 | 5 | 1998-2005 | 12 | 55 | -5.5 | 0.582 |
| Caesar Creek Lake, OH. | -84.0582 | 39.4881 | 5 | 2002-2010 | 18 | 23.5 | 14.7 | 0.422 |
| Cal-Wood Outdoor Learning Center, CO. | -105.3902 | 40.15 | 7 | 2000-2014 | 11 | 150.5 | -1.3 | 0.801 |
| Camas National Wildlife Refuge, ID. | -112.2643 | 43.9653 | 12 | 2005-2018 | 77 | 77.5 | -2.9 | 0.743 |
| Cambridge (rare Charitable Research Reserve), ON. | -80.355 | 43.3817 | 11 | 2006-2018 | 323 | 193.5 | -9.3 | 0.332 |
| Cape May, NJ. | -74.8667 | 39.0167 | 26 | 1993-2018 | 1685 | 1258.25 | 5.6 | 0.346 |
| Carden, ON, Canada. | -79.05 | 44.6333 | 16 | 1998-2015 | 1003 | 405 | 1.7 | 0.739 |
| Carlton County, MN. | -92.34 | 46.64 | 14 | 1993-2007 | 160 | 87.5 | 12.7 | 0.019 |
| Central Berkshire Co. | -73.2537 | 42.4086 | 21 | 1993-2018 | 698 | 517 | -4.0 | 0.270 |
| Central Franklin Co. | -72.5097 | 42.5795 | 24 | 1993-2018 | 703 | 947.5 | -3.3 | 0.430 |
| Central Polk Co. | -95.3658 | 47.595 | 12 | 2007-2018 | 523 | 144 | -9.1 | 0.379 |
| Central Polk County, MN. | -95.37 | 47.6 | 13 | 1994-2006 | 582 | 107.9 | 40.1 | 0.044 |
| Central Suffolk Co. | -72.7536 | 40.8627 | 14 | 2005-2018 | 263 | 200.5 | -12.4 | 0.054 |
| Chagrin River Valley, OH. | -81.4382 | 41.4784 | 19 | 1996-2017 | 283 | 291.5 | 3.1 | 0.553 |
| Chatfield Nature Preserve, CO. | -105.17 | 39.88 | 5 | 1994-2002 | 9 | 18 | 5.5 | 0.456 |
| Chautauqua, NY. | -79.1639 | 42.271 | 14 | 1993-2012 | 218 | 87.25 | -3.5 | 0.308 |
| Chelsea, MI. | -84.0204 | 42.3181 | 23 | 1996-2018 | 247 | 324.25 | -2.2 | 0.540 |
| Chippewa Nature Center, MI. | -84.42 | 43.62 | 5 | 1993-1997 | 159 | 46.5 | -3.1 | 0.932 |
| Chippokes Plantation State Park, VA. | -76.7208 | 37.1397 | 16 | 2000-2018 | 67 | 285 | -0.5 | 0.836 |
| Choctawhatchee Bay Northwest | -86.4974 | 30.524 | 9 | 2001-2016 | 15 | 126.75 | 4.6 | 0.207 |
| Christmas, FL. | -81 | 28.5 | 24 | 1993-2018 | 173 | 815.75 | 6.0 | 0.001 |
| Clear Creek, ON, Canada | -81.7729 | 42.5147 | 15 | 2001-2015 | 1995 | 514.5 | -2.5 | 0.679 |
| Cleveland County Audubon Society, OK. | -97.33 | 35.19 | 10 | 1993-2003 | 26 | 260 | -3.5 | 0.572 |
| Cleveland Heights-Holden Arboretum, OH. | -81.36 | 41.69 | 14 | 1993-2006 | 416 | 165.5 | 6.0 | 0.408 |
| Coffee Co. | -86.1 | 35.4833 | 5 | 1998-2009 | 12 | 68 | -13.4 | 0.099 |
| Colt Creek | -82.0558 | 28.3128 | 8 | 2010-2018 | 16 | 187 | 1.3 | 0.930 |
| Concord, MA. | -71.4167 | 42.4333 | 18 | 1994-2018 | 473 | 291.5 | -5.7 | 0.205 |
| Cook Co. | -90.3438 | 47.8548 | 8 | 2007-2018 | 45 | 22 | -15.1 | 0.017 |
| Coral Gables, FL. | -80.269 | 25.7051 | 23 | 1995-2018 | 339 | 429.5 | 10.1 | 0.000 |
| Corkscrew Swamp Sanctuary, FL. | -81.5217 | 26.3666 | 5 | 1995-2015 | 6 | 96 | 2.2 | 0.060 |
| Cornucopia, WI. | -91.0932 | 46.8037 | 19 | 1994-2018 | 332 | 138.04 | -0.4 | 0.944 |
| Cosumnes River, CA. | -121.4358 | 38.3503 | 20 | 1994-2017 | 223 | 221.8 | 0.2 | 0.971 |
| Crex Meadows, WI. | -92.6062 | 45.8455 | 26 | 1993-2018 | 1059 | 146.61 | 4.2 | 0.250 |
| Croatan National Forest, NC. | -76.98 | 34.87 | 10 | 1999-2012 | 90 | 118.25 | -3.3 | 0.803 |
| Cumberland Co. | -75.2004 | 39.3325 | 22 | 1993-2015 | 2209 | 849.25 | 4.3 | 0.633 |
| Danby-Tinmouth, VT. | -73.0833 | 43.35 | 8 | 1999-2008 | 121 | 55.25 | 23.5 | 0.424 |
| Dardenelles, CA. | -119.8333 | 38.3411 | 15 | 1994-2010 | 59 | 243.5 | -4.4 | 0.331 |
| Decatur Co., TN. | -88.0338 | 35.6278 | 9 | 2006-2018 | 37 | 88.5 | -5.8 | 0.749 |
| Deep Portage, MN. | -94.3849 | 46.8995 | 25 | 1993-2018 | 723 | 262 | 3.7 | 0.137 |
| Delmarva Tip, VA. | -75.9576 | 37.2024 | 18 | 1999-2018 | 266 | 516 | -2.2 | 0.731 |
| Delta National Forest, MS. | -90.7558 | 32.7597 | 13 | 2006-2018 | 109 | 206.5 | 17.4 | 0.028 |
| Dismal Swamp, VA. | -76.483 | 36.6718 | 14 | 1993-2015 | 63 | 344.25 | 0.4 | 0.951 |
| Disney Wilderness Preserve, FL. | -81.4082 | 28.124 | 14 | 1999-2018 | 49 | 316.5 | 8.2 | 0.112 |
| Dubuque, IA. | -90.7 | 42.4833 | 18 | 1999-2018 | 536 | 135.4 | -1.6 | 0.620 |
| Duke Farms, NJ. | -74.6891 | 40.6096 | 11 | 2006-2018 | 842 | 150 | -1.5 | 0.879 |
| Durham, NC. | -78.9008 | 36.0935 | 18 | 1999-2018 | 903 | 561.1 | 1.9 | 0.847 |
| East Bedford Co. | -78.6167 | 40.1167 | 19 | 2000-2018 | 737 | 553 | 1.5 | 0.837 |
| East Glastonbury, CT. | -72.5333 | 41.6667 | 11 | 1995-2009 | 181 | 124.75 | 3.9 | 0.863 |
| East Haddam, CT. | -72.4 | 41.48 | 6 | 1995-2000 | 114 | 71.5 | -3.1 | 0.710 |
| Eastern Catskills | -74.0298 | 42.0565 | 9 | 2007-2018 | 273 | 318 | -7.6 | 0.172 |
| El Dorado Nature Center, CA. | -118.0869 | 33.8095 | 11 | 2005-2017 | 83 | 54 | -13.6 | 0.016 |
| El Dorado Springs, MO. | -94 | 37.75 | 8 | 1993-2002 | 240 | 36.54 | 15.5 | 0.239 |
| Elizabethton, TN. | -82.1324 | 36.3626 | 12 | 2002-2018 | 84 | 105 | -0.3 | 0.967 |
| Elizabethtown, NY. | -73.6 | 44.22 | 10 | 1994-2003 | 108 | 104 | -22.7 | 0.114 |
| Emmet and Cheboygan Counties, MI. | -84.7508 | 45.4345 | 19 | 1993-2018 | 564 | 242 | -10.5 | 0.021 |
| Erie Co. | -82.5667 | 41.3333 | 7 | 1999-2008 | 76 | 46.75 | 33.1 | 0.191 |
| Eugene, OR. | -123.0886 | 44.0335 | 8 | 2000-2017 | 17 | 137.95 | 1.1 | 0.778 |
| Falmouth, MA. | -70.585 | 41.6095 | 19 | 1999-2018 | 245 | 165.45 | 1.3 | 0.883 |
| Farmington Hills, MI. | -83.3748 | 42.4697 | 17 | 1994-2018 | 125 | 85 | 3.8 | 0.272 |
| Farmington Valley, CT. | -72.8674 | 41.8169 | 22 | 1996-2018 | 511 | 455 | 9.9 | 0.056 |
| FaulknerPulaski Counties | -92.3983 | 34.9017 | 5 | 2003-2018 | 17 | 130.5 | -4.9 | 0.321 |
| Ferrisburg, VT. | -73.25 | 44.22 | 15 | 1993-2007 | 152 | 112.5 | 12.0 | 0.120 |
| Fields Pond Audubon Center, ME. | -68.7277 | 44.7384 | 8 | 2004-2018 | 43 | 81.5 | -4.8 | 0.562 |
| Fincastle, VA. | -79.8806 | 37.5122 | 19 | 1995-2018 | 78 | 377 | 0.2 | 0.943 |
| Finger Lakes National Forest, NY. | -76.7908 | 42.529 | 17 | 1997-2018 | 68 | 129.25 | 0.2 | 0.946 |
| Flagler Co. Northeast | -81.2187 | 29.5539 | 6 | 2001-2018 | 26 | 48.5 | 1.6 | 0.791 |
| Forsyth Co. | -80.2936 | 36.1369 | 15 | 1995-2015 | 114 | 153 | 6.1 | 0.252 |
| Fort QuAppelle, SK, Canada. | -103.7836 | 50.7696 | 12 | 1994-2012 | 60 | 73.25 | 12.3 | 0.052 |
| Fox River-Seney, MI. | -85.95 | 46.35 | 24 | 1994-2018 | 1494 | 235.63 | -0.3 | 0.924 |
| Francis Marion National Forest, SC. | -79.4754 | 33.1408 | 10 | 1996-2014 | 48 | 265 | 8.3 | 0.135 |
| Furnace Hills, PA. | -76.3027 | 40.2262 | 25 | 1993-2018 | 309 | 445.21 | -0.5 | 0.891 |
| Galloway Township, NJ. | -74.5605 | 39.485 | 22 | 1993-2014 | 328 | 204.78 | 2.2 | 0.707 |
| Giant Sequoia National Monument South | -118.4833 | 36 | 13 | 2003-2016 | 142 | 319.5 | -3.3 | 0.275 |
| Giles County, VA. | -80.63 | 37.32 | 10 | 1997-2006 | 78 | 195.75 | -4.1 | 0.720 |
| Gilmanton, WI. | -91.6788 | 44.4221 | 13 | 2005-2018 | 364 | 75.5 | 1.8 | 0.795 |
| Gilpin Co. | -105.4105 | 39.8309 | 12 | 1994-2015 | 36 | 484 | -5.8 | 0.066 |
| Goethe State Forest, FL. | -82.6394 | 29.0836 | 5 | 2006-2013 | 11 | 98.5 | -12.4 | 0.229 |
| Goose Pond | -87.2317 | 39.031 | 11 | 2008-2018 | 576 | 258.5 | 12.9 | 0.770 |
| Gratiot-Saginaw, MI. | -84.409 | 43.2486 | 15 | 1997-2014 | 91 | 98 | 1.2 | 0.779 |
| Great Plains State Park, OK. | -98.9873 | 34.7387 | 14 | 1996-2015 | 83 | 211.75 | 2.6 | 0.414 |
| Great Swamp, NJ. | -74.4624 | 40.78 | 24 | 1993-2018 | 542 | 724 | -2.7 | 0.480 |
| Greater Muskegon, MI. | -86.1643 | 43.2342 | 23 | 1993-2018 | 412 | 184.35 | 3.5 | 0.228 |
| Green Island-Lost Mound | -90.3435 | 42.1956 | 8 | 2008-2018 | 390 | 46.5 | -5.8 | 0.730 |
| Green River Wildlife Area, IL. | -89.5161 | 41.585 | 12 | 2001-2012 | 655 | 126.25 | -4.3 | 0.845 |
| Greenbrook Sanctuary, NJ. | -73.9327 | 40.9144 | 6 | 1994-2017 | 10 | 40 | -4.3 | 0.019 |
| Grindstone Mountain, TN. | -85.0586 | 35.0834 | 9 | 2002-2018 | 235 | 150.5 | 8.2 | 0.535 |
| Haliburton Highlands, ON, Canada. | -78.6833 | 44.8833 | 14 | 2000-2014 | 2938 | 797 | 3.3 | 0.490 |
| Halls Mills, NY. | -74.5859 | 41.8927 | 16 | 2003-2018 | 445 | 326 | -0.5 | 0.949 |
| Harford Co. | -76.2333 | 39.55 | 9 | 2007-2018 | 46 | 54 | 32.9 | 0.139 |
| Harris Neck National Wildlife Refuge, GA. | -81.3 | 31.6833 | 9 | 1993-2013 | 21 | 83.5 | -1.7 | 0.381 |
| Harris Ranch | -101.85 | 31.75 | 5 | 2007-2016 | 2983 | 107 | 108.8 | 0.014 |
| Harvey Co. | -97.4667 | 38.0667 | 10 | 2007-2016 | 117 | 232.5 | -9.4 | 0.510 |
| Hastings Reservation, CA. | -121.4993 | 36.336 | 14 | 1998-2017 | 74 | 219.58 | 4.0 | 0.273 |
| Hawk Mountain, PA. | -75.9167 | 40.5833 | 25 | 1993-2018 | 462 | 566.5 | 7.6 | 0.217 |
| Hazleton | -75.9167 | 40.9667 | 9 | 1996-2008 | 32 | 82.5 | -4.7 | 0.415 |
| Headwaters | -83.5434 | 42.7725 | 10 | 2007-2018 | 58 | 73.5 | -1.4 | 0.899 |
| Hendricks, PA. | -75.47 | 40.33 | 9 | 1995-2005 | 209 | 154.25 | -16.1 | 0.044 |
| High Falls, GA. | -84.0179 | 33.1787 | 5 | 2004-2009 | 29 | 74 | -38.3 | 0.413 |
| High Island, TX. | -94.47 | 29.55 | 5 | 1996-2002 | 11 | 63 | -17.1 | 0.268 |
| High Line Canal, CO. | -104.8472 | 39.7243 | 11 | 1994-2008 | 50 | 83 | -9.2 | 0.195 |
| Highlands Hammock, FL. | -81.564 | 27.4343 | 12 | 1999-2017 | 19 | 186 | 3.0 | 0.237 |
| Hillsdale, MI. | -84.593 | 41.8544 | 18 | 1997-2018 | 116 | 139 | -4.0 | 0.304 |
| Hobcaw Barony, SC. | -79.1645 | 33.4312 | 8 | 1997-2014 | 26 | 170 | 6.0 | 0.097 |
| Hog Island, ON, Canada. | -76.9158 | 45.842 | 23 | 1993-2015 | 763 | 259.5 | 1.0 | 0.803 |
| Holden Arboretum | -81.3592 | 41.6868 | 12 | 2007-2018 | 701 | 242 | -7.8 | 0.142 |
| Holtwood | -76.315 | 39.9094 | 18 | 1996-2018 | 197 | 248.55 | 1.0 | 0.796 |
| Homestead | -80.51 | 25.4916 | 8 | 2002-2013 | 41 | 116 | 12.5 | 0.561 |
| Horseshoe Lake, IL. | -90.0915 | 38.695 | 13 | 2001-2011 | 408 | 107 | -1.5 | 0.924 |
| Houston, TX. | -95.3667 | 29.7333 | 12 | 2000-2018 | 92 | 170.5 | -2.2 | 0.504 |
| Indian Cave State Park, NE. | -95.6149 | 40.3151 | 23 | 1993-2015 | 1085 | 173.33 | -3.3 | 0.447 |
| Iowa City, IA. | -91.6076 | 41.7522 | 20 | 1999-2018 | 709 | 184.85 | 5.8 | 0.208 |
| Iredell Co. | -80.85 | 35.9167 | 6 | 2000-2012 | 51 | 46 | -17.7 | 0.020 |
| Island Ford, VA. | -78.6899 | 38.3519 | 17 | 1999-2018 | 122 | 275.4 | -1.0 | 0.835 |
| Itasca State Park, MN. | -95.1709 | 47.1917 | 25 | 1994-2018 | 2611 | 545.65 | 9.3 | 0.044 |
| Jackson | -88.7764 | 35.5513 | 8 | 2007-2018 | 28 | 87.5 | -0.1 | 0.963 |
| Johnson County, IN. | -86.07 | 39.48 | 9 | 1993-2002 | 40 | 56 | -14.4 | 0.200 |
| Key West, FL. | -81.6968 | 24.586 | 17 | 2001-2018 | 141 | 147.5 | 5.0 | 0.304 |
| Killarney Provincial Park, ON, Canada. | -81.4046 | 46.0139 | 13 | 2001-2015 | 514 | 304 | 0.2 | 0.983 |
| Kissimmee Prairie Sanctuary, FL. | -80.9813 | 27.5526 | 6 | 1998-2018 | 9 | 131 | 2.5 | 0.312 |
| La Porte Co. | -86.8252 | 41.6167 | 10 | 2007-2018 | 58 | 52.5 | -0.4 | 0.971 |
| Lake Bronson State Park, MN. | -96.53 | 48.75 | 11 | 1995-2006 | 270 | 82.9 | 32.5 | 0.152 |
| Lake Co. East | -81.0548 | 41.723 | 21 | 1997-2018 | 144 | 393.25 | 5.9 | 0.226 |
| Lake Dore, ON, Canada. | -77.08 | 45.64 | 19 | 1993-2012 | 574 | 215 | -2.2 | 0.730 |
| Lake Meyer | -91.9039 | 43.1688 | 10 | 2008-2018 | 98 | 106 | -4.6 | 0.598 |
| Lake Placid, NY. | -73.9833 | 44.25 | 21 | 1994-2018 | 332 | 158.75 | 9.6 | 0.003 |
| Lakehurst, NJ. | -74.37 | 40 | 10 | 1995-2005 | 109 | 232.3 | -18.5 | 0.027 |
| LaSalle County, IL. | -88.95 | 41.2 | 12 | 1993-2005 | 326 | 69.5 | -5.8 | 0.442 |
| Lava Beds National Monument | -121.5253 | 41.7668 | 10 | 2009-2018 | 120 | 363.5 | 14.9 | 0.342 |
| Letchworth State Park, NY. | -77.9713 | 42.6573 | 14 | 1996-2010 | 359 | 384.5 | 6.9 | 0.562 |
| Lexington, VA. | -79.477 | 37.848 | 11 | 1997-2010 | 67 | 137 | 8.8 | 0.046 |
| Long Point, ON, Canada. | -80.3833 | 42.6167 | 23 | 1993-2015 | 1145 | 991.5 | 1.4 | 0.670 |
| Lookout Mountain, GA. | -85.3893 | 34.8074 | 9 | 2001-2017 | 16 | 106.95 | 6.9 | 0.066 |
| Lower East Pearl River | -89.7372 | 30.387 | 14 | 1993-2016 | 75 | 207.75 | 0.4 | 0.887 |
| Lower Hatchie | -89.7774 | 35.6525 | 10 | 2008-2018 | 40 | 99 | 24.5 | 0.404 |
| Lower Penobscot, ME. | -68.8999 | 44.5466 | 6 | 2006-2014 | 82 | 61 | -8.3 | 0.265 |
| MacGregor Point Provincial Park, ON, Canada. | -81.4384 | 44.3731 | 15 | 1998-2013 | 1532 | 500.8 | 4.2 | 0.443 |
| Madison, WI. | -89.3838 | 43.0899 | 25 | 1993-2018 | 447 | 395.9 | -0.6 | 0.701 |
| Magnolia, TX. | -95.7109 | 30.2738 | 6 | 2004-2009 | 34 | 50.4 | 14.7 | 0.598 |
| Maidens, VA. | -77.9575 | 37.6423 | 12 | 1996-2014 | 40 | 141.5 | 5.4 | 0.272 |
| Manion Corners, ON, Canada. | -76.0653 | 45.2565 | 13 | 1999-2015 | 314 | 426.5 | 14.9 | 0.148 |
| Manitoulin Island, ON, Canada. | -82.75 | 45.8333 | 11 | 1994-2009 | 167 | 70 | 11.6 | 0.184 |
| Marin Co. | -122.5655 | 37.9663 | 9 | 1997-2015 | 10 | 94 | -0.1 | 0.946 |
| Mariton Wildlife Sanctuary, PA. | -75.2044 | 40.6093 | 12 | 1995-2018 | 83 | 40.95 | 5.2 | 0.362 |
| Martha LaFite Thompson Nature Sanctuary, MO. | -94.3133 | 39.32 | 12 | 2006-2018 | 43 | 86.5 | -0.2 | 0.966 |
| Marthas Vineyard, MA. | -70.6333 | 41.3667 | 9 | 1999-2008 | 520 | 204.25 | 6.8 | 0.794 |
| Mary Gray Bird Sanctuary, IN. | -85.2072 | 39.5765 | 20 | 1993-2018 | 180 | 157.33 | 0.2 | 0.949 |
| Mason Co. | -89.9579 | 40.1628 | 11 | 2001-2014 | 194 | 87.5 | -1.4 | 0.902 |
| McDonough Co. | -90.7719 | 40.3683 | 17 | 1994-2014 | 177 | 86.68 | 0.9 | 0.753 |
| McNair, MN. | -91.6717 | 47.32 | 6 | 2005-2017 | 12 | 56.9 | -14.4 | 0.042 |
| Mercer, NJ. | -74.7663 | 40.2694 | 16 | 2003-2018 | 630 | 491.5 | 36.3 | 0.285 |
| Meriden, CT. | -72.7448 | 41.5243 | 17 | 1993-2018 | 94 | 155.5 | 5.1 | 0.212 |
| Metro New Orleans, LA. | -90.112 | 29.9416 | 19 | 1993-2012 | 840 | 1550.5 | 0.7 | 0.976 |
| Midland, TX. | -102.0279 | 31.9869 | 14 | 1994-2017 | 48 | 335 | -2.3 | 0.458 |
| Milwaukee County (Riverside), WI. | -88.03 | 43.13 | 8 | 1994-2005 | 148 | 39.45 | 0.7 | 0.922 |
| Minidoka National Wildlife Refuge, ID. | -113.3544 | 42.6479 | 10 | 2001-2018 | 71 | 65 | 12.5 | 0.480 |
| Moccasin Wallow | -82.5645 | 27.5581 | 5 | 2009-2017 | 11 | 38.5 | -4.1 | 0.845 |
| Mohawk Dam, OH. | -82.0821 | 40.4245 | 12 | 2005-2017 | 283 | 312.5 | -15.6 | 0.001 |
| Monastery-Panola-Arabia Mountains, GA. | -84.0804 | 33.6191 | 6 | 2003-2011 | 59 | 209.5 | -14.5 | 0.523 |
| MonroeBrown, IN. | -86.3277 | 39.094 | 15 | 1997-2018 | 182 | 502 | 4.9 | 0.216 |
| Monterey | -121.8185 | 36.5418 | 6 | 2011-2016 | 20 | 201.5 | 38.4 | 0.002 |
| Monticello, NY. | -74.6321 | 41.5684 | 24 | 1994-2018 | 356 | 378.55 | 4.7 | 0.429 |
| Montour Preserve, PA. | -76.5479 | 41.0632 | 14 | 2001-2015 | 142 | 123.5 | -6.9 | 0.621 |
| Moose Lake State Park, MN. | -92.7304 | 46.4393 | 16 | 2003-2018 | 284 | 78.2 | -3.5 | 0.339 |
| Mount Baden-Powell, CA. | -117.77 | 34.37 | 8 | 1996-2005 | 25 | 176.25 | -16.6 | 0.059 |
| Mount Diablo, CA. | -121.8871 | 37.9511 | 20 | 1993-2014 | 155 | 635.32 | 2.1 | 0.825 |
| Mount Lassen | -121.404 | 40.4609 | 11 | 2007-2018 | 50 | 494.5 | 4.7 | 0.708 |
| Mount Magazine, AR. | -93.63 | 35.17 | 9 | 1993-2002 | 347 | 88.5 | 54.9 | 0.184 |
| Mount Snow, VT. | -72.8946 | 42.9682 | 15 | 1999-2018 | 101 | 129.51 | -5.1 | 0.306 |
| Mud Lake, WI. | -89.3044 | 43.4118 | 26 | 1993-2018 | 999 | 271.1 | 7.3 | 0.007 |
| Muriel Shirey Memorial, MN. | -92.125 | 46.7767 | 8 | 2006-2018 | 27 | 35.5 | -5.4 | 0.632 |
| Muscatatuck National Wildlife Refuge, IN. | -85.8 | 38.95 | 5 | 2002-2013 | 57 | 41 | -19.2 | 0.009 |
| Muskoka, ON, Canada. | -79.6196 | 45.0219 | 13 | 1999-2015 | 314 | 181 | -19.3 | 0.274 |
| Muttontown, NY. | -73.5352 | 40.853 | 26 | 1993-2018 | 2228 | 927 | -4.8 | 0.107 |
| Nashville South | -86.8667 | 36.0167 | 10 | 2008-2018 | 35 | 162 | -3.6 | 0.540 |
| Nassawango Creek, MD. | -75.4714 | 38.229 | 21 | 1994-2018 | 287 | 390.63 | -0.9 | 0.892 |
| Nelson Co. | -78.8999 | 37.8323 | 7 | 2008-2015 | 19 | 86.5 | -14.1 | 0.215 |
| Nevis, MN. | -94.8437 | 46.985 | 31 | 1995-2016 | 1627 | 332.93 | 7.6 | 0.001 |
| Nickel Preserve, OK. | -94.842 | 36.0239 | 7 | 2001-2015 | 33 | 85.5 | 8.8 | 0.345 |
| Niobrara Valley Preserve, NE. | -100.03 | 42.78 | 8 | 1993-2004 | 47 | 58 | -7.1 | 0.098 |
| Nixon Park, PA. | -76.7322 | 39.8854 | 8 | 2000-2014 | 127 | 105 | -6.7 | 0.606 |
| Noble Co. | -85.4246 | 41.3961 | 18 | 1996-2018 | 150 | 149 | 5.9 | 0.304 |
| North Delaware (Wilmington), DE. | -75.6416 | 39.7153 | 14 | 1996-2014 | 365 | 182.05 | 11.9 | 0.013 |
| North Essex Co. | -70.9091 | 42.7052 | 18 | 1998-2018 | 401 | 350.5 | -9.6 | 0.310 |
| North Fork Kern River Valley, CA. | -118.4706 | 35.7735 | 10 | 2001-2016 | 53 | 288.25 | -10.0 | 0.436 |
| North Mountain, PA. | -76.17 | 41.32 | 10 | 1998-2007 | 292 | 132 | 2.7 | 0.906 |
| North Park Village, IL. | -87.68 | 41.98 | 8 | 1997-2004 | 43 | 34.5 | 4.2 | 0.549 |
| Northampton, MA. | -72.6043 | 42.3751 | 18 | 2001-2018 | 2039 | 763.75 | 3.8 | 0.736 |
| Northern Berkshire Co. | -73.1254 | 42.6279 | 20 | 1993-2018 | 628 | 519 | -5.8 | 0.105 |
| Northern Kettle Moraine, WI. | -88.1603 | 43.665 | 22 | 1997-2018 | 575 | 190.5 | -3.2 | 0.196 |
| Northern Westchester Co. | -73.6804 | 41.2012 | 25 | 1993-2018 | 2829 | 1608 | -7.6 | 0.094 |
| Northern Worcester Co. | -71.8122 | 42.4333 | 20 | 1996-2018 | 1147 | 874.5 | -3.7 | 0.488 |
| Noxubee National Wildlife Refuge, MS. | -88.7929 | 33.2783 | 25 | 1993-2018 | 160 | 632.25 | 2.2 | 0.341 |
| Occoquan Bay, VA. | -77.2404 | 38.6392 | 18 | 2001-2018 | 129 | 294.5 | 15.4 | 0.006 |
| Ochoco, OR. | -120.2667 | 44.3667 | 5 | 1994-2016 | 7 | 63.5 | -2.3 | 0.422 |
| Okeechobee, FL. | -80.8282 | 27.2898 | 14 | 2005-2018 | 107 | 170 | 7.9 | 0.590 |
| Olivet, MI. | -84.9286 | 42.4628 | 18 | 1994-2018 | 111 | 118.25 | 2.5 | 0.341 |
| Organ Mountains, NM. | -106.65 | 32.3666 | 9 | 1993-2010 | 39 | 159.25 | 3.1 | 0.582 |
| Orillia, ON, Canada. | -79.38 | 44.63 | 12 | 1994-2007 | 563 | 228.25 | 15.3 | 0.195 |
| Oshawa, ON, Canada. | -78.9258 | 44.0181 | 22 | 1993-2015 | 1814 | 1046.7 | -1.0 | 0.879 |
| Ottawa Co. - Grand River | -85.9317 | 43.0243 | 12 | 2002-2015 | 132 | 102.5 | -7.9 | 0.415 |
| Owego, NY. | -76.2333 | 42.1 | 9 | 1999-2008 | 91 | 78 | 11.1 | 0.498 |
| Owls Hill (Nashville), TN. | -86.87 | 36.02 | 9 | 1998-2007 | 24 | 117.25 | 7.5 | 0.287 |
| Palm Beach Co. Central | -80.1741 | 26.6509 | 24 | 1995-2018 | 353 | 369.25 | 0.1 | 0.985 |
| Palm Beach Co. North | -80.1311 | 26.883 | 23 | 1995-2018 | 123 | 421.25 | -1.1 | 0.788 |
| Palm Beach Co. South | -80.1466 | 26.4391 | 23 | 1996-2018 | 359 | 346.5 | 0.9 | 0.718 |
| Palm Harbor, FL. | -82.7333 | 28.05 | 5 | 1995-2015 | 175 | 118.5 | -3.9 | 0.152 |
| Palos Verdes, CA. | -118.3667 | 33.7333 | 19 | 1994-2016 | 234 | 542 | 10.2 | 0.112 |
| Patagonia, AZ. | -110.6908 | 31.5072 | 21 | 1993-2014 | 342 | 1307.5 | -3.0 | 0.550 |
| Peaks of Otter, VA. | -79.61 | 37.4456 | 22 | 1993-2018 | 193 | 416.5 | 1.4 | 0.643 |
| Pelee Island, ON, Canada. | -82.68 | 41.77 | 9 | 1997-2007 | 1810 | 292.5 | 20.6 | 0.156 |
| Petroglyphs, ON, Canada. | -78.1566 | 44.5881 | 16 | 1998-2015 | 2796 | 596.5 | 8.0 | 0.473 |
| Pettigrew State Park, NC. | -76.42 | 35.83 | 13 | 1995-2016 | 57 | 149.5 | 6.8 | 0.141 |
| PiedmontRum Creek, GA. | -83.7471 | 33.0937 | 5 | 1997-2014 | 6 | 130.25 | -0.3 | 0.919 |
| Pinery Provincial Park, ON, Canada. | -81.8921 | 43.2307 | 21 | 1994-2015 | 681 | 1252 | -9.2 | 0.013 |
| Pinnacles National Monument, CA. | -121.1978 | 36.4647 | 14 | 2000-2014 | 66 | 405.08 | -2.6 | 0.498 |
| Pioneers Park Nature Center, NE. | -96.7966 | 40.7701 | 16 | 2002-2018 | 314 | 94.25 | 26.8 | 0.193 |
| Point Pelee, ON, Canada. | -82.0167 | 42.35 | 9 | 1996-2008 | 3723 | 505.25 | -7.6 | 0.053 |
| Pomona, CA. | -117.72 | 34.12 | 7 | 1995-2005 | 22 | 113.75 | 4.7 | 0.267 |
| Pontotoc Ridge, OK. | -96.6081 | 34.5127 | 20 | 1996-2018 | 141 | 182.5 | 3.5 | 0.268 |
| Portal, AZ. | -109.1667 | 31.85 | 14 | 1999-2018 | 95 | 386 | -5.0 | 0.165 |
| Powdermill Nature Reserve, PA. | -79.2718 | 40.1602 | 7 | 2000-2010 | 226 | 140.5 | 26.1 | 0.240 |
| Powell Gardens, MO. | -94.03 | 38.9441 | 14 | 2003-2016 | 555 | 264.5 | -10.3 | 0.056 |
| Prince Georges Co. | -76.8257 | 38.7354 | 18 | 1993-2017 | 313 | 464.25 | -11.7 | 0.014 |
| Providence Co. | -71.5167 | 41.8667 | 12 | 2004-2018 | 167 | 190 | 16.2 | 0.338 |
| Putnam Co. East | -85.3167 | 36.1333 | 12 | 2003-2016 | 61 | 188.75 | 7.9 | 0.371 |
| Putnam Co. West | -85.6583 | 36.1545 | 8 | 2003-2011 | 37 | 75.5 | 5.3 | 0.206 |
| Quail Hollow, Hartville, OH. | -81.3291 | 40.8898 | 17 | 2000-2018 | 389 | 204.5 | 9.4 | 0.008 |
| Ramsey Canyon, AZ. | -110.2238 | 31.424 | 21 | 1995-2018 | 265 | 1150.5 | -1.9 | 0.694 |
| Ramsey Co. | -93.1667 | 45.0903 | 20 | 1995-2018 | 266 | 107.5 | 1.3 | 0.826 |
| Raritan Canal, NJ. | -74.5557 | 40.4297 | 24 | 1993-2018 | 1006 | 640.25 | 5.7 | 0.132 |
| Raven Rock State Park, NC. | -78.82 | 35.41 | 5 | 1997-2006 | 15 | 75.75 | 4.9 | 0.271 |
| Raven, TX. | -95.7167 | 30.4958 | 10 | 2006-2017 | 19 | 85 | -2.4 | 0.658 |
| Red Tail Farm, IN. | -87.118 | 39.798 | 24 | 1995-2018 | 119 | 141.5 | -1.9 | 0.632 |
| Reelfoot Lake, TN. | -89.37 | 36.43 | 14 | 2005-2018 | 60 | 119 | -3.8 | 0.291 |
| Regina, SK, Canada. | -104.6167 | 50.45 | 8 | 1997-2007 | 58 | 84.8 | 16.1 | 0.439 |
| Reston, VA. | -77.3372 | 38.953 | 5 | 1998-2018 | 20 | 35 | -0.1 | 0.980 |
| Retzer Nature Center, WI. | -88.3 | 43 | 9 | 1994-2005 | 220 | 43.5 | -19.5 | 0.045 |
| Rick Evans Grandview Prairie WMA | -93.7676 | 33.7889 | 10 | 2004-2018 | 98 | 172 | -5.7 | 0.445 |
| Riveredge Nature Center, WI. | -87.9793 | 43.3959 | 14 | 1993-2013 | 424 | 213.1 | -1.9 | 0.828 |
| Riverlands, PA. | -76.1369 | 41.1023 | 18 | 1994-2017 | 108 | 231 | 5.2 | 0.177 |
| Roan Mountain | -82.1105 | 36.1067 | 24 | 1993-2018 | 116 | 177.5 | 5.3 | 0.001 |
| Roanoke, VA. | -79.93 | 37.3 | 9 | 1996-2007 | 53 | 170 | 6.8 | 0.738 |
| Rochester, MN. | -92.4242 | 44.0323 | 20 | 1999-2018 | 1919 | 328.9 | 4.3 | 0.260 |
| Rochester, NY. | -77.4975 | 43.0887 | 21 | 1998-2018 | 287 | 291.95 | 0.1 | 0.987 |
| Rockland Co. | -74.0531 | 41.132 | 14 | 1995-2014 | 147 | 161.5 | 0.4 | 0.931 |
| Rogue River, MI | -85.6113 | 43.2629 | 19 | 1994-2015 | 90 | 155.75 | 1.1 | 0.773 |
| Roland, GA. | -84.4157 | 32.8594 | 5 | 2009-2018 | 12 | 104.5 | 12.0 | 0.647 |
| Rondeau Provincial Park, ON, Canada. | -81.9284 | 42.3015 | 14 | 1999-2015 | 2091 | 626 | -7.6 | 0.053 |
| Rowe Sanctuary, NE. | -98.88 | 40.62 | 11 | 1993-2005 | 57 | 70 | -0.9 | 0.844 |
| Salisbury, CT. | -73.4 | 42 | 11 | 1994-2007 | 311 | 470.75 | 14.5 | 0.300 |
| San Francisco, CA. | -122.4385 | 37.7643 | 6 | 2009-2018 | 14 | 197.5 | 6.0 | 0.711 |
| San Joaquin Co. | -121.3018 | 38.1544 | 18 | 1993-2015 | 75 | 122.5 | 0.7 | 0.944 |
| San Juan Capistrano, CA. | -117.65 | 33.48 | 7 | 1994-2005 | 50 | 44.5 | -0.5 | 0.976 |
| SandhillSeneca, WI. | -90.1126 | 44.3656 | 26 | 1993-2018 | 510 | 202.12 | 0.8 | 0.728 |
| Sandusky Bay | -82.9333 | 41.4167 | 15 | 1997-2015 | 203 | 109.25 | -15.0 | 0.335 |
| Sandy Creek, PA. | -80.036 | 41.3183 | 15 | 2003-2018 | 169 | 131.75 | -7.0 | 0.311 |
| Santa Ynez Canyon | -118.6044 | 34.0896 | 8 | 1994-2001 | 35 | 97.5 | 3.0 | 0.932 |
| Saskatoon East | -106.5274 | 52.1368 | 7 | 1997-2013 | 30 | 144.9 | 12.0 | 0.011 |
| Sentinel Rock State Park, VT. | -71.9481 | 44.8286 | 8 | 2001-2010 | 55 | 53 | -13.6 | 0.617 |
| Seven Ponds Nature Center, MI. | -83.2428 | 43.0014 | 9 | 2002-2018 | 60 | 56.5 | 16.5 | 0.022 |
| Severn Township, ON, Canada. | -79.57 | 44.73 | 8 | 1994-2003 | 115 | 59.25 | -2.7 | 0.778 |
| Shaw Nature Reserve | -90.8192 | 38.4668 | 13 | 1997-2018 | 86 | 141.3 | 3.0 | 0.635 |
| Shawnee Hills, IL. | -88.58 | 37.6 | 12 | 1993-2007 | 74 | 111.25 | -7.0 | 0.205 |
| Shawnee State Park and Forest, OH. | -83.1354 | 38.7122 | 23 | 1993-2018 | 179 | 638 | -2.1 | 0.585 |
| Shenandoah National Park, VA. | -78.4583 | 38.5915 | 21 | 1997-2018 | 1361 | 1261.5 | -2.0 | 0.683 |
| Shiawassee National Wildlife Refuge | -83.999 | 43.3579 | 9 | 2007-2016 | 717 | 226.5 | -29.6 | 0.040 |
| Shimek State Forest, IA. | -91.6866 | 40.5884 | 16 | 2001-2018 | 159 | 141 | 6.7 | 0.274 |
| Shreveport, LA. | -93.7138 | 32.3692 | 15 | 2001-2016 | 74 | 525.25 | -8.6 | 0.346 |
| Sister Bay, WI. | -87.1047 | 45.1783 | 15 | 1999-2015 | 554 | 114.6 | -2.7 | 0.183 |
| Skunks Misery | -81.7927 | 42.7151 | 19 | 2000-2018 | 992 | 704.5 | 14.1 | 0.053 |
| Sky Meadows State ParkThompson WMA | -77.9638 | 39.0153 | 9 | 2009-2018 | 59 | 148 | -8.5 | 0.792 |
| Soddy-Daisy, TN. | -85.15 | 35.2933 | 23 | 1998-2018 | 93 | 268.25 | 3.5 | 0.503 |
| Sonoma Co. | -122.6833 | 38.3333 | 5 | 1996-2004 | 13 | 27.75 | 14.5 | 0.177 |
| South Lyon Area, MI. | -83.6591 | 42.5779 | 11 | 2002-2012 | 63 | 158.5 | 4.9 | 0.536 |
| South San Francisco Bay, CA. | -122.05 | 37.48 | 12 | 1996-2007 | 68 | 55.58 | 16.9 | 0.001 |
| Southern Berkshire Co. | -73.3333 | 42.15 | 24 | 1993-2018 | 837 | 663.08 | -7.7 | 0.021 |
| Southern Bucks Co. | -74.9003 | 40.1813 | 18 | 1994-2018 | 255 | 186 | 3.7 | 0.549 |
| Southern Carbon County, PA. | -75.78 | 40.87 | 8 | 1996-2006 | 54 | 53 | -15.8 | 0.394 |
| Southern Kettle Moraine, WI. | -88.53 | 42.88 | 7 | 1993-2002 | 342 | 42.7 | 7.4 | 0.609 |
| Southern Lake Norman, NC. | -80.869 | 35.4425 | 15 | 2001-2018 | 174 | 348 | 18.6 | 0.004 |
| Southern New Haven Co. | -72.9218 | 41.3658 | 12 | 1995-2018 | 42 | 54.5 | -1.6 | 0.682 |
| Southwest Washtenaw, MI. | -84.0389 | 42.1023 | 32 | 1996-2018 | 628 | 546 | 14.7 | 0.000 |
| Split OakMoss Park, FL. | -81.2321 | 28.3482 | 5 | 2002-2018 | 7 | 76.25 | 3.1 | 0.093 |
| Springdale, NJ. | -74.8191 | 41.0732 | 25 | 1993-2018 | 1928 | 1101.5 | 2.8 | 0.622 |
| Staten Island, NY. | -74.147 | 40.572 | 19 | 1993-2018 | 119 | 185.25 | -2.2 | 0.388 |
| Stevenson, AL. | -85.8231 | 34.8541 | 10 | 2003-2015 | 39 | 80.5 | 9.9 | 0.225 |
| Stillhouse Hollow, TX. | -97.6 | 30.9667 | 6 | 2006-2013 | 14 | 85.5 | 3.6 | 0.899 |
| Stormville, NY. | -73.7511 | 41.59 | 22 | 1993-2017 | 126 | 140.5 | -6.2 | 0.087 |
| Storrs, CT. | -72.2333 | 41.8 | 15 | 1993-2009 | 261 | 188.5 | 3.8 | 0.688 |
| Strandburg, SD. | -96.7607 | 45.041 | 17 | 1995-2012 | 179 | 75.3 | 8.3 | 0.325 |
| Stroudsburg, PA. | -75.2164 | 40.9897 | 14 | 1998-2013 | 180 | 114.5 | 7.4 | 0.483 |
| Sunderland, ON, Canada. | -79.1836 | 44.2567 | 19 | 1997-2015 | 2371 | 1441 | 0.8 | 0.890 |
| Surry Co., NC. | -80.5333 | 36.35 | 6 | 2009-2018 | 47 | 62 | 40.9 | 0.087 |
| Tallahassee | -84.3045 | 30.449 | 6 | 2005-2017 | 8 | 160 | -1.7 | 0.365 |
| Tallgrass Prairie Preserve, OK. | -96.4126 | 36.8323 | 19 | 1993-2011 | 969 | 144.25 | 8.0 | 0.014 |
| Tampa, FL. | -82.35 | 28.05 | 11 | 2002-2017 | 50 | 126.5 | 13.0 | 0.051 |
| Tarrant Co. | -97.4804 | 32.7247 | 19 | 1994-2015 | 75 | 424 | -2.6 | 0.513 |
| Tennessee River Gorge, TN. | -85.3904 | 35.1244 | 20 | 1997-2018 | 68 | 366.75 | 1.6 | 0.683 |
| Tewaukon Lake, ND. | -97.4587 | 45.9943 | 12 | 1999-2011 | 103 | 57.2 | -7.6 | 0.146 |
| Tishomingo National Wildlife Refuge, OK. | -96.6446 | 34.1925 | 6 | 2006-2013 | 24 | 36 | -10.1 | 0.371 |
| Toronto (Centre), ON, Canada. | -79.4524 | 43.6563 | 21 | 1995-2015 | 2162 | 1093 | -1.0 | 0.800 |
| Toronto (Toronto Entomologists Assoc.), ON, Canada. | -79.25 | 43.78 | 13 | 1994-2007 | 506 | 293.5 | 14.6 | 0.179 |
| Towners Woods, OH. | -81.2717 | 41.112 | 18 | 2000-2018 | 279 | 215.75 | 5.2 | 0.180 |
| Transylvania Co. | -82.7369 | 35.2877 | 9 | 1996-2012 | 33 | 104.5 | 4.6 | 0.496 |
| Trapper Creek, ID. | -113.98 | 42.1678 | 5 | 2006-2012 | 8 | 142 | 6.8 | 0.498 |
| Trempealeau, WI. | -91.4112 | 44.0701 | 25 | 1993-2018 | 565 | 292.7 | -0.2 | 0.932 |
| Truro, MA. | -70.1167 | 42.0167 | 10 | 2004-2014 | 127 | 67 | -7.8 | 0.608 |
| Tulsa, OK. | -95.9224 | 36.2063 | 13 | 2001-2018 | 100 | 256.5 | -1.0 | 0.859 |
| Twinsburg, OH. | -81.5 | 41.26 | 11 | 2000-2018 | 117 | 125.5 | 4.3 | 0.175 |
| Utica, NY. | -75.3263 | 43.0828 | 16 | 1993-2012 | 61 | 158.3 | 6.0 | 0.172 |
| Victoria, TX. | -97.8 | 28.85 | 9 | 1998-2009 | 29 | 149.5 | -0.1 | 0.995 |
| Vischer Ferry, NY. | -73.833 | 42.7178 | 17 | 1993-2018 | 80 | 211.75 | -1.9 | 0.383 |
| Wake Co. | -78.6599 | 35.8563 | 18 | 1999-2018 | 198 | 303.5 | 12.7 | 0.008 |
| Wallkill River National Wildlife Refuge, NJ. | -74.5643 | 41.2007 | 18 | 2000-2018 | 751 | 327.5 | 2.5 | 0.674 |
| Washington Co., RI. | -71.7 | 41.4333 | 9 | 2006-2015 | 711 | 166 | -39.5 | 0.250 |
| Washington, DC. | -76.9834 | 38.9001 | 15 | 1996-2016 | 94 | 120.5 | -9.3 | 0.412 |
| Waterford, VA. | -77.65 | 39.2 | 20 | 1997-2018 | 1506 | 591 | 7.4 | 0.018 |
| Waubonsie, IA. | -95.7401 | 40.7662 | 25 | 1993-2018 | 506 | 171.55 | 0.8 | 0.794 |
| Wautoma, WI. | -89.25 | 44.07 | 5 | 1997-2004 | 45 | 33.4 | 9.1 | 0.016 |
| Wazee, WI. | -90.6597 | 44.277 | 24 | 1994-2018 | 335 | 144.47 | 3.9 | 0.098 |
| Wehr Nature Center, WI. | -87.9791 | 42.9304 | 20 | 1993-2017 | 244 | 116.5 | -11.5 | 0.000 |
| Wekiva River, FL. | -81.3667 | 28.7833 | 13 | 1996-2017 | 47 | 341 | 5.7 | 0.088 |
| West Anne Arundel Co. | -76.6848 | 39.0783 | 12 | 1999-2013 | 36 | 143.5 | -6.3 | 0.454 |
| West ReddingFairfield Co. | -73.4489 | 41.2958 | 22 | 1995-2018 | 264 | 176.25 | -1.9 | 0.661 |
| West Rutland, VT. | -73.0379 | 43.5941 | 15 | 1994-2012 | 250 | 115.5 | 8.1 | 0.043 |
| Western Garrett Co. | -79.4833 | 39.4833 | 6 | 2001-2008 | 66 | 46 | 70.8 | 0.001 |
| Western Hamilton Co. Parks | -84.7306 | 39.188 | 26 | 1993-2018 | 714 | 356.6 | -4.9 | 0.115 |
| Western Montgomery Co. | -77.3667 | 39.1667 | 23 | 1993-2018 | 178 | 878.8 | -2.2 | 0.428 |
| Western Niagara Co. | -78.9154 | 43.2127 | 25 | 1993-2018 | 748 | 1093.16 | -3.8 | 0.377 |
| WestportFairfield, CT. | -73.32 | 41.17 | 8 | 1994-2002 | 60 | 32.25 | -13.1 | 0.387 |
| Whitefish Point, MI. | -85.1347 | 46.6799 | 16 | 1996-2018 | 90 | 109 | -1.8 | 0.702 |
| Whitney Point, NY. | -75.8833 | 42.3333 | 13 | 1998-2015 | 65 | 64.75 | 1.4 | 0.871 |
| Wichita Mountains Wildlife Refuge, OK. | -98.6659 | 34.742 | 21 | 1993-2015 | 217 | 761.4 | 1.9 | 0.521 |
| Willow Slough | -121.7164 | 38.5827 | 18 | 1993-2017 | 268 | 95.24 | -4.4 | 0.475 |
| Wilmington, NC. | -77.8833 | 34.2667 | 10 | 1997-2012 | 213 | 120 | -18.8 | 0.015 |
| Windsor, NY. | -75.6167 | 42.1 | 13 | 1995-2009 | 71 | 71.25 | -8.2 | 0.152 |
| Windsor, ON, Canada. | -82.9638 | 42.2368 | 20 | 1994-2015 | 629 | 557.25 | -3.1 | 0.294 |
| Wolf Creek, OH. | -81.7756 | 41.2083 | 16 | 2001-2018 | 634 | 140 | 9.6 | 0.217 |
| Yankton, SD. | -97.3702 | 42.9959 | 19 | 1993-2015 | 257 | 69.8 | 3.8 | 0.608 |
| Yellow River State Forest, IA. | -91.2109 | 43.1272 | 18 | 2001-2018 | 431 | 150.95 | 7.5 | 0.296 |
| Yuba Pass, CA. | -120.5519 | 39.6619 | 15 | 2001-2018 | 39 | 439.75 | 2.7 | 0.367 |

**Supplementary Table 2.** **Covariate effects on summer-breeding monarch abundance trends based on model averaging of the AIC_c_-best generalized linear models.** Precipitation and temperature refer to the average cumulative precipitation and average annual temperature, respectively, in each count circle between 1993-2018. Precipitation trend and temperature trend refers to the trend in precipitation and temperature (standard deviations per year), respectively, between 1901-2018. Cropland refers to the proportion cropland in each count circle between 2008-2018. Glyphosate use refers to the areal-weighted number of pounds of glyphosate applied in counties overlapping with each count circle, averaged from 1993-2017. Road density refers to the proportion of area within each count circle that is covered by roads.

| Covariate | Estimate | Standard error | 2.5% | 97.5% |
| --- | --- | --- | --- | --- |
| Intercept | 0.016089 | 0.006436 | 0.003422 | 0.028755 |
| Temperature | 0.013451 | 0.009613 | -0.00543 | 0.032336 |
| Precipitation | -0.01081 | 0.008271 | -0.02706 | 0.005451 |
| Precipitation trend | -0.01715 | 0.006996 | -0.03091 | -0.00338 |
| Temperature trend | -0.01081 | 0.006773 | -0.02414 | 0.002515 |
| Cropland | -0.00869 | 0.007231 | -0.02291 | 0.005541 |
| Glyphosate use | -0.0077 | 0.006953 | -0.02138 | 0.005985 |
| Road density | -0.00283 | 0.00647 | -0.01556 | 0.009904 |

**Supplementary Table 3.** **Covariate effects on summer-breeding monarch abundance trends based on model averaging of the AIC_c_-best generalized linear models.** Precipitation and temperature refer to the average cumulative precipitation and average annual temperature, respectively, in each count circle between 1993-2018. Cropland refers to the proportion cropland in each count circle between 2008-2018. Glyphosate use refers to the areal-weighted number of pounds of glyphosate applied in counties overlapping with each count circle, averaged from 1993-2017. Road density refers to the proportion of area within each count circle that is covered by roads.

| Covariate | Estimate | Standard error | 2.5% | 97.5% |
| --- | --- | --- | --- | --- |
| Intercept | 0.01609 | 0.00645 | 0.00339 | 0.02879 |
| Precipitation | -0.0129 | 0.00706 | -0.0268 | 0.00101 |
| Temperature | 0.0187 | 0.00705 | 0.00483 | 0.03256 |
| Cropland | -0.0072 | 0.00668 | -0.0204 | 0.00594 |
| Glyphosate use | -0.0085 | 0.00675 | -0.0218 | 0.00481 |
| Road density | -0.0039 | 0.00646 | -0.0166 | 0.00883 |

**Supplementary Table 4.** **Model selection results from models that included three land use and four climate covariates.** Covariate effects are given in Supplementary Table 2.

| Model | AICc | delta | weight | Precip. | Temp. | Precip. trend | Temp. trend | Cropland | Glyphosate use | Road density |
| --- | --- | --- | --- | --- | --- | --- | --- | --- | --- | --- |
| 1 | -459.107 | 0 | 0.04302 |  |  | + | + |  |  |  |
| 2 | -458.867 | 0.239731 | 0.038161 |  |  | + |  |  |  |  |
| 3 | -458.855 | 0.251916 | 0.037929 |  |  | + | + | + |  |  |
| 4 | -458.55 | 0.556701 | 0.032568 |  |  | + | + |  | + |  |
| 5 | -457.921 | 1.18615 | 0.023774 |  | + | + |  |  |  |  |
| 6 | -457.816 | 1.291142 | 0.022558 |  | + | + | + |  |  |  |
| 7 | -457.771 | 1.335595 | 0.022062 | + | + |  |  |  |  |  |
| 8 | -457.656 | 1.450994 | 0.020825 |  |  | + |  |  | + |  |
| 9 | -457.581 | 1.526282 | 0.020056 | + | + |  | + | + |  |  |
| 10 | -457.481 | 1.626385 | 0.019077 |  |  | + |  | + |  |  |
| 11 | -457.363 | 1.744039 | 0.017987 | + | + |  | + |  | + |  |
| 12 | -457.307 | 1.800237 | 0.017489 | + |  | + | + | + |  |  |
| 13 | -457.275 | 1.832147 | 0.017212 | + | + |  |  |  | + |  |
| 14 | -457.254 | 1.853333 | 0.01703 |  |  | + | + | + | + |  |
| 15 | -457.231 | 1.876292 | 0.016836 |  |  | + | + |  |  | + |
| 16 | -457.193 | 1.913988 | 0.016522 |  | + | + | + | + |  |  |
| 17 | -457.182 | 1.925056 | 0.016431 |  | + | + | + |  | + |  |
| 18 | -457.176 | 1.931319 | 0.016379 | + | + |  | + |  |  |  |
| 19 | -457.137 | 1.969887 | 0.016066 | + |  | + | + |  |  |  |
| 20 | -457.069 | 2.038172 | 0.015527 |  |  | + | + | + |  | + |
| 21 | -457.01 | 2.097011 | 0.015077 |  |  | + |  |  |  | + |
| 22 | -456.896 | 2.210765 | 0.014243 | + |  | + |  |  |  |  |
| 23 | -456.868 | 2.239548 | 0.01404 | + | + | + |  |  |  |  |
| 24 | -456.867 | 2.240109 | 0.014036 | + | + |  |  | + |  |  |
| 25 | -456.861 | 2.246148 | 0.013994 |  | + |  |  |  |  |  |
| 26 | -456.859 | 2.248353 | 0.013978 | + |  | + | + |  | + |  |
| 27 | -456.764 | 2.342752 | 0.013334 | + | + | + | + |  | + |  |
| 28 | -456.698 | 2.40881 | 0.0129 |  |  | + | + |  | + | + |
| 29 | -456.682 | 2.424789 | 0.012798 |  | + | + |  |  | + |  |
| 30 | -456.636 | 2.470598 | 0.012508 | + | + | + | + | + |  |  |
| 31 | -456.602 | 2.505457 | 0.012292 | + | + | + | + |  |  |  |
| 32 | -456.351 | 2.756561 | 0.010842 | + | + |  | + | + | + |  |
| 33 | -456.317 | 2.79047 | 0.010659 |  | + | + |  | + |  |  |
| 34 | -456.293 | 2.814494 | 0.010532 | + | + | + |  |  | + |  |
| 35 | -456.279 | 2.827963 | 0.010461 |  | + |  | + |  |  |  |
| 36 | -456.071 | 3.036344 | 0.009426 |  | + | + |  |  |  | + |
| 37 | -456.067 | 3.040094 | 0.009409 | + | + |  |  |  |  | + |
| 38 | -456.015 | 3.091722 | 0.009169 | + | + |  | + | + |  | + |
| 39 | -455.943 | 3.163657 | 0.008845 |  | + | + | + |  |  | + |
| 40 | -455.853 | 3.254308 | 0.008453 | + |  | + |  |  | + |  |
| 41 | -455.839 | 3.268136 | 0.008395 | + |  | + | + | + | + |  |
| 42 | -455.814 | 3.293364 | 0.008289 |  |  | + |  |  | + | + |
| 43 | -455.771 | 3.336432 | 0.008113 |  |  | + |  | + | + |  |
| 44 | -455.721 | 3.385797 | 0.007915 | + | + |  | + |  | + | + |
| 45 | -455.69 | 3.417342 | 0.007791 | + |  | + |  | + |  |  |
| 46 | -455.674 | 3.433015 | 0.00773 | + | + | + |  | + |  |  |
| 47 | -455.669 | 3.437994 | 0.007711 |  |  | + |  | + |  | + |
| 48 | -455.634 | 3.472604 | 0.007579 |  | + | + | + | + | + |  |
| 49 | -455.628 | 3.479323 | 0.007553 | + | + |  |  |  | + | + |
| 50 | -455.622 | 3.484933 | 0.007532 |  | + |  | + | + |  |  |
| 51 | -455.547 | 3.559712 | 0.007256 | + |  | + | + | + |  | + |
| 52 | -455.524 | 3.583147 | 0.007171 | + | + |  |  | + | + |  |
| 53 | -455.484 | 3.622848 | 0.00703 | + | + | + | + | + | + |  |
| 54 | -455.459 | 3.647665 | 0.006944 | + | + |  | + |  |  | + |
| 55 | -455.452 | 3.654949 | 0.006918 |  |  | + | + | + | + | + |
| 56 | -455.399 | 3.7083 | 0.006736 |  | + | + | + | + |  | + |
| 57 | -455.334 | 3.773534 | 0.00652 |  | + | + | + |  | + | + |
| 58 | -455.329 | 3.77819 | 0.006505 |  | + |  |  | + |  |  |
| 59 | -455.294 | 3.813472 | 0.006391 |  | + |  |  |  | + |  |
| 60 | -455.254 | 3.853224 | 0.006265 | + |  | + | + |  |  | + |
| 61 | -455.251 | 3.856186 | 0.006256 | + | + |  |  | + |  | + |
| 62 | -455.168 | 3.939106 | 0.006002 |  | + |  |  |  |  | + |
| 63 | -455.074 | 4.032581 | 0.005728 |  | + |  | + |  | + |  |
| 64 | -455.046 | 4.061247 | 0.005647 | + | + | + |  |  |  | + |
| 65 | -455.032 | 4.075387 | 0.005607 | + |  | + |  |  |  | + |
| 66 | -455.013 | 4.093806 | 0.005555 | + |  | + | + |  | + | + |
| 67 | -454.977 | 4.130432 | 0.005455 | + | + | + | + |  | + | + |
| 68 | -454.92 | 4.187238 | 0.005302 | + | + | + | + | + |  | + |
| 69 | -454.848 | 4.259151 | 0.005115 |  | + | + |  |  | + | + |
| 70 | -454.791 | 4.316118 | 0.004971 | + | + |  | + | + | + | + |
| 71 | -454.754 | 4.353246 | 0.00488 | + | + | + | + |  |  | + |
| 72 | -454.675 | 4.432018 | 0.004691 |  | + | + |  | + | + |  |
| 73 | -454.574 | 4.532644 | 0.004461 |  | + |  | + |  |  | + |
| 74 | -454.519 | 4.587581 | 0.00434 | + | + | + |  |  | + | + |
| 75 | -454.502 | 4.604587 | 0.004303 |  | + | + |  | + |  | + |
| 76 | -454.386 | 4.721208 | 0.00406 | + | + | + |  | + | + |  |
| 77 | -454.069 | 5.037648 | 0.003465 | + |  | + | + | + | + | + |
| 78 | -454.052 | 5.054737 | 0.003436 | + |  | + |  | + | + |  |
| 79 | -454.022 | 5.085121 | 0.003384 |  | + |  | + | + |  | + |
| 80 | -454.013 | 5.093827 | 0.003369 | + |  | + |  |  | + | + |
| 81 | -453.943 | 5.16361 | 0.003254 |  |  | + |  | + | + | + |
| 82 | -453.929 | 5.178061 | 0.003231 | + | + | + |  | + |  | + |
| 83 | -453.91 | 5.197298 | 0.0032 | + | + |  |  | + | + | + |
| 84 | -453.89 | 5.216969 | 0.003168 | + |  | + |  | + |  | + |
| 85 | -453.886 | 5.221322 | 0.003161 |  |  |  | + | + |  |  |
| 86 | -453.825 | 5.28211 | 0.003067 |  | + | + | + | + | + | + |
| 87 | -453.766 | 5.340944 | 0.002978 | + | + | + | + | + | + | + |
| 88 | -453.733 | 5.374458 | 0.002928 |  | + |  | + | + | + |  |
| 89 | -453.688 | 5.418905 | 0.002864 |  | + |  |  | + |  | + |
| 90 | -453.624 | 5.483178 | 0.002773 |  | + |  |  |  | + | + |
| 91 | -453.57 | 5.536705 | 0.0027 | + |  |  | + | + |  |  |
| 92 | -453.421 | 5.685801 | 0.002506 |  | + |  |  | + | + |  |
| 93 | -453.419 | 5.688282 | 0.002503 |  |  |  |  |  |  |  |
| 94 | -453.407 | 5.700405 | 0.002488 |  | + |  | + |  | + | + |
| 95 | -453.293 | 5.813635 | 0.002351 |  |  |  | + |  |  |  |
| 96 | -452.846 | 6.261539 | 0.001879 |  | + | + |  | + | + | + |
| 97 | -452.637 | 6.470271 | 0.001693 | + | + | + |  | + | + | + |
| 98 | -452.626 | 6.480826 | 0.001684 |  |  |  |  | + |  |  |
| 99 | -452.406 | 6.700647 | 0.001509 |  |  |  | + | + |  | + |
| 100 | -452.241 | 6.866379 | 0.001389 | + |  | + |  | + | + | + |
| 101 | -452.135 | 6.971929 | 0.001317 | + |  |  | + | + |  | + |
| 102 | -452.125 | 6.982382 | 0.001311 |  | + |  | + | + | + | + |
| 103 | -452.05 | 7.056881 | 0.001263 | + |  |  |  |  |  |  |
| 104 | -451.97 | 7.136833 | 0.001213 | + |  |  | + |  |  |  |
| 105 | -451.897 | 7.210137 | 0.00117 |  |  |  | + |  | + |  |
| 106 | -451.833 | 7.274481 | 0.001133 |  |  |  | + | + | + |  |
| 107 | -451.826 | 7.281269 | 0.001129 | + |  |  |  | + |  |  |
| 108 | -451.806 | 7.301436 | 0.001117 |  |  |  |  |  |  | + |
| 109 | -451.772 | 7.335342 | 0.001099 |  | + |  |  | + | + | + |
| 110 | -451.675 | 7.431723 | 0.001047 |  |  |  |  |  | + |  |
| 111 | -451.666 | 7.44096 | 0.001042 |  |  |  | + |  |  | + |
| 112 | -451.611 | 7.49628 | 0.001014 | + |  |  | + | + | + |  |
| 113 | -451.103 | 8.004189 | 0.000786 |  |  |  |  | + |  | + |
| 114 | -451.048 | 8.059403 | 0.000765 | + |  |  | + |  | + |  |
| 115 | -450.598 | 8.509425 | 0.000611 | + |  |  |  |  | + |  |
| 116 | -450.574 | 8.533283 | 0.000604 |  |  |  |  | + | + |  |
| 117 | -450.433 | 8.674238 | 0.000562 | + |  |  |  |  |  | + |
| 118 | -450.343 | 8.764574 | 0.000538 |  |  |  | + | + | + | + |
| 119 | -450.339 | 8.768102 | 0.000537 | + |  |  | + |  |  | + |
| 120 | -450.331 | 8.776128 | 0.000535 | + |  |  |  | + |  | + |
| 121 | -450.305 | 8.80206 | 0.000528 |  |  |  | + |  | + | + |
| 122 | -450.174 | 8.933308 | 0.000494 | + |  |  | + | + | + | + |
| 123 | -450.081 | 9.025991 | 0.000472 |  |  |  |  |  | + | + |
| 124 | -449.807 | 9.300303 | 0.000411 | + |  |  |  | + | + |  |
| 125 | -449.475 | 9.632515 | 0.000348 | + |  |  | + |  | + | + |
| 126 | -449.038 | 10.06951 | 0.00028 |  |  |  |  | + | + | + |
| 127 | -449.017 | 10.0896 | 0.000277 | + |  |  |  |  | + | + |
| 128 | -448.305 | 10.8023 | 0.000194 | + |  |  |  | + | + | + |

**Supplementary Table 5.** **Model selection results from models that included three land use and two climate covariates.** Covariate effects are given in Supplementary Table 3.

| Model | AICc | delta | weight | Cropland | Glyphosate use | Road density | Precipitation | Temperature |
| --- | --- | --- | --- | --- | --- | --- | --- | --- |
| 1 | -457.771 | 0 | 0.154815 |  |  |  | + | + |
| 2 | -457.275 | 0.496552 | 0.120778 |  | + |  | + | + |
| 3 | -456.867 | 0.904514 | 0.098492 | + |  |  | + | + |
| 4 | -456.861 | 0.910553 | 0.098195 |  |  |  |  | + |
| 5 | -456.067 | 1.704499 | 0.066021 |  |  | + | + | + |
| 6 | -455.628 | 2.143728 | 0.053004 |  | + | + | + | + |
| 7 | -455.524 | 2.247552 | 0.050323 | + | + |  | + | + |
| 8 | -455.329 | 2.442595 | 0.045647 | + |  |  |  | + |
| 9 | -455.294 | 2.477877 | 0.044849 |  | + |  |  | + |
| 10 | -455.251 | 2.520591 | 0.043901 | + |  | + | + | + |
| 11 | -455.168 | 2.603511 | 0.042118 |  |  | + |  | + |
| 12 | -453.91 | 3.861703 | 0.022452 | + | + | + | + | + |
| 13 | -453.688 | 4.08331 | 0.020097 | + |  | + |  | + |
| 14 | -453.624 | 4.147583 | 0.019461 |  | + | + |  | + |
| 15 | -453.421 | 4.350206 | 0.017586 | + | + |  |  | + |
| 16 | -453.419 | 4.352687 | 0.017565 |  |  |  |  |  |
| 17 | -452.626 | 5.145231 | 0.011818 | + |  |  |  |  |
| 18 | -452.05 | 5.721285 | 0.00886 |  |  |  | + |  |
| 19 | -451.826 | 5.945674 | 0.00792 | + |  |  | + |  |
| 20 | -451.806 | 5.965841 | 0.007841 |  |  | + |  |  |
| 21 | -451.772 | 5.999747 | 0.007709 | + | + | + |  | + |
| 22 | -451.675 | 6.096128 | 0.007346 |  | + |  |  |  |
| 23 | -451.103 | 6.668594 | 0.005518 | + |  | + |  |  |
| 24 | -450.598 | 7.17383 | 0.004286 |  | + |  | + |  |
| 25 | -450.574 | 7.197688 | 0.004235 | + | + |  |  |  |
| 26 | -450.433 | 7.338643 | 0.003947 |  |  | + | + |  |
| 27 | -450.331 | 7.440533 | 0.003751 | + |  | + | + |  |
| 28 | -450.081 | 7.690396 | 0.00331 |  | + | + |  |  |
| 29 | -449.807 | 7.964707 | 0.002886 | + | + |  | + |  |
| 30 | -449.038 | 8.733919 | 0.001965 | + | + | + |  |  |
| 31 | -449.017 | 8.754006 | 0.001945 |  | + | + | + |  |
| 32 | -448.305 | 9.466707 | 0.001362 | + | + | + | + |  |

**Supplementary Table 6. Overall trends among 456 North American butterfly species between 1993-2018, summarized from Crossley et al. (2021).** Median trend represents the median change in abundance of each butterfly species across the 50x50 km grid cells occupied, and is interpreted as percent change per year per hour of sampling effort. Lower and Upper represent the 95% confidence intervals around the median trend estimate. Total no. butterflies represents the total number of butterflies observed over the sampling period. No. sites represents the number of 50x50 grid cells in which a species occurred. No. years represents the number of years between the first and last year when a species was observed. Note that some extreme positive or negative trends may reflect poor sampling coverage.

| Species | Median trend | Lower | Upper | Total no. butterflies | No. sites | No. years | First year | Last year |
| --- | --- | --- | --- | --- | --- | --- | --- | --- |
| *Chlosyne whitneyi* | -60.38 | -89.79 | 42.13 | 178 | 1 | 7 | 2000 | 2009 |
| *Calephelis rawsoni* | -41.14 | -69.9 | 15.02 | 12 | 1 | 6 | 2005 | 2013 |
| *Cogia calchas* | -31.16 | -41.13 | -19.42 | 66 | 1 | 6 | 2000 | 2008 |
| *Amblyscirtes reversa* | -22.3 | -33.11 | -10.6 | 136 | 2 | 14 | 1993 | 2014 |
| *Papilio aristodemus* | -19.98 | -44.1 | 14.24 | 19 | 1 | 5 | 2004 | 2016 |
| *Erebia theano* | -18.96 | -32.03 | -3.629 | 450 | 1 | 10 | 1994 | 2007 |
| *Hesperia leonardus* | -17.46 | -49.33 | 32.55 | 82 | 1 | 8 | 2011 | 2018 |
| *Ministrymon azia* | -16.31 | -39.33 | 15.62 | 11 | 1 | 6 | 2002 | 2009 |
| *Boloria chariclea* | -15.56 | -26.01 | -5.523 | 787 | 2 | 12 | 2001 | 2018 |
| *Pontia sisymbrii* | -15.39 | -30.81 | -0.65155 | 71 | 2 | 14 | 2002 | 2017 |
| *Amblyscirtes phylace* | -15.24 | -28.01 | 0.8697 | 33 | 2 | 11 | 1993 | 2009 |
| *Plebejus neurona* | -15.05 | -24.75 | -3.954 | 33 | 1 | 10 | 1997 | 2010 |
| *Hemiargus ammon* | -14.79 | -55.36 | 62.19 | 190 | 1 | 6 | 1999 | 2005 |
| *Anaea floridalis* | -14.56 | -37.94 | 17.64 | 17 | 1 | 7 | 1997 | 2005 |
| *Pyrgus xanthus* | -14.54 | -25.55 | -1.65 | 14 | 1 | 5 | 1993 | 2005 |
| *Erynnis meridianus* | -14.1 | -30.65 | 7.3 | 26 | 1 | 5 | 1998 | 2011 |
| *Hemiargus thomasi* | -13.83 | -40.79 | 24.92 | 369 | 1 | 11 | 1998 | 2009 |
| *Boloria freija* | -12.28 | -21.71 | -1.909 | 269 | 1 | 13 | 1998 | 2014 |
| *Ancyloxypha arene* | -12.21 | -19.2 | -4.303 | 96 | 1 | 13 | 1993 | 2012 |
| *Amblyscirtes carolina* | -11.293 | -28.02 | 10.705 | 389 | 2 | 12 | 1999 | 2014 |
| *Oeneis macounii* | -11.002 | -21.8 | 0.428 | 117 | 3 | 15 | 1993 | 2014 |
| *Calephelis arizonensis* | -10.717 | -17.54 | -0.547 | 398 | 3 | 17 | 1993 | 2010 |
| *Erebia callias* | -10.58 | -20.93 | 1.082 | 1394 | 1 | 10 | 1994 | 2007 |
| *Lycaena cupreus* | -10.424 | -15.96 | -4.6566 | 241 | 3 | 22 | 1993 | 2018 |
| *Chlorostrymon simaethis* | -9.328 | -19.97 | 2.61 | 37 | 1 | 9 | 2001 | 2016 |
| *Ministrymon leda* | -9.263 | -14.24 | -3.816 | 3577 | 5 | 25 | 1993 | 2018 |
| *Zestusa dorus* | -9.118 | -21.19 | 4.729 | 14 | 1 | 8 | 1997 | 2017 |
| *Hesperia lindseyi* | -8.965 | -18.21 | 0.4529 | 207 | 2 | 21 | 1993 | 2014 |
| *Erynnis brizo* | -8.687 | -35.34 | 28.94 | 7 | 1 | 5 | 2003 | 2011 |
| *Polites carus* | -8.541 | -15.22 | -1.1267 | 97 | 3 | 18 | 1993 | 2012 |
| *Celotes nessus* | -8.436 | -14.84 | -2.4802 | 556 | 7 | 23 | 1993 | 2018 |
| *Eurema nise* | -8.387 | -16.03 | -0.9688 | 205 | 3 | 17 | 1993 | 2016 |
| *Pyrrhopyge araxes* | -8.166 | -13.89 | -2.1318 | 3101 | 4 | 23 | 1993 | 2018 |
| *Hesperia uncas* | -8.019 | -14.78 | -0.8248 | 27 | 1 | 11 | 1993 | 2015 |
| *Atrytonopsis edwardsii* | -7.934 | -18.64 | 4.415 | 295 | 2 | 12 | 1996 | 2013 |
| *Vanessa annabella* | -7.789 | -11.68 | -3.746 | 1954 | 28 | 26 | 1993 | 2018 |
| *Euphyes pilatka* | -7.126 | -13.96 | -0.895 | 197 | 7 | 17 | 2002 | 2018 |
| *Cyllopsis pyracmon* | -7.076 | -13.07 | 0.2717 | 403 | 4 | 24 | 1993 | 2018 |
| *Piruna pirus* | -7.018 | -12.63 | -1.2003 | 1626 | 7 | 23 | 1993 | 2015 |
| *Erynnis telemachus* | -7.014 | -19.03 | 6.786 | 12 | 1 | 6 | 1994 | 2006 |
| *Cogia caicus* | -6.927 | -11.674 | -2.0052 | 439 | 2 | 25 | 1993 | 2018 |
| *Thorybes drusius* | -6.738 | -13.414 | 0.03633 | 1717 | 4 | 25 | 1993 | 2018 |
| *Colias gigantea* | -6.604 | -24.02 | 15.31 | 31 | 2 | 9 | 1994 | 2003 |
| *Thorybes mexicanus* | -6.325 | -13.933 | 0.6262 | 1023 | 7 | 26 | 1993 | 2018 |
| *Junonia genoveva* | -6.298 | -17.82 | 3.738 | 1874 | 5 | 21 | 1993 | 2015 |
| *Atrytonopsis lunus* | -6.273 | -10.489 | -1.5719 | 276 | 4 | 26 | 1993 | 2018 |
| *Apodemia palmeri* | -6.234 | -15.46 | 3.49623 | 1335 | 4 | 25 | 1993 | 2018 |
| *Oarisma edwardsii* | -5.946 | -11.064 | -0.4082 | 163 | 2 | 21 | 1993 | 2018 |
| *Poladryas arachne* | -5.938 | -10.448 | 4.015 | 423 | 3 | 23 | 1993 | 2015 |
| *Phyciodes batesii* | -5.8959 | -10.126 | -0.3133 | 605 | 15 | 24 | 1994 | 2018 |
| *Hesperia pahaska* | -5.838 | -9.974 | -1.782689 | 741 | 7 | 26 | 1993 | 2018 |
| *Systasea pulverulenta* | -5.712 | -17.87 | 7.66 | 40 | 2 | 14 | 1993 | 2016 |
| *Phyciodes texana* | -5.633 | -9.827 | -0.5928 | 1768 | 16 | 26 | 1993 | 2018 |
| *Phyciodes pallida* | -5.629 | -12.04 | 3.4008 | 247 | 5 | 24 | 1993 | 2016 |
| *Phyciodes vesta* | -5.557 | -14.548 | 1.4447 | 422 | 6 | 24 | 1993 | 2016 |
| *Cercyonis meadii* | -5.425 | -16.25 | 6.611 | 73 | 2 | 15 | 1993 | 2014 |
| *Thymelicus lineola* | -5.415 | -13.88 | 2.967 | 1219857 | 166 | 26 | 1993 | 2018 |
| *Colias scudderii* | -5.177 | -18.58 | 10.37 | 27 | 1 | 9 | 1997 | 2013 |
| *Speyeria adiaste* | -5.172 | -14.03 | 4.555 | 325 | 1 | 18 | 1998 | 2017 |
| *Lycaena dione* | -5.17 | -10.362 | 0.7929 | 951 | 20 | 26 | 1993 | 2018 |
| *Amblyscirtes belli* | -5.161 | -16.88 | 9.044 | 161 | 3 | 14 | 2001 | 2018 |
| *Satyrium auretorum* | -5.138 | -15.34 | 6.994 | 698 | 6 | 24 | 1993 | 2018 |
| *Heliopetes domicella* | -5.05 | -21.9 | 15.41 | 9 | 1 | 6 | 1994 | 2005 |
| *Asterocampa leilia* | -5.028 | -13.553 | 3.4024 | 2682 | 6 | 26 | 1993 | 2018 |
| *Habrodais grunus* | -5.001 | -12.44 | 3.8474 | 2942 | 10 | 24 | 1993 | 2017 |
| *Satyrium sylvinus* | -4.94 | -9.971 | 1.2322 | 2119 | 19 | 26 | 1993 | 2018 |
| *Euchloe ausonides* | -4.903 | -7.707 | -2.066 | 1240 | 16 | 26 | 1993 | 2018 |
| *Eurema mexicana* | -4.831 | -11.521 | 1.3417 | 4899 | 13 | 26 | 1993 | 2018 |
| *Phocides polybius* | -4.816 | -21.27 | 15.07 | 7 | 1 | 6 | 2004 | 2016 |
| *Lycaena dorcas* | -4.669 | -12.651 | 3.5716 | 1339 | 7 | 22 | 1997 | 2018 |
| *Erebia epipsodea* | -4.62575 | -10.172 | 2.265 | 6914 | 15 | 26 | 1993 | 2018 |
| *Megisto rubricata* | -4.397 | -10.547 | 3.222 | 2757 | 6 | 26 | 1993 | 2018 |
| *Achlyodes thraso* | -4.306 | -11.617 | 3.413 | 494 | 4 | 18 | 1993 | 2016 |
| *Codatractus arizonensis* | -4.301 | -9.979 | 1.965 | 753 | 3 | 26 | 1993 | 2018 |
| *Amblyscirtes hegon* | -4.245 | -12.991 | 3.7122 | 66 | 4 | 18 | 1996 | 2017 |
| *Hypaurotis crysalus* | -4.211 | -19.96 | 3.905 | 145 | 3 | 16 | 1993 | 2018 |
| *Boloria frigga* | -4.17 | -15.62 | 8.723 | 28 | 1 | 8 | 1998 | 2011 |
| *Lycaena xanthoides* | -4.1491 | -7.595 | 3.35 | 6370 | 12 | 26 | 1993 | 2018 |
| *Anthocharis lanceolata* | -4.12 | -14.88 | 6.231 | 156 | 3 | 18 | 1995 | 2017 |
| *Calephelis perditalis* | -4.108 | -13.26 | 6.371 | 186 | 2 | 15 | 1996 | 2014 |
| *Achalarus casica* | -4.072 | -9.967 | 0.43397 | 246 | 5 | 24 | 1993 | 2018 |
| *Chlosyne theona* | -4.068 | -8.632 | 1.7197 | 678 | 5 | 25 | 1993 | 2018 |
| *Hesperia viridis* | -4.065 | -11.112 | 3.899 | 298 | 3 | 21 | 1993 | 2015 |
| *Colias eurydice* | -4.058 | -9.552 | 2.0303 | 2294 | 8 | 26 | 1993 | 2018 |
| *Glaucopsyche piasus* | -4.052 | -9.314 | 1.6544 | 639 | 12 | 26 | 1993 | 2018 |
| *Erynnis afranius* | -4.0209 | -12.36 | 5.6558 | 141 | 6 | 20 | 1993 | 2013 |
| *Erynnis martialis* | -3.944 | -10.23 | 4.7399 | 214 | 4 | 26 | 1993 | 2018 |
| *Paramacera allyni* | -3.779 | -9.567 | 2.671 | 40 | 2 | 17 | 1993 | 2018 |
| *Amblyscirtes exoteria* | -3.745 | -7.899 | 0.6814 | 576 | 4 | 26 | 1993 | 2018 |
| *Lycaena rubidus* | -3.7428 | -10.461 | 4.971 | 807 | 9 | 26 | 1993 | 2018 |
| *Chlosyne harrisii* | -3.741 | -7.953 | 0.905 | 1754 | 24 | 26 | 1993 | 2018 |
| *Boloria montinus* | -3.7207 | -14.72 | 7.812 | 1258 | 2 | 15 | 1994 | 2009 |
| *Callophrys eryphon* | -3.72 | -7.911 | 1.0668 | 751 | 15 | 25 | 1993 | 2018 |
| *Nymphalis milberti* | -3.675 | -6.9 | -0.46906 | 3777 | 68 | 26 | 1993 | 2018 |
| *Papilio indra* | -3.64 | -10.088 | 3.0701 | 186 | 8 | 22 | 1993 | 2018 |
| *Speyeria diana* | -3.5917 | -8.898 | 3.444 | 1520 | 15 | 26 | 1993 | 2018 |
| *Callophrys augustinus* | -3.57 | -7.679 | 1.1289 | 1141 | 20 | 26 | 1993 | 2018 |
| *Eurema boisduvaliana* | -3.467 | -12.16 | 6.116 | 18 | 1 | 5 | 1993 | 2012 |
| *Apodemia mormo* | -3.441 | -10.123 | 2.995 | 874 | 7 | 22 | 1993 | 2016 |
| *Strymon bazochii* | -3.377 | -30.48 | 34.25 | 12 | 1 | 5 | 2004 | 2016 |
| *Calephelis virginiensis* | -3.359 | -9.749 | 4.432 | 1626 | 21 | 26 | 1993 | 2018 |
| *Copaeodes minimus* | -3.31543 | -9.12 | 3.89 | 10762 | 70 | 26 | 1993 | 2018 |
| *Erynnis scudderi* | -3.268 | -10.32 | 4.312 | 17 | 1 | 7 | 1993 | 2014 |
| *Satyrium californica* | -3.199 | -8.115 | 1.8996 | 3622 | 22 | 26 | 1993 | 2018 |
| *Everes amyntula* | -3.19318 | -8.755 | 1.9868 | 3422 | 26 | 26 | 1993 | 2018 |
| *Agriades glandon* | -3.189 | -7.845 | 2.239 | 3084 | 13 | 26 | 1993 | 2018 |
| *Euphydryas editha* | -3.186 | -10.375 | 4.549 | 4385 | 13 | 26 | 1993 | 2018 |
| *Chioides catillus* | -3.171 | -11.618 | 6.345 | 1266 | 10 | 23 | 1993 | 2018 |
| *Erynnis icelus* | -3.155 | -8.751 | 2.555 | 1495 | 21 | 26 | 1993 | 2018 |
| *Phyciodes selenis* | -3.152 | -7.881 | 1.929 | 124579 | 93 | 26 | 1993 | 2018 |
| *Chlosyne hoffmanni* | -3.1172 | -10.232 | 5.485 | 7772 | 6 | 26 | 1993 | 2018 |
| *Anteos clorinde* | -3.1 | -16.43 | 12.33 | 10 | 1 | 6 | 1993 | 2007 |
| *Polites sonora* | -3.0847 | -7.112 | 1.8999 | 7909 | 10 | 26 | 1993 | 2018 |
| *Kricogonia lyside* | -3.062 | -10.9 | 6.465 | 12723 | 16 | 26 | 1993 | 2018 |
| *Lycaena arota* | -3.0212 | -12.395 | 7.372 | 3369 | 12 | 26 | 1993 | 2018 |
| *Speyeria coronis* | -2.9892 | -8.531 | 2.831 | 4972 | 21 | 26 | 1993 | 2018 |
| *Plebejus saepiolus* | -2.986 | -8.659 | 3.112 | 35138 | 33 | 26 | 1993 | 2018 |
| *Phoebis neleis* | -2.962 | -29.8 | 34.02 | 32 | 1 | 7 | 2012 | 2018 |
| *Amblyscirtes oslari* | -2.9409 | -9.27 | 4.2006 | 67 | 2 | 15 | 1993 | 2018 |
| *Adelpha bredowii* | -2.933 | -5.514 | -0.2198 | 11330 | 35 | 26 | 1993 | 2018 |
| *Systasea zampa* | -2.846 | -7.61 | 1.893 | 226 | 4 | 24 | 1993 | 2018 |
| *Lycaena hyllus* | -2.838 | -6.439 | 1.0136 | 3287 | 73 | 26 | 1993 | 2018 |
| *Anthocharis sara* | -2.769 | -7.044 | 1.5338 | 1572 | 17 | 26 | 1993 | 2018 |
| *Neominois ridingsii* | -2.762 | -11.26 | 6.467 | 1118 | 3 | 17 | 1994 | 2015 |
| *Erynnis propertius* | -2.7346 | -7.756 | 2.945 | 1696 | 19 | 26 | 1993 | 2018 |
| *Glaucopsyche lygdamus* | -2.722 | -7.938 | 2.6584 | 5709 | 45 | 26 | 1993 | 2018 |
| *Colias meadii* | -2.717 | -14 | 9.918 | 202 | 1 | 10 | 1994 | 2007 |
| *Calycopis isobeon* | -2.653 | -9.151 | 4.579 | 697 | 11 | 25 | 1993 | 2018 |
| *Speyeria edwardsii* | -2.639 | -7.963 | 2.96 | 313 | 4 | 21 | 1993 | 2014 |
| *Piruna polingi* | -2.4863 | -8.651 | 4.0932 | 1226 | 2 | 23 | 1993 | 2018 |
| *Oarisma garita* | -2.4417 | -6.92 | 3.748 | 3610 | 17 | 26 | 1993 | 2018 |
| *Speyeria aphrodite* | -2.434 | -7.816 | 3.06 | 27492 | 105 | 26 | 1993 | 2018 |
| *Pyrgus oileus* | -2.433 | -8.821 | 5.091 | 12868 | 62 | 26 | 1993 | 2018 |
| *Achalarus toxeus* | -2.418 | -31.03 | 37.5 | 230 | 1 | 9 | 1999 | 2009 |
| *Amblyscirtes celia* | -2.3712 | -9.198 | 6.6669 | 539 | 10 | 22 | 1993 | 2017 |
| *Poanes yehl* | -2.3569 | -9.522 | 7.5134 | 459 | 6 | 24 | 1995 | 2018 |
| *Carterocephalus palaemon* | -2.35204 | -8.018 | 3.8154 | 1266 | 15 | 25 | 1994 | 2018 |
| *Speyeria atlantis* | -2.313 | -8.825 | 4.564 | 17476 | 67 | 26 | 1993 | 2018 |
| *Callophrys johnsoni* | -2.256 | -12.85 | 9.257 | 60 | 2 | 12 | 2002 | 2018 |
| *Amblyscirtes cassus* | -2.25 | -8.175 | 3.422 | 565 | 5 | 26 | 1993 | 2018 |
| *Vanessa virginiensis* | -2.172 | -5.606 | 1.3719 | 29016 | 310 | 26 | 1993 | 2018 |
| *Polygonia gracilis* | -2.109 | -5.221 | 1.3109 | 2209 | 21 | 26 | 1993 | 2018 |
| *Oarisma poweshiek* | -2.0886 | -15.21 | 10.3 | 182 | 3 | 17 | 1996 | 2012 |
| *Hesperopsis alpheus* | -2.087 | -7.175 | 3.306 | 82 | 1 | 21 | 1995 | 2018 |
| *Oeneis chryxus* | -2.071 | -8.388 | 3.8025 | 2660 | 10 | 26 | 1993 | 2018 |
| *Colias eurytheme* | -2.0527 | -7.1491 | 3.472 | 267483 | 388 | 26 | 1993 | 2018 |
| *Neonympha areolata* | -2.0522 | -6.534 | 3.8016 | 2180 | 14 | 26 | 1993 | 2018 |
| *Nastra lherminier* | -2.0512 | -7.22 | 3.94 | 3406 | 56 | 26 | 1993 | 2018 |
| *Lycaeides melissa* | -2.0336 | -8.5035 | 4.679 | 16454 | 46 | 26 | 1993 | 2018 |
| *Pyrgus ruralis* | -2.0193 | -8.759 | 6.094 | 218 | 5 | 24 | 1993 | 2018 |
| *Panoquina panoquin* | -2.01 | -7.65 | 2.334 | 15480 | 13 | 26 | 1993 | 2018 |
| *Parnassius phoebus* | -2.0085 | -9.534 | 6.123 | 5701 | 10 | 26 | 1993 | 2018 |
| *Eunica monima* | -1.993 | -19.67 | 19.45 | 372 | 1 | 14 | 1997 | 2016 |
| *Callophrys niphon* | -1.979 | -13.06 | 12.381 | 1058 | 3 | 15 | 1995 | 2014 |
| *Callophrys sheridanii* | -1.949 | -7.725 | 3.772 | 247 | 6 | 23 | 1995 | 2018 |
| *Speyeria callippe* | -1.915 | -8.279 | 5.5017 | 9802 | 30 | 26 | 1993 | 2018 |
| *Vanessa cardui* | -1.8651 | -7.529 | 4.129 | 39197 | 249 | 26 | 1993 | 2018 |
| *Colias interior* | -1.8621 | -7.465 | 4.557 | 11749 | 27 | 26 | 1993 | 2018 |
| *Copaeodes aurantiacus* | -1.834 | -5.261 | 2.79921 | 2359 | 17 | 26 | 1993 | 2018 |
| *Atrytone arogos* | -1.8314 | -13.437 | 7.284 | 2751 | 7 | 24 | 1993 | 2018 |
| *Pieris napi* | -1.82768 | -5.792 | 2.76 | 14098 | 62 | 26 | 1993 | 2018 |
| *Mestra amymone* | -1.8223 | -9.061 | 6.218 | 293 | 4 | 18 | 1993 | 2017 |
| *Pyrgus philetas* | -1.8133 | -9.719 | 5.677 | 992 | 5 | 25 | 1993 | 2018 |
| *Lycaena helloides* | -1.8018 | -6.07 | 3.1638 | 5480 | 38 | 26 | 1993 | 2018 |
| *Amblyscirtes aenus* | -1.765 | -6.961 | 4.4346 | 1404 | 7 | 26 | 1993 | 2018 |
| *Colias philodice* | -1.75 | -6.598 | 3.232 | 228386 | 321 | 26 | 1993 | 2018 |
| *Nastra neamathla* | -1.7452 | -8.091 | 4.832 | 62 | 3 | 21 | 1996 | 2018 |
| *Cercyonis pegala* | -1.7409 | -6.889 | 3.764 | 272727 | 305 | 26 | 1993 | 2018 |
| *Appias drusilla* | -1.722 | -11.858 | 9.766 | 917 | 8 | 22 | 1995 | 2018 |
| *Euphyes dukesi* | -1.713 | -7.432 | 5.501 | 335 | 6 | 22 | 1994 | 2018 |
| *Pontia occidentalis* | -1.709 | -7.793 | 6.124 | 4570 | 27 | 26 | 1993 | 2018 |
| *Boloria epithore* | -1.709 | -6.646 | 3.549 | 6112 | 11 | 26 | 1993 | 2018 |
| *Cyllopsis pertepida* | -1.671 | -7.825 | 5.476 | 98 | 4 | 23 | 1993 | 2018 |
| *Polygonia satyrus* | -1.6483 | -4.693 | 1.556 | 2101 | 30 | 26 | 1993 | 2018 |
| *Polygonia interrogationis* | -1.6429 | -3.938 | 0.7742 | 48041 | 283 | 26 | 1993 | 2018 |
| *Plebejus icarioides* | -1.637 | -8.7228 | 5.915 | 22934 | 35 | 26 | 1993 | 2018 |
| *Callophrys dumetorum* | -1.6006 | -6.732 | 4.209 | 443 | 5 | 23 | 1993 | 2015 |
| *Papilio ornythion* | -1.583 | -18.46 | 18.72 | 9 | 1 | 6 | 2002 | 2016 |
| *Papilio canadensis* | -1.5824 | -6.58 | 4.1393 | 11022 | 57 | 26 | 1993 | 2018 |
| *Polites origenes* | -1.563 | -6.3077 | 3.713 | 8758 | 123 | 26 | 1993 | 2018 |
| *Limenitis weidemeyerii* | -1.5469 | -6.741 | 3.005 | 1599 | 12 | 26 | 1993 | 2018 |
| *Lycaena phlaeas* | -1.53059 | -7.3979 | 4.141 | 48049 | 138 | 26 | 1993 | 2018 |
| *Satyrodes eurydice* | -1.4872 | -6.1502 | 3.468 | 41732 | 120 | 26 | 1993 | 2018 |
| *Cyllopsis gemma* | -1.4797 | -6.559 | 3.577 | 2024 | 51 | 26 | 1993 | 2018 |
| *Satyrium saepium* | -1.462 | -10.326 | 8.172 | 14516 | 23 | 26 | 1993 | 2018 |
| *Cogia hippalus* | -1.461 | -6.897 | 4.176 | 4926 | 6 | 26 | 1993 | 2018 |
| *Plebejus acmon* | -1.4569 | -5.8859 | 3.309 | 20127 | 54 | 26 | 1993 | 2018 |
| *Boloria selene* | -1.4483 | -5.131 | 2.825 | 5371 | 68 | 26 | 1993 | 2018 |
| *Poanes hobomok* | -1.442 | -5.645 | 2.761 | 9811 | 104 | 26 | 1993 | 2018 |
| *Emesis ares* | -1.438 | -12.871 | 4.018 | 391 | 3 | 21 | 1993 | 2018 |
| *Phocides pigmalion* | -1.433 | -7.033 | 3.864 | 785 | 11 | 26 | 1993 | 2018 |
| *Callophrys spinetorum* | -1.4327 | -7.576 | 3.487 | 392 | 12 | 26 | 1993 | 2018 |
| *Colias pelidne* | -1.43266 | -15.23 | 13.98 | 82 | 2 | 12 | 1994 | 2014 |
| *Euphyes conspicua* | -1.431 | -6.206 | 3.2115 | 10126 | 53 | 26 | 1993 | 2018 |
| *Codatractus mysie* | -1.388 | -18.22 | 18.78 | 33 | 1 | 6 | 1998 | 2013 |
| *Heliopetes ericetorum* | -1.3848 | -6.296 | 4.048 | 2147 | 10 | 26 | 1993 | 2018 |
| *Phyciodes tulcis* | -1.3449 | -16.58 | 18.14 | 100 | 2 | 10 | 2000 | 2016 |
| *Lerodea eufala* | -1.3083 | -4.813 | 2.721 | 3141 | 74 | 26 | 1993 | 2018 |
| *Coenonympha tullia* | -1.3022 | -7.0237 | 4.583 | 61777 | 128 | 26 | 1993 | 2018 |
| *Apodemia nais* | -1.2924 | -9.062 | 7.84 | 858 | 4 | 25 | 1993 | 2018 |
| *Lycaena nivalis* | -1.28226 | -6.388 | 3.242 | 3118 | 9 | 26 | 1993 | 2018 |
| *Limenitis archippus* | -1.2744 | -4.6521 | 2.346 | 31806 | 298 | 26 | 1993 | 2018 |
| *Euphydryas gillettii* | -1.272 | -8.37 | 6.218 | 110 | 1 | 18 | 2000 | 2018 |
| *Satyrium acadica* | -1.2692 | -6.0704 | 4.813 | 7777 | 60 | 26 | 1993 | 2018 |
| *Battus polydamas* | -1.2625 | -5.373 | 4.775 | 2507 | 18 | 24 | 1995 | 2018 |
| *Euphilotes enoptes* | -1.2534 | -6.258 | 4.26147 | 10923 | 21 | 26 | 1993 | 2018 |
| *Erynnis tristis* | -1.2518 | -5.184 | 2.38708 | 3206 | 30 | 26 | 1993 | 2018 |
| *Hesperia nevada* | -1.2365 | -9.849 | 8.495 | 170 | 3 | 17 | 1994 | 2015 |
| *Pieris rapae* | -1.2101 | -6.1847 | 3.783 | 905161 | 379 | 26 | 1993 | 2018 |
| *Cercyonis sthenele* | -1.2 | -9.744 | 8.4416 | 3646 | 18 | 26 | 1993 | 2018 |
| *Satyrium tetra* | -1.172 | -8.038 | 11.851 | 5496 | 9 | 26 | 1993 | 2018 |
| *Polites mystic* | -1.1716 | -5.72847 | 3.093 | 22074 | 108 | 26 | 1993 | 2018 |
| *Parnassius clodius* | -1.1562 | -7.527 | 3.73636 | 8070 | 12 | 26 | 1993 | 2018 |
| *Vanessa atalanta* | -1.1027 | -4.869 | 2.728 | 82633 | 376 | 26 | 1993 | 2018 |
| *Thorybes pylades* | -1.0661 | -3.865 | 2.2731 | 21454 | 150 | 26 | 1993 | 2018 |
| *Thorybes bathyllus* | -1.04 | -5.8512 | 3.904 | 6739 | 97 | 26 | 1993 | 2018 |
| *Megisto cymela* | -1.0122 | -5.787 | 3.852 | 111127 | 232 | 26 | 1993 | 2018 |
| *Asterocampa celtis* | -1.012 | -6.873 | 4.947 | 157448 | 189 | 26 | 1993 | 2018 |
| *Danaus gilippus* | -0.9926 | -6.781 | 6.327 | 31203 | 67 | 26 | 1993 | 2018 |
| *Achalarus lyciades* | -0.967 | -5.8837 | 4.171 | 4326 | 70 | 26 | 1993 | 2018 |
| *Papilio palamedes* | -0.9578 | -5.9109 | 4.0358 | 34518 | 63 | 26 | 1993 | 2018 |
| *Papilio zelicaon* | -0.9289 | -7.08225 | 4.7805 | 4683 | 38 | 26 | 1993 | 2018 |
| *Papilio polyxenes* | -0.915 | -4.423 | 2.946 | 41379 | 332 | 26 | 1993 | 2018 |
| *Erynnis funeralis* | -0.912778 | -4.445 | 3.3807 | 2583 | 39 | 26 | 1993 | 2018 |
| *Eurema proterpia* | -0.9027 | -9.405 | 9.096 | 4337 | 5 | 21 | 1993 | 2018 |
| *Satyrium behrii* | -0.8921 | -5.65 | 4.875 | 5318 | 12 | 26 | 1993 | 2018 |
| *Dymasia dymas* | -0.8758 | -9.48 | 8.34552 | 9382 | 6 | 26 | 1993 | 2018 |
| *Pholisora catullus* | -0.8228 | -4.414 | 3.198 | 13519 | 136 | 26 | 1993 | 2018 |
| *Oeneis nevadensis* | -0.8199 | -6.069 | 5.079 | 2408 | 5 | 20 | 1994 | 2018 |
| *Erynnis pacuvius* | -0.80649 | -5.606 | 4.385 | 1628 | 18 | 26 | 1993 | 2018 |
| *Libytheana carinenta* | -0.8 | -5.707 | 4.608 | 69515 | 140 | 26 | 1993 | 2018 |
| *Parrhasius m-album* | -0.7931 | -8.29 | 6.9223 | 355 | 15 | 20 | 1997 | 2018 |
| *Calycopis cecrops* | -0.7624 | -5.865 | 4.744 | 12882 | 119 | 26 | 1993 | 2018 |
| *Papilio cresphontes* | -0.7383 | -3.9088 | 2.7345 | 12636 | 143 | 26 | 1993 | 2018 |
| *Pontia protodice* | -0.7379 | -7.461 | 6.356 | 28762 | 114 | 26 | 1993 | 2018 |
| *Limenitis arthemis* | -0.7015 | -4.2007 | 2.888 | 106071 | 343 | 26 | 1993 | 2018 |
| *Phyciodes campestris* | -0.6846 | -5.659 | 4.4851 | 15303 | 38 | 26 | 1993 | 2018 |
| *Chlosyne gabbii* | -0.6787 | -6.471 | 7.481 | 403 | 7 | 26 | 1993 | 2018 |
| *Enodia anthedon* | -0.67313 | -3.316 | 2.1567 | 21878 | 193 | 26 | 1993 | 2018 |
| *Euphilotes battoides* | -0.6713 | -5.331 | 4.5507 | 10627 | 20 | 26 | 1993 | 2018 |
| *Polites draco* | -0.6702 | -5.944 | 5.579 | 228 | 6 | 21 | 1993 | 2015 |
| *Neophasia menapia* | -0.6596 | -12.891 | 14.06 | 94 | 2 | 12 | 2001 | 2016 |
| *Speyeria cybele* | -0.6456 | -5.215 | 4.258 | 183124 | 276 | 26 | 1993 | 2018 |
| *Amblyscirtes vialis* | -0.6365 | -5.45 | 4.845 | 1479 | 37 | 26 | 1993 | 2018 |
| *Polygonia oreas* | -0.634057 | -8.776 | 6.798 | 130 | 4 | 25 | 1993 | 2018 |
| *Hesperia sassacus* | -0.6267 | -5.442 | 4.237 | 314 | 11 | 24 | 1993 | 2018 |
| *Electrostrymon angelia* | -0.6235 | -7.042 | 7.901 | 654 | 8 | 22 | 1995 | 2018 |
| *Atalopedes campestris* | -0.6026 | -6.621 | 5.602 | 107153 | 158 | 26 | 1993 | 2018 |
| *Amblyscirtes nereus* | -0.6026 | -5.054 | 3.3587 | 190 | 3 | 23 | 1993 | 2018 |
| *Satyrium titus* | -0.5831 | -4.1666 | 3.394 | 12255 | 139 | 26 | 1993 | 2018 |
| *Polites vibex* | -0.57671 | -4.859 | 3.853 | 13848 | 68 | 26 | 1993 | 2018 |
| *Leptotes marina* | -0.5583 | -7.604 | 6.879 | 23398 | 28 | 26 | 1993 | 2018 |
| *Satyrium liparops* | -0.54947 | -3.726 | 2.8998 | 3227 | 95 | 26 | 1993 | 2018 |
| *Celastrina neglectamajor* | -0.5478 | -12.847 | 9.377 | 545 | 5 | 21 | 1994 | 2017 |
| *Urbanus dorantes* | -0.52559 | -5.202 | 4.828 | 1444 | 28 | 26 | 1993 | 2018 |
| *Papilio multicaudata* | -0.5231 | -3.629 | 3.3457 | 6271 | 32 | 26 | 1993 | 2018 |
| *Thorybes confusis* | -0.5163 | -4.75 | 4.517 | 454 | 18 | 26 | 1993 | 2018 |
| *Danaus eresimus* | -0.5049 | -7.84 | 8.456 | 2801 | 15 | 25 | 1994 | 2018 |
| *Phyciodes tharos* | -0.4975 | -5.997 | 5.572 | 327395 | 369 | 26 | 1993 | 2018 |
| *Caria ino* | -0.454 | -7.644 | 6.877 | 236 | 2 | 21 | 1993 | 2016 |
| *Plebejus lupini* | -0.4263 | -6.63 | 5.87846 | 1824 | 11 | 26 | 1993 | 2018 |
| *Anatrytone logan* | -0.4177 | -5.273 | 4.976 | 33109 | 209 | 26 | 1993 | 2018 |
| *Asterocampa clyton* | -0.4104 | -6.7651 | 6.332 | 115558 | 146 | 26 | 1993 | 2018 |
| *Pyrgus albescens* | -0.3966 | -6.346 | 5.288 | 1672 | 5 | 21 | 1993 | 2017 |
| *Phyciodes picta* | -0.3596 | -7.665 | 6.4809 | 433 | 5 | 23 | 1993 | 2018 |
| *Staphylus ceos* | -0.3253 | -4.99 | 6.688 | 3425 | 6 | 26 | 1993 | 2018 |
| *Satyrium edwardsii* | -0.291 | -6.18 | 7.061 | 17479 | 50 | 26 | 1993 | 2018 |
| *Hermeuptychia sosybius* | -0.2811 | -5.869 | 4.756 | 56471 | 111 | 26 | 1993 | 2018 |
| *Polites themistocles* | -0.2809 | -3.58848 | 3.354 | 17314 | 212 | 26 | 1993 | 2018 |
| *Phoebis agarithe* | -0.2737 | -5.72 | 5.4617 | 9301 | 28 | 26 | 1993 | 2018 |
| *Speyeria mormonia* | -0.2185 | -8.366 | 9.57 | 3349 | 12 | 26 | 1993 | 2018 |
| *Ochlodes sylvanoides* | -0.1927 | -8.564 | 7.6384 | 2771 | 21 | 26 | 1993 | 2018 |
| *Chlosyne cyneas* | -0.1876 | -11.377 | 16.072 | 42 | 2 | 12 | 1993 | 2017 |
| *Papilio rutulus* | -0.1786 | -3.30272 | 2.9902 | 17363 | 59 | 26 | 1993 | 2018 |
| *Euphyes bimacula* | -0.1674 | -7.95 | 8.184 | 847 | 10 | 26 | 1993 | 2018 |
| *Anartia jatrophae* | -0.1391 | -5.9 | 7.175 | 34460 | 38 | 26 | 1993 | 2018 |
| *Strymon melinus* | -0.1248 | -4.38394 | 4.161 | 37580 | 274 | 26 | 1993 | 2018 |
| *Nymphalis antiopa* | -0.116 | -2.9383 | 2.947 | 15538 | 239 | 26 | 1993 | 2018 |
| *Asbolis capucinus* | -0.09286 | -3.966 | 3.56 | 913 | 19 | 24 | 1995 | 2018 |
| *Urbanus proteus* | -0.07469 | -5.421 | 5.992 | 3209 | 53 | 26 | 1993 | 2018 |
| *Texola elada* | -0.05002 | -10.1333 | 10.612 | 11990 | 8 | 26 | 1993 | 2018 |
| *Amblyscirtes aesculapius* | -0.0465 | -5.058 | 5.997 | 858 | 22 | 25 | 1993 | 2018 |
| *Feniseca tarquinius* | -0.03147 | -3.913 | 4.129 | 729 | 40 | 26 | 1993 | 2018 |
| *Speyeria hydaspe* | -0.01534 | -6.682 | 6.866 | 4225 | 17 | 26 | 1993 | 2018 |
| *Colias cesonia* | 0.03871 | -6.779 | 7.565 | 8166 | 32 | 26 | 1993 | 2018 |
| *Poanes massasoit* | 0.04937 | -2.836 | 2.908 | 10412 | 52 | 26 | 1993 | 2018 |
| *Atlides halesus* | 0.08876 | -4.471 | 4.896 | 608 | 18 | 26 | 1993 | 2018 |
| *Paratrytone snowi* | 0.08951 | -6.245 | 8.025 | 96 | 3 | 17 | 1993 | 2018 |
| *Hesperia comma* | 0.1098 | -7.159 | 7.5902 | 6661 | 20 | 26 | 1993 | 2018 |
| *Papilio glaucus* | 0.1109 | -3.593 | 4.18351 | 123356 | 330 | 26 | 1993 | 2018 |
| *Lycaena gorgon* | 0.115 | -6.701 | 5.5537 | 8419 | 9 | 26 | 1993 | 2018 |
| *Colias alexandra* | 0.1461 | -6.391 | 8.022 | 4716 | 19 | 26 | 1993 | 2018 |
| *Euphyes arpa* | 0.15287 | -5.049 | 7.619 | 756 | 8 | 24 | 1995 | 2018 |
| *Lycaena epixanthe* | 0.2127 | -5.181 | 6.078 | 23435 | 27 | 26 | 1993 | 2018 |
| *Speyeria egleis* | 0.2159 | -8.976 | 11.8 | 4789 | 9 | 26 | 1993 | 2018 |
| *Euphydryas phaeton* | 0.2287 | -5.462 | 6.2662 | 30949 | 87 | 26 | 1993 | 2018 |
| *Ancyloxypha numitor* | 0.24007 | -4.67492 | 5.646 | 61865 | 280 | 26 | 1993 | 2018 |
| *Poanes viator* | 0.24316 | -6.5557 | 7.5987 | 36690 | 79 | 26 | 1993 | 2018 |
| *Satyrium fuliginosa* | 0.285 | -8.224 | 10.103 | 429 | 5 | 23 | 1994 | 2018 |
| *Anaea andria* | 0.29202 | -3.712 | 4.312 | 2530 | 43 | 26 | 1993 | 2018 |
| *Polygonia comma* | 0.2957 | -3.4061 | 4.478 | 16770 | 219 | 26 | 1993 | 2018 |
| *Callophrys gryneus* | 0.30137 | -7.237 | 8.194 | 16270 | 77 | 26 | 1993 | 2018 |
| *Boloria bellona* | 0.3686 | -4.643 | 5.7241 | 28625 | 133 | 26 | 1993 | 2018 |
| *Erynnis zarucco* | 0.3877 | -5.719 | 7.198 | 6106 | 44 | 26 | 1993 | 2018 |
| *Colias occidentalis* | 0.4052 | -3.157 | 4.047 | 3092 | 3 | 26 | 1993 | 2018 |
| *Wallengrenia egeremet* | 0.4198 | -4.28292 | 4.959 | 49782 | 200 | 26 | 1993 | 2018 |
| *Euphyes vestris* | 0.4718 | -3.8817 | 4.882 | 118246 | 315 | 26 | 1993 | 2018 |
| *Erynnis persius* | 0.475 | -3.906 | 4.994 | 1844 | 21 | 26 | 1993 | 2018 |
| *Wallengrenia otho* | 0.5002 | -3.659 | 4.8 | 4260 | 66 | 25 | 1994 | 2018 |
| *Euchloe hyantis* | 0.5146 | -5.768 | 7.237 | 107 | 1 | 15 | 2001 | 2017 |
| *Enodia portlandia* | 0.5212 | -4.62 | 5.67919 | 1293 | 25 | 26 | 1993 | 2018 |
| *Chlosyne gorgone* | 0.5231 | -5.973 | 7.47 | 2463 | 26 | 26 | 1993 | 2018 |
| *Pyrgus communis* | 0.54119 | -4.73011 | 6.151 | 21699 | 150 | 26 | 1993 | 2018 |
| *Hemiargus isola* | 0.54598 | -5.544 | 7.576 | 17205 | 44 | 26 | 1993 | 2018 |
| *Ochlodes agricola* | 0.5699 | -5.049 | 6.971 | 14630 | 22 | 26 | 1993 | 2018 |
| *Epargyreus clarus* | 0.5727 | -4.458 | 5.869 | 201337 | 336 | 26 | 1993 | 2018 |
| *Amblyscirtes nysa* | 0.6047 | -3.4126 | 5.013 | 676 | 7 | 26 | 1993 | 2018 |
| *Papilio eurymedon* | 0.6228 | -2.3447 | 3.79 | 14691 | 44 | 26 | 1993 | 2018 |
| *Limenitis lorquini* | 0.6902 | -2.5144 | 4.453 | 13225 | 40 | 26 | 1993 | 2018 |
| *Papilio troilus* | 0.7255 | -3.237 | 4.8455 | 60437 | 255 | 26 | 1993 | 2018 |
| *Danaus plexippus* | 0.7298 | -2.875 | 4.568 | 135705 | 403 | 26 | 1993 | 2018 |
| *Calephelis mutica* | 0.7314 | -11.223 | 14.642 | 286 | 3 | 21 | 1997 | 2018 |
| *Pontia beckerii* | 0.74586 | -4.753 | 8.189 | 3073 | 14 | 26 | 1993 | 2018 |
| *Poanes taxiles* | 0.7479 | -4.618 | 7.1788 | 3967 | 11 | 26 | 1993 | 2018 |
| *Polites peckius* | 0.7587 | -4.167 | 6.061 | 25926 | 167 | 26 | 1993 | 2018 |
| *Satyrium calanus* | 0.7835 | -4.453 | 6.291 | 25691 | 169 | 26 | 1993 | 2018 |
| *Polygonus leo* | 0.8383 | -6.465 | 9.344 | 860 | 8 | 24 | 1995 | 2018 |
| *Polygonia faunus* | 0.8431 | -3.911 | 6.191 | 481 | 19 | 26 | 1993 | 2018 |
| *Chlosyne lacinia* | 0.856 | -5.681 | 6.71 | 7929 | 14 | 26 | 1993 | 2018 |
| *Euptoieta claudia* | 0.8834 | -4.3754 | 6.693 | 43910 | 203 | 26 | 1993 | 2018 |
| *Hylephila phyleus* | 0.8907 | -3.936 | 6.02503 | 73077 | 197 | 26 | 1993 | 2018 |
| *Battus philenor* | 0.8961 | -4.437 | 6.699 | 50023 | 169 | 26 | 1993 | 2018 |
| *Hemiargus ceraunus* | 0.9133 | -6.773 | 9.829 | 29742 | 58 | 26 | 1993 | 2018 |
| *Speyeria zerene* | 0.9514 | -3.659 | 5.758 | 2840 | 16 | 26 | 1993 | 2018 |
| *Celastrina ladon* | 0.961 | -2.866 | 5.0348 | 110296 | 345 | 26 | 1993 | 2018 |
| *Staphylus hayhurstii* | 0.9715 | -3.397 | 5.541 | 795 | 24 | 25 | 1994 | 2018 |
| *Hesperia attalus* | 0.9995 | -17.52 | 20.22 | 117 | 2 | 13 | 1995 | 2015 |
| *Ochlodes yuma* | 1.001 | -7.363 | 9.777 | 157 | 2 | 20 | 1994 | 2018 |
| *Callophrys polios* | 1.0401 | -12.81 | 18.66 | 63 | 2 | 8 | 1999 | 2014 |
| *Calpodes ethlius* | 1.0869 | -3.43 | 5.978 | 1092 | 18 | 26 | 1993 | 2018 |
| *Satyrodes appalachia* | 1.09843 | -3.1179 | 5.6984 | 16510 | 129 | 26 | 1993 | 2018 |
| *Lycaena editha* | 1.113 | -5.8272 | 8.9504 | 1474 | 10 | 26 | 1993 | 2018 |
| *Phyciodes mylitta* | 1.1661 | -2.10801 | 4.704 | 11932 | 44 | 26 | 1993 | 2018 |
| *Pompeius verna* | 1.1927 | -4.501 | 7.371 | 46607 | 192 | 26 | 1993 | 2018 |
| *Coenonympha haydenii* | 1.2105 | -5.874 | 9.074 | 953 | 2 | 23 | 1993 | 2018 |
| *Phoebis philea* | 1.2121 | -4.4537 | 6.571 | 2588 | 19 | 26 | 1993 | 2018 |
| *Chlosyne nycteis* | 1.241 | -5.292 | 8.447 | 36199 | 128 | 26 | 1993 | 2018 |
| *Eurytides marcellus* | 1.2454 | -4.06 | 6.6081 | 21895 | 81 | 26 | 1993 | 2018 |
| *Brephidium exile* | 1.26 | -7.158 | 11.258 | 21440 | 21 | 26 | 1993 | 2018 |
| *Cercyonis oetus* | 1.2671 | -6.868 | 8.966 | 9972 | 20 | 26 | 1993 | 2018 |
| *Polites sabuleti* | 1.2732 | -7.594 | 9.11 | 1830 | 10 | 26 | 1993 | 2018 |
| *Lerema accius* | 1.324 | -5.917 | 9.054 | 11973 | 112 | 26 | 1993 | 2018 |
| *Lycaena heteronea* | 1.3256 | -6.4185 | 9.0777 | 3814 | 15 | 26 | 1993 | 2018 |
| *Erynnis juvenalis* | 1.3581 | -3.995 | 7.414 | 1049 | 20 | 25 | 1993 | 2018 |
| *Erynnis lucilius* | 1.3837 | -7.662 | 12.113 | 1988 | 7 | 21 | 1994 | 2015 |
| *Chlosyne palla* | 1.417 | -5.971 | 9.819 | 8329 | 19 | 26 | 1993 | 2018 |
| *Phoebis sennae* | 1.422 | -5.152 | 8.508 | 73455 | 186 | 26 | 1993 | 2018 |
| *Eurema nicippe* | 1.4388 | -5.198 | 8.677 | 53577 | 149 | 26 | 1993 | 2018 |
| *Erynnis horatius* | 1.4684 | -4.2583 | 7.318 | 75707 | 187 | 26 | 1993 | 2018 |
| *Panoquina ocola* | 1.483 | -4.302 | 7.5881 | 6495 | 56 | 26 | 1993 | 2018 |
| *Nathalis iole* | 1.557 | -7.039 | 9.976 | 75994 | 93 | 26 | 1993 | 2018 |
| *Poanes melane* | 1.5637 | -2.096 | 5.95 | 5465 | 18 | 26 | 1993 | 2018 |
| *Megathymus cofaqui* | 1.5803 | -11.468 | 19 | 27 | 2 | 12 | 2003 | 2018 |
| *Eurema daira* | 1.599 | -6.321 | 9.614 | 13929 | 43 | 26 | 1993 | 2018 |
| *Polygonia progne* | 1.623 | -0.8532 | 4.445 | 2581 | 52 | 26 | 1993 | 2018 |
| *Euphyes dion* | 1.7188 | -3.262 | 7.017 | 4188 | 53 | 26 | 1993 | 2018 |
| *Calephelis nemesis* | 1.7277 | -3.974 | 7.711 | 1326 | 12 | 26 | 1993 | 2018 |
| *Boloria kriemhild* | 1.728 | -9 | 14.728 | 87 | 2 | 15 | 1993 | 2018 |
| *Erynnis baptisiae* | 1.7605 | -3.356 | 7.152 | 24378 | 135 | 26 | 1993 | 2018 |
| *Problema bulenta* | 1.799 | -4.218 | 9.384 | 968 | 5 | 26 | 1993 | 2018 |
| *Erebia discoidalis* | 1.821 | -20.39 | 30.31 | 61 | 1 | 5 | 2000 | 2014 |
| *Enodia creola* | 1.833 | -3.316 | 7.978 | 624 | 15 | 26 | 1993 | 2018 |
| *Hesperia columbia* | 2.01 | -49.43 | 105.6 | 10 | 1 | 6 | 2001 | 2017 |
| *Hesperia juba* | 2.154 | -2.997 | 7.982 | 417 | 12 | 26 | 1993 | 2018 |
| *Eurema dina* | 2.213 | -5.584 | 10.183 | 1065 | 2 | 21 | 1995 | 2018 |
| *Everes comyntas* | 2.2498 | -2.801 | 7.705 | 187988 | 298 | 26 | 1993 | 2018 |
| *Lycaena mariposa* | 2.272 | -2.333 | 8.271 | 500 | 5 | 25 | 1993 | 2018 |
| *Emesis zela* | 2.302 | -2.267 | 7.129 | 660 | 3 | 26 | 1993 | 2018 |
| *Phyciodes phaon* | 2.397 | -6.622 | 10.272 | 35758 | 77 | 26 | 1993 | 2018 |
| *Oligoria maculata* | 2.41959 | -4.2229 | 11.43 | 1170 | 20 | 25 | 1993 | 2018 |
| *Junonia coenia* | 2.4776 | -2.2543 | 7.389 | 96821 | 296 | 26 | 1993 | 2018 |
| *Nymphalis vaualbum* | 2.613 | -1.883 | 7.131 | 3208 | 31 | 26 | 1993 | 2018 |
| *Melanis pixe* | 2.617 | -7.44 | 14.34 | 26 | 2 | 10 | 1993 | 2016 |
| *Pyrgus scriptura* | 2.7183 | -5.469 | 10.37 | 491 | 3 | 26 | 1993 | 2018 |
| *Papilio machaon* | 2.8559 | -10.165 | 19.018 | 70 | 3 | 17 | 1997 | 2017 |
| *Eurema lisa* | 2.9398 | -1.9796 | 8.208 | 38806 | 144 | 26 | 1993 | 2018 |
| *Agraulis vanillae* | 3.04 | -3.201 | 9.9663 | 53863 | 134 | 26 | 1993 | 2018 |
| *Strymon acis* | 3.208 | -9.848 | 18.511 | 151 | 2 | 16 | 1997 | 2013 |
| *Calephelis borealis* | 3.284 | -1.004 | 8.882 | 2082 | 7 | 26 | 1993 | 2018 |
| *Junonia evarete* | 3.2953 | -5.6998 | 15.48 | 6438 | 11 | 26 | 1993 | 2018 |
| *Strymon istapa* | 3.316 | -2.4423 | 8.813 | 976 | 14 | 23 | 1993 | 2018 |
| *Atrytonopsis hianna* | 3.322 | -5.793 | 14.64 | 67 | 2 | 17 | 1994 | 2018 |
| *Lycaeides idas* | 3.3286 | -5.868 | 13.204 | 8335 | 15 | 26 | 1993 | 2018 |
| *Eunica tatila* | 3.331 | -11.23 | 20.13 | 170 | 1 | 9 | 2004 | 2015 |
| *Heliopetes laviana* | 3.344 | -4.6256 | 11.8617 | 465 | 2 | 21 | 1993 | 2016 |
| *Amblyscirtes fimbriata* | 3.3546 | -3.547 | 10.944 | 503 | 2 | 25 | 1993 | 2018 |
| *Heliconius charithonia* | 3.3986 | -1.2847 | 9.25 | 26195 | 42 | 26 | 1993 | 2018 |
| *Euphydryas chalcedona* | 3.416 | -7.667 | 13.859 | 38960 | 33 | 26 | 1993 | 2018 |
| *Chlosyne leanira* | 3.449 | -5.879 | 15.338 | 641 | 6 | 22 | 1993 | 2018 |
| *Leptotes cassius* | 3.482 | -4.1554 | 12.756 | 26702 | 34 | 26 | 1993 | 2018 |
| *Problema byssus* | 3.687 | -4.989 | 12.802 | 1676 | 20 | 26 | 1993 | 2018 |
| *Dryas iulia* | 3.7379 | -3.989 | 10.014 | 3235 | 14 | 26 | 1993 | 2018 |
| *Doxocopa laure* | 3.746 | -15.5 | 27.53 | 11 | 1 | 5 | 2004 | 2016 |
| *Satyrium caryaevorum* | 3.7715 | -1.4571 | 9.177 | 2508 | 36 | 26 | 1993 | 2018 |
| *Amblyscirtes eos* | 3.8068 | -3.0425 | 11.295 | 359 | 5 | 25 | 1993 | 2018 |
| *Poanes zabulon* | 3.833 | -3.4239 | 11.579 | 12091 | 77 | 26 | 1993 | 2018 |
| *Colias palaeno* | 3.87 | -5.533 | 14.25 | 21 | 1 | 6 | 2001 | 2017 |
| *Panoquina panoquinoides* | 3.8756 | -11.614 | 20.317 | 141 | 3 | 18 | 2000 | 2018 |
| *Marpesia petreus* | 4.3594 | -2.5462 | 12.63 | 3682 | 12 | 25 | 1994 | 2018 |
| *Poanes aaroni* | 4.669 | -5.675 | 18.153 | 7412 | 10 | 26 | 1993 | 2018 |
| *Hesperia dacotae* | 4.929 | -22.13 | 41.35 | 7 | 1 | 6 | 1996 | 2015 |
| *Ascia monuste* | 5.3594 | -3.0232 | 13.477 | 29358 | 44 | 26 | 1993 | 2018 |
| *Ephyriades brunneus* | 5.4275 | -4.528 | 15.8 | 690 | 5 | 22 | 1997 | 2018 |
| *Eumaeus atala* | 5.43 | -1.7328 | 12.66 | 10637 | 6 | 24 | 1995 | 2018 |
| *Autochton cellus* | 5.544 | 0.02655 | 11.091 | 2051 | 7 | 26 | 1993 | 2018 |
| *Speyeria idalia* | 6.008 | -4.364 | 16.134 | 12760 | 17 | 26 | 1993 | 2018 |
| *Nastra julia* | 6.215 | -6.315 | 23.06 | 30 | 3 | 12 | 2000 | 2016 |
| *Hesperia ottoe* | 6.341 | -7.861 | 18.71 | 118 | 3 | 16 | 1993 | 2014 |
| *Heliopetes macaira* | 6.397 | -1.9017 | 14.96 | 122 | 3 | 16 | 1994 | 2016 |
| *Amblyscirtes texanae* | 6.44 | -1.1272 | 17.035 | 125 | 3 | 21 | 1995 | 2018 |
| *Erora quaderna* | 6.5 | -0.8965 | 15.73 | 257 | 3 | 24 | 1993 | 2018 |
| *Astraptes fulgerator* | 6.793 | -20.24 | 42.94 | 7 | 1 | 5 | 2004 | 2016 |
| *Euptoieta hegesia* | 7.058 | -4.91 | 18.31 | 115 | 3 | 17 | 1993 | 2016 |
| *Satyrium kingi* | 7.085 | -1.0317 | 16.45 | 73 | 2 | 12 | 1996 | 2017 |
| *Siproeta stelenes* | 7.545 | -1.7336 | 17.74 | 97 | 4 | 21 | 1995 | 2017 |
| *Oeneis uhleri* | 8.105 | -3.9373 | 21.02 | 901 | 7 | 18 | 1995 | 2014 |
| *Polites baracoa* | 8.311 | 4.1251 | 15.39 | 1174 | 3 | 24 | 1995 | 2018 |
| *Cymaenes tripunctus* | 8.452 | -2.4812 | 20.426 | 563 | 13 | 21 | 1998 | 2018 |
| *Myscelia ethusa* | 8.574 | -2.435 | 22.02 | 201 | 2 | 20 | 1994 | 2016 |
| *Nymphalis californica* | 8.684 | -5.4045 | 24.859 | 33685 | 21 | 26 | 1993 | 2018 |
| *Euphyes berryi* | 8.707 | -5.317 | 33.17 | 164 | 4 | 15 | 1998 | 2015 |
| *Colias christina* | 8.749 | -32.17 | 74.49 | 21 | 1 | 6 | 2004 | 2011 |
| *Brephidium isophthalma* | 8.895 | -6.6693 | 33.526 | 31315 | 10 | 25 | 1993 | 2018 |
| *Timochares ruptifasciatus* | 9.357 | -6.398 | 27.27 | 26 | 1 | 8 | 2006 | 2016 |
| *Ministrymon clytie* | 9.475 | -3.35527 | 23.36 | 200 | 2 | 15 | 2000 | 2016 |
| *Staphylus mazans* | 9.864 | -7.679 | 30.48 | 92 | 2 | 9 | 1994 | 2010 |
| *Satyrium favonius* | 10.542 | -2.758 | 25.86 | 24 | 2 | 10 | 1993 | 2013 |
| *Phaeostrymon alcestis* | 10.593 | -10.877 | 35.94 | 535 | 2 | 11 | 2002 | 2015 |
| *Phoebis statira* | 11.018 | -0.5664 | 16.923 | 2395 | 5 | 22 | 1997 | 2018 |
| *Strymon martialis* | 11.087 | 1.7048 | 22.05 | 137 | 3 | 18 | 1998 | 2018 |
| *Cymaenes odilia* | 11.34 | 1.1609 | 22.2 | 53 | 2 | 13 | 2000 | 2016 |
| *Oeneis taygete* | 11.34 | -21.51 | 60.63 | 36 | 1 | 7 | 1993 | 2005 |
| *Plebejus shasta* | 11.47 | -1.067 | 25.8 | 465 | 1 | 16 | 1993 | 2010 |
| *Anaea aidea* | 15.61 | 6.03 | 25.5 | 117 | 2 | 16 | 1995 | 2016 |
| *Amblyscirtes elissa* | 15.65 | 5.383 | 23.95 | 514 | 3 | 24 | 1993 | 2018 |
| *Urbanus procne* | 18.19 | 7.648 | 30.47 | 786 | 3 | 18 | 1993 | 2016 |
| *Chlosyne janais* | 18.98 | 3.859 | 37.47 | 39 | 1 | 5 | 2007 | 2016 |
| *Amblyscirtes tolteca* | 19.88 | -7.631 | 56.91 | 94 | 2 | 10 | 2004 | 2018 |
| *Piruna cingo* | 21.38 | 8.781 | 36.87 | 2747 | 3 | 22 | 1993 | 2018 |
| *Megathymus streckeri* | 21.61 | -2.382 | 51.39 | 12 | 1 | 5 | 1993 | 2003 |
| *Euchloe olympia* | 23.16 | -11.13 | 71.92 | 52 | 1 | 6 | 1995 | 2004 |
| *Phyciodes orseis* | 33.34 | -4.057 | 80.57 | 56 | 1 | 6 | 2011 | 2018 |
| *Anartia fatima* | 35.06 | 13.12 | 60.14 | 24 | 1 | 5 | 2004 | 2016 |
| *Quasimellana eulogius* | 43.9 | 15.41 | 78.46 | 34 | 1 | 8 | 2001 | 2016 |
| *Phyciodes frisia* | 101 | 35.48 | 197.5 | 564 | 1 | 7 | 2003 | 2016 |
| *Electrostrymon mathewi* | 157.9 | 61.55 | 312.2 | 32 | 1 | 5 | 1996 | 2000 |
